# Thermodynamic evaluation of the impact of DNA mismatches in PCR-type SARS-CoV-2 primers and probes

**DOI:** 10.1101/2020.11.04.368449

**Authors:** Pâmella Miranda, Gerald Weber

## Abstract

**Background:** DNA mismatches can affect the efficiency of PCR techniques if the intended target has mismatches in primers or probes regions. The accepted rule is that mismatches are detrimental as they reduce the hybridization temperatures, yet a more quantitative assessment is rarely performed.

**Methods:** We calculate the hybridization temperatures of primer/probe sets after aligning to SARS-COV-2, SARS-COV-1 and non-SARS genomes, considering all possible combinations of single, double and triple consecutive mismatches. We consider the mismatched hybridization temperature within a range of 5 °C to the fully matched reference temperature.

**Results:** We obtained the alignments of 19 PCR primers sets that were recently reported for the detection of SARS-CoV-2 and to 21665 SARS-CoV-2 genomes as well as 323 genomes of other viruses of the coronavirus family of which 10 are SARS-CoV-1. We find that many incompletely aligned primers become fully aligned to most of the SARS-CoV-2 when mismatches are considered. However, we also found that many cross-align to SARS-CoV-1 and non-SARS genomes.

**Conclusions:** Some primer/probe sets only align substantially to most SARS-CoV-2 genomes if mismatches are taken into account. Unfortunately, by the same mechanism, almost 75% of these sets also align to some SARS-CoV-1 and non-SARS viruses. It is therefore recommended to consider mismatch hybridization for the design of primers whenever possible, especially to avoid undesired cross-reactivity.

## 1. Introduction

The coronavirus disease 2019 (COVID-19) pandemic has caused a flurry of activity regarding the detection of SARS-CoV-2, in particular a substantial amount of new RT-PCR primers were developed for this specific purpose [1–19]. A number of factors can influence the reliability of the PCR detection, such as sample contamination [20], cross-reactivity with other viruses [14], contamination of reagents [21], non-specific annealing [22] and poor amplification efficiency [23]. A crucial primer design factor is its hybridization melting temperature [24] that is related to the annealing of oligonucleotides. A set of primers with close melting temperatures and in the ideal range for primer extension usually ensures good PCR performance [25].

A factor that may interfere with the hybridization temperatures are the presence of mismatches, that is non-Watson-Crick base pairs, between the primer and the DNA target. This affects the stability of the duplex, usually leading to a decrease in the hybridization temperature [26, 27]. As a result, the presence of mismatches may influence the performance of primers restraining the amplification of DNA target. New mismatches arise due to mutations in primer regions of the target DNA, and may lead to false-negative results [20, 28–30]. This is of special concern for the case of RNA viruses that have a high mutation rates [19, 30]. Mutations that occur in the SARS-CoV-2 genome [31, 32] imply that the presence of mismatches between primer/probe and the template eventually become inevitable. On the other hand, it is known that mismatch presence may affect only the first few cycles of PCR [33, 34] and with proper design may even be advantageous [35]. Therefore, as a rule of thumb, the occurrence of single mismatches are admitted in the hope that they may not affect the detection of the target and its amplification [36]. Unfortunately, the thermodynamic instability caused by the presence of mismatches is rarely quantified in primer design for a number of reasons. One of which is that the prediction of hybridization temperatures involving mismatches carries large uncertainties. Unlike Watson-Crick complementary base pairs, AT and CG, the hydrogen bonding and stacking interactions of mismatches are strongly dependent on the adjacent base pairs. Temperature predictions rely on experimental melting temperature data which typically do not cover the full combinatorial spectrum of mismatches and were carried out under high sodium buffer conditions [37, 38]. However, this has now changed. A recent development from our group has reworked the parametrization for a comprehensive set of 4032 sequences containing up to three consecutive mismatches [39]. This now enables the analysis in unprecedented detail of the effect of mismatches in primer/probe hybridization.

Here, we analyse how and if mismatches do influence the melting temperatures of primer/probes hybridised to SARS-CoV-2 genomes. We collected 19 PCR primer/probe sets (297 primers and 43 probes) which cover seven different gene regions of SARS-CoV-2 genome (N, E, S, M, ORF1ab, RdRp and nsp2 genes) [1–19]. These primer/probes were aligned to 21665 genomes of SARS-CoV-2 and 323 genomes of other coronaviruses. Melting temperatures are calculated with a mesoscopic model using the newly developed parameters for up to three consecutive mismatches [39]. Using the mesoscopic model for the calculation of mismatches has an important advantage over nearest-neighbour models [37] as it naturally accounts for end effects, that is, mismatches located near the primer end may have different hybridization temperatures than those that are centred which reflects experimental observations on PCR efficiencies [34].

## 2. Materials and Methods

### 2.1. Genomes and primer sets

We collected *N_G_* = 21665 genomes of SARS-CoV-2 at NCBI [40], in 8 October 2020, and ensured that all were at least 25000 bp in size. The accession codes of these genomes are shown in supplementary table S1. To verify cross-reactivity we also performed the same analysis for *N*_h.c._ = 323 human coronaviruses (229E, NL63, OC43, HKU1, MERS), including SARS-CoV-1, and their accession codes are shown in supplementary table S2 and S3.

A total of 19 different primer/probe sets for RT-PCR were obtained from Refs. [1–19], their full details are shown in supplementary table S4. Note that several publications include primers from earlier reports. In particular, CDC primers [2] are included in several publications. Therefore, for each set we only considered those that were not repeated from other publications. Note that some primers and probes were designed for both SARS-CoV-1 and SARS-CoV-2 [4, 14].

### 2.2. Primer and genome alignment

Each primer or probe sequence is aligned against a given genome using a Smith-Waterman algorithm [41], where matching base pairs AT and CG where given score 2, mismatches score −1, and no gaps were considered. Alignments were carried out in two strand configurations, first for the genome sequences as obtained from the database and taking the primer/probe sequence as complementary strand

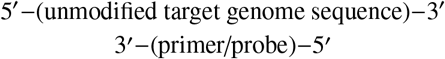

and next by taking the complementary of the genome sequence

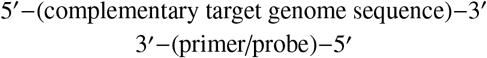

These alignments are carried out regardless if the primer was identified as forward or reverse. In all cases the nominal directions of the primers were identified correctly.

A primer/probe that was completely aligned to a target genome, without mismatches, was termed as strictly matched. If there were up to three contiguous mismatches in the alignment it was called as partially matched. The limit of three contiguous mismatches relates to the available melting temperature parameters. Alignments with four or more contiguous mismatches were considered as not aligned.

As an example of partial alignment, we show the RdRp_SARSr-R1 primer (bottom strand) in the MT457390 genome

5’-TATGCTAATAGTGT<u>T</u>TTTAACATTTG-3’
3’-ATACGATTATCACA<u>C</u>AAATTGTAAAC-5’

where the mismatched site is underlined.

### 2.3. Calculation of melting temperatures

Hybridization temperatures *T_m_* are calculated from

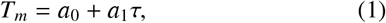

where *τ* is a statistical index calculated from the classical partition function of a model Hamiltonian, and *a*_0_ and *a*_1_ are regression coefficients obtained from a set of 4096 experimental melting temperatures of which 4032 are from sequences containing up to three consecutive mismatched base pairs [39]. The buffer conditions for these parameters are 50 mM sodium chloride, 10 mM sodium phosphate pH 7.4, and total strand concentration 1.0 *μ*M. For a complete description of the melting temperature calculation and experimental conditions see Ref. [39, 42]. The index *τ* was calculated for each primer/probe, using the parameters reported in Ref. [39], after aligning against a reference genome. The calculation of *τ* also yields the average displacement profile which shows the expected base-pair opening along the oligonucletide duplex, for details see Eq.(5) from Ref. 39.

### 2.4. Coverage evaluation

We calculated the hybridization temperatures from Eq. (1) for each primer/probe assuming a complete Watson-Crick complementarity which we called the reference temperature *T*_ref._, which are shown in supplementary table S5.

All 19 primers/probes were aligned against *N_G_* genomes and we kept only those alignments with up to three consecutive mismatches. The coverage for a strictly non-mismatched alignment *C*_strict_ was calculated as

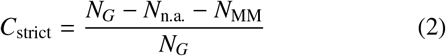

where *N_G_* the total number of genomes which are at least 25000 bp in size, *N*_n.a._ number of genomes for which no alignment was found, and *N*_MM_ the number of genomes for which a partial alignment with up to three consecutive mismatches was found.

Next, for each of the *N*_MM_ partial alignments we calculated the hybridization temperatures *T*_MM_ from Eq. (1) taking into account the mismatches, and the difference to the reference temperature *T*_ref._ is

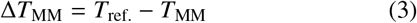

*T*_MM_ is usually, but not always, lower than *T*_ref._ [39]. We will consider the partially mismatched coverage *C*_part_. as

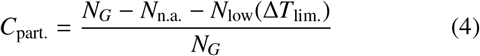

where *N*_low_ is the number of primers/probes satisfying

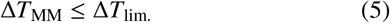

Here, we will use Δ*T*_lim._ = 5 °C, that is we will consider that mismatched primers/probes with *T*_MM_ no more than 5 °C below the reference temperature *T*_ref._ are still acceptable.

## 3. Results and Discussion

After aligning all primer/probe sets to all genomes we calculated their hybridization temperatures taking into account up to three consecutive mismatches, as detailed in the methods section. Table 1 summarises all sets analysed, their range of reference hybridization temperatures, strict and partial coverage for SARS-CoV-2 and for non-SARS-CoV-2 viruses. The detailed results for each primer are shown in supplementary table S5 for SARS-CoV-2. If the PCR primers can in principle bind to non-SARS-CoV-2 targets then this set may not specific [22]. Considering this, we analysed the 19 primer/probe sets in relation to genomes of other coronaviruses (non-SARS) as well as SARS-CoV-1 to verify if there was some cross-reactivity. The detailed results are shown in supplementary tables S6 for SARS-COV-1, and S7 for non-SARS viruses.

**Table 1:** Summary of the results for all primer/probe sets. Shown are the number of primers/probes *N_pp_* for each set, the range of reference temperatures *T*_ref._, the range of strict and partially mismatched coverages, for SARS-CoV-2, SARS-CoV-1 and non-SARS genomes. Detailed results for each primer are shown in supplementary table S5, S6 and S7.

| Name of Set | $N_{pp}$ | $T_{ref.}$ (°C) | SARS-CoV-2 | | SARS-CoV-1 | | non-SARS | |
| --- | --- | --- | --- | --- | --- | --- | --- | --- |
| | | | $C_{strict}$ (%) | $C_{part.}$ (%) | $C_{strict}$ (%) | $C_{part.}$ (%) | $C_{strict}$ (%) | $C_{part.}$ (%) |
| CDC [2] | 6 | 61.1–75.5 | 98.4–99.3 | 99.1–99.4 | 0.0–80.0 | 0.0–80.0 | 0.0 | 0.0 |
| WHO [1] | 21 | 51.3–70.3 | 97.9–99.4 | 97.9–99.5 | 0.0 | 0.0 | 0.0 | 0.0 |
| Luminex [3] | 6 | 59.4–81.0 | 64.9–99.5 | 65.1–99.5 | 0.0 | 0.0–100 | 0.0 | 0.0 |
| Bhadra et al. [8] | 6 | 61.3–64.2 | 98.7–99.4 | 98.7–99.4 | 0.0–100 | 0.0–100 | 0.0 | 0.0 |
| Corman et al. [4] | 21 | 61.7–81.7 | 0.0–99.4 | 0.0–99.4 | 0.0–100 | 0.0–100 | 0.0 | 0.0–19.2 |
| Davda et al. [5] | 16 | 56.5–70.3 | 95.3–99.4 | 95.3–99.4 | 0.0–80.0 | 0.0–90 | 0.0 | 0.0 |
| Grant et al. [6] | 2 | 62.4–79.9 | 97.5–99.3 | 97.5–99.5 | 0.0–100 | 0.0–100 | 0.0 | 0.0 |
| Hirotsu et al. [7] | 3 | 60.6–68.8 | 0.0–99.4 | 0.0–99.4 | 0.0 | 0.0 | 0.0 | 0.0 |
| Lanza et al. [9] | 27 | 60.5–75.0 | 98.5–99.4 | 98.8–99.5 | 0.0 | 0.0–90.0 | 0.0 | 0.0 |
| Li et al. [10] | 2 | 67.5–70.4 | 98.2–99.2 | 98.3–99.2 | 0.0 | 0.0–100 | 0.0 | 0.0 |
| Lu et al. [11] | 3 | 64.0–74.7 | 98.6–99.4 | 99.3–99.4 | 0.0 | 0.0 | 0.0 | 0.0 |
| Munnink et al. [12] | 171 | 65.4–74.9 | 45.9–99.5 | 46.0–99.7 | 0.0–100 | 0.0–100 | 0.0 | 0.0–0.639 |
| Nalla et al. [13] | 4 | 51.7–68.0 | 0.0–99.5 | 0.0–99.5 | 0.0–90.0 | 0.0–100 | 0.0–68.7 | 0.0–68.7 |
| Niu et al. [14] | 6 | 59.2–84.2 | 0.0–99.4 | 0.0–99.4 | 0.0–100 | 0.0–100 | 0.0 | 0.0 |
| Park et al. [15] | 20 | 59.3–65.4 | 94.9–99.5 | 94.9–99.5 | 0.0–100 | 0.0–100 | 0.0 | 0.0 |
| Rahman et al. [16] | 6 | 64.2–76.1 | 97.6–99.4 | 97.6–99.4 | 0.0 | 0.0–100 | 0.0 | 0.0 |
| Toptan et al. [17] | 6 | 62.2–65.4 | 98.8–99.5 | 98.8–99.5 | 0.0 | 0.0 | 0.0 | 0.0 |
| Vogels et al. [18] | 11 | 58.6–65.5 | 0.0–99.3 | 0.0–99.3 | 0.0–100 | 0.0–100 | 0.0 | 0.0 |
| Yip et al. [19] | 2 | 61.4–63.4 | 99.1–99.2 | 99.1–99.3 | 0.0 | 0.0 | 0.0 | 0.0 |

The typical design rules for PCR primers and probes recommend that the range of hybridization temperatures in a given set should be narrow, of the order of 10 °C [25, 43]. Several authors even suggest that the range for primer pair should be no more than 5°C [22, 44] or even less than 1°C [45]. However, it is evident, from table 1, that very few sets have temperature ranges below 10 °C, while some even exceed 20 °C. For example, the Luminex set, which includes the primer/probe set of China CDC, shows differences in the primer temperatures up to 21.6 °C. However, when mismatches are considered the hybridization temperatures may go far below the design range. In Fig. 1 we show an example of a displacement profile where a single AC mismatch completely disturbs the surrounding AT base pairs and the hybridization temperature drops to *T*_MM_ = 48.5 °C, down from a reference temperature of *T*_ref._ = 61.1 °. However, a presence of one or more mismatches does not necessarily imply in a reduction of hybridization temperature. For example, the SARS-CoV-2_89_RIGHT primer when aligned to MT259228.1 has two consecutive mismatches towards the 5’ end, see supplementary figure S1. Even though these mismatches induce a small end fraying, it has a calculated temperature of *T*_MM_ = 68.7 °C which is even somewhat higher than its reference temperature *T*_ref._ = 68.3 °C. This stability is caused by an increased stacking interaction between the GA and AA mismatches [39].

**Figure 1:**
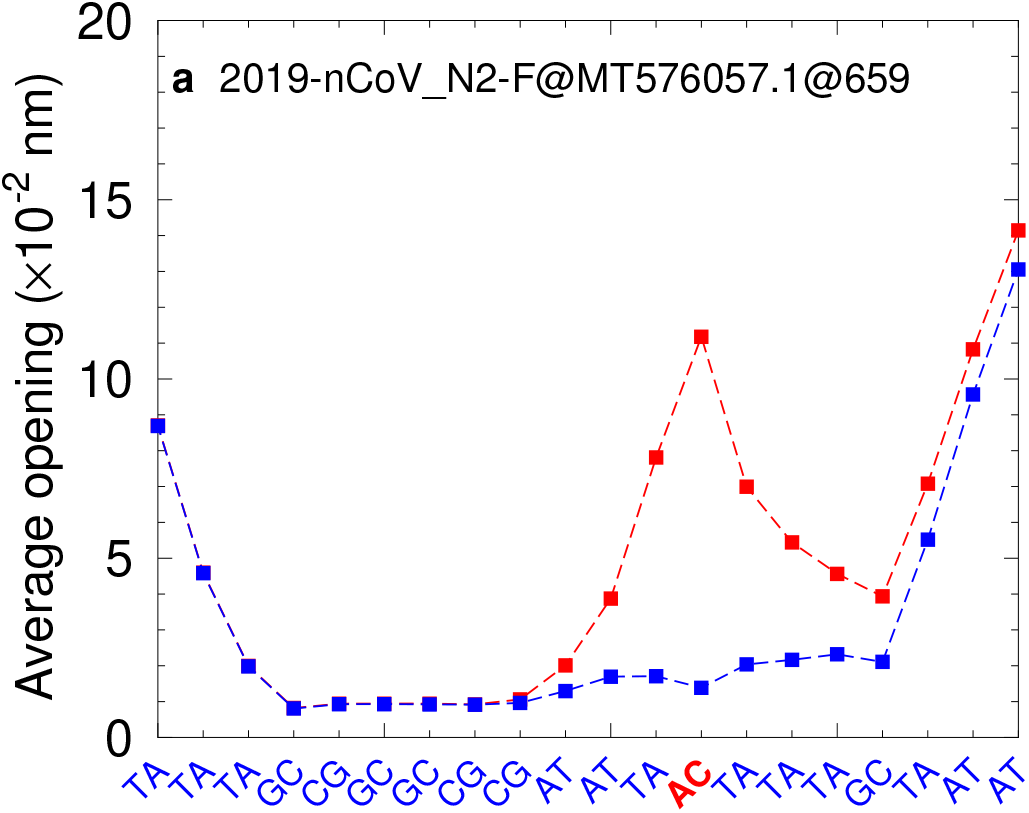
Displacement profile for CDC 2019-nCoV_N2-F when aligned to SARS-COV-2 MT576057.1 at position 659 has a mismatch AC (red symbols) instead of CG (blue symbols).

In terms of SARS-CoV-2 strict coverage, most sets have *C*_strict_ typically beyond 90%, which is expected as the primer design is guided by existing genomes. However, a number of specific probes, such as RdRp_SARSr-P1-1 from Corman et al. [4] go from 0 to 99.4% only if mismatches are taken into account. Indeed, as pointed out by Pillonel et al. [46] several of the probes from Corman et al. [4] do not fully match the available SARS-CoV-2 genomes (1623 at the time [46]). However, it was also observed that the mismatches had little effect on their efficiencies [13, 28] which is consistent with our calculations. The reason for the presence of mismatches in this case, as explained in Corman and Drosten [28], was the incomplete genomic information available at the time when this set was designed. It is worth noticing that while accounting for mismatches increases the coverage of this particular primer/probe set, it also increases the coverage for SARS-CoV-1 and even non-SARS as shown in Tab. 1.

We observed that in some cases the presence of few mismatches substantially decrease the hybridization temperature, leading to a complete absence of coverage. For example, four primers from Vogels et al. [18] do not align with any genome at all, not even when considering the mismatches as their hybridization temperatures *T*_MM_ are too low in comparison to reference temperature *T*_ref._. In contrast, for several cases when mismatches are taken into account the coverage becomes almost complete. A special example is probe 2019-nCoV_N1-P from CDC set that had 223 further mismatch alignments increasing the strict coverage of 98.4% to partial coverage of 99.4%. Similar findings were observed for SARS-CoV-2_6_LEFT [12] and NIID_WH-1_F501 of WHO [1].

The cross-reactivity, that is, the coverage of SARS-Cov-1 and non-SARS, appear in most primer/probe sets when mismatches are taken into account. Of the 19 primer/probe sets, we found only 5 sets that do not present cross-reactivity at all, see Tab. 1.

## 4. Conclusion

We evaluated the impact of mismatches in the hybridization of primers and probes for the detection of SARS-CoV-2 and other genomes. We have shown that the effect of mismatches on the probe/primer hybridization is not straightforward and can only be fully evaluated with a detailed calculation with up-to-date model parameters. In particular, our calculations showed that a substantial amount of the existing primers/probes may cross-react to SARS-CoV-1 and non-SARS genomes, which further highlights the need for taking mismatch hybridization into account.

## Supplementary data

Figure S1, Tables S1–S7.

