## Supplementary Material for "Thermodynamic evaluation of the impact of DNA mismatches in PCR-type SARS-CoV-2 primers and probes"

*November 4, 2020*

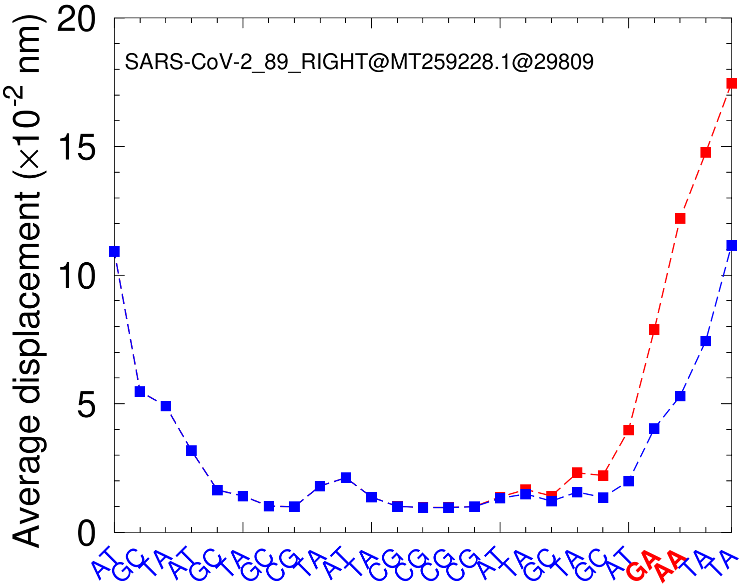

Figure S1: Displacement profile for SARS-CoV-2_89_RIGHT when aligned to SARS-COV-2 MT259228.1 at position 29809 has consecutive GA-AA mismatches (red symbols) instead of AT-AT base pairs (blue symbols).

Table S1: List of SARS-CoV-2 genome accession codes. Contiguous genome codes such as LR794478.1 to LR794483.1 are shown in compact notation LR7944(78–83).1

| | | | | |
|---|---|---|---|---|
| LC52823(2,3).1 | LC529905.1 | LC53441(8,9).1 | LC542809.1 | LC542976.1 |
| LC546038.1 | LC5475(18–33).1 | LC549340.1 | LC55325(7,8).1 | LC553259.2 |
| LC55326(0–3).1 | LC5532(69,70).1 | LC5563(15–24).1 | LC5654(07–14).1 | LC5678(45–59).1 |
| LC57096(1–3).1 | LC5709(67–99).1 | LC5710(02–13).1 | LC571019.1 | LC57102(2–5).1 |
| LC57103(0,1).1 | LC5710(34–41).1 | LC5720(67–70).1 | LC5732(73–86).1 | LC573289.1 |
| LR75799(5–8).1 | LR793257.1 | LR7943(80–91).1 | LR794(398–401).1 | LR7944(05–12).1 |
| LR794414.1 | LR79441(7,8).1 | LR79442(0–7).1 | LR79443(0–4).1 | LR794437.1 |
| LR79444(0–3).1 | LR794446.1 | LR79444(8,9).1 | LR7944(58–62).1 | LR79446(4–7).1 |
| LR7944(69–71).1 | LR794473.1 | LR7944(78–83).1 | LR79448(5–8).1 | LR79449(0–6).1 |
| LR794498.1 | LR7945(00–14).1 | LR79451(6–9).1 | LR79452(2,3).1 | LR7945(25–32).1 |
| LR7945(35–42).1 | LR79454(4–6).1 | LR7945(48–52).1 | LR79455(4–6).1 | LR7945(58–62).1 |
| LR79456(4–6).1 | LR7945(69–73).1 | LR794575.1 | LR79457(7,8).1 | LR79458(0–2).1 |
| LR794584.1 | LR79458(7,8).1 | LR79459(1–3).1 | LR794598.1 | LR79460(2,3).1 |
| LR79460(5–7).1 | LR7946(09–11).1 | LR794613.1 | LR7946(19–21).1 | LR7946(25–33).1 |
| LR79463(7–9).1 | LR79464(1–3).1 | LR7946(45–55).1 | LR7946(57–62).1 | LR79466(5–8).1 |
| LR79467(0,1).1 | LR794673.1 | LR7946(75–82).1 | LR794684.1 | LR81(3996–4000).1 |
| LR814001.2 | LR814002.4 | LR81400(3,4).3 | LR81400(5–7).2 | LR814008.3 |
| LR8140(09–13).2 | LR81402(0–3).2 | LR814024.3 | LR81402(5–9).2 | LR814030.3 |
| LR81403(1–4).2 | LR81403(5,6).3 | LR81403(7–9).2 | LR814040.3 | LR814041.4 |
| LR81404(2–6).2 | LR814047.4 | LR814048.3 | LR8141(09,10).1 | LR814111.2 |
| LR81411(2–5).1 | LR8141(18–23).1 | LR81412(6,7).1 | LR814128.4 | LR81413(0–3).1 |
| LR81413(5–8).1 | LR81414(1–3).1 | LR8141(46–50).1 | LR81415(2–6).1 | LR814157.3 |
| LR8141(58–62).1 | LR814163.3 | LR81416(4,5).1 | LR814166.3 | LR8141(67–80).1 |
| LR814183.1 | LR814184.2 | LR814185.1 | LR814186.2 | LR8141(89–92).1 |
| LR814193.2 | LR814(196–200).1 | LR814201.2 | LR814202.1 | LR814205.3 |

(continued on next page)

(Table S1 continued)

| | | | | |
|---|---|---|---|---|
| LR81420(6,7).2 | LR8142(08–11).1 | LR814214.1 | LR814215.3 | LR814216.2 |
| LR814219.2 | LR81422(0–5).1 | LR8142(28–33).1 | LR8142(36–40).1 | LR814243.3 |
| LR8142(44–63).1 | LR8142(66–72).1 | LR81556(3–8).1 | LR81557(4,5).3 | LR81557(6–9).1 |
| LR815581.1 | LR82182(0,1).1 | LR821822.3 | LR82182(3,4).1 | LR821826.3 |
| LR821827.1 | LR821828.3 | LR8218(29–41).1 | LR821842.3 | LR821843.1 |
| LR821844.2 | LR82184(5–9).1 | LR821850.2 | LR82185(1–3).1 | LR821854.2 |
| LR821855.1 | LR8218(57–64).1 | LR821865.4 | LR8218(66–74).1 | LR821875.3 |
| LR82187(6–8).1 | LR821879.4 | LR821880.1 | LR821881.4 | LR82188(2–7).1 |
| LR821888.3 | LR8218(89–95).1 | LR821896.4 | LR821897.1 | LR821898.2 |
| LR821(899–911).1 | LR821912.3 | LR8219(13–33).1 | LR82193(4,5).4 | LR821936.1 |
| LR821937.2 | LR8219(38–50).1 | LR821951.3 | LR82195(2–5).1 | LR821956.3 |
| LR82195(7–9).1 | LR821960.2 | LR82196(1,2).1 | LR821963.3 | LR8219(64–75).1 |
| LR821976.4 | LR82197(7,8).1 | LR821979.4 | LR821980.1 | LR821981.2 |
| LR821982.3 | LR82198(3–5).1 | LR8219(87–91).1 | LR821992.4 | LR821993.1 |
| LR821994.2 | LR82199(5–8).1 | LR821999.3 | LR82200(0–5).1 | LR824035.1 |
| LR824(037–130).1 | LR824(132–485).1 | LR82448(7,8).1 | LR824(490–531).1 | LR82456(3–7).1 |
| LR82457(0,1).1 | LR824(599,600).1 | LR860(559–655).1 | LR860(688–707).1 | LR8607(36–61).1 |
| LR86077(5–7).1 | LR860(792–811).1 | LR8608(40–66).1 | LR86089(5–9).1 | LR86091(3,4).1 |
| LR861(051–100).1 | LR86110(2–7).1 | LR8614(84–96).1 | LR8616(66–77).1 | LR86168(7–9).1 |
| LR862(399–407).1 | LR862(439–507).1 | LR87718(1–3).1 | LR877186.1 | LR877330.1 |
| LR877(382–431).1 | LR877(721–825).1 | LR877883.1 | LR87(7886–8363).1 | LR880(573–644).1 |
| LR881(869–933).1 | LR8820(37–49).1 | LR882(062–374).1 | LR882(380–492).1 | MN908947.3 |
| MN938384.1 | MN975262.1 | MN985325.1 | MN98866(8,9).1 | MN988713.1 |
| MN99446(7,8).1 | MN9965(27–31).1 | MN997409.1 | MT007544.1 | MT012098.1 |
| MT0195(29–33).1 | MT020781.2 | MT02088(0,1).1 | MT02706(2–4).1 | MT034054.1 |
| MT03987(3,4).1 | MT03988(7,8).1 | MT039890.1 | MT04425(7,8).1 | MT049951.1 |
| MT05049(1–3).1 | MT066156.1 | MT06617(5,6).1 | MT072688.1 | MT077125.1 |
| MT0798(43–54).1 | MT093571.1 | MT093631.2 | MT10605(2–4).1 | MT11441(2–9).1 |

(continued on next page)

(Table S1 continued)

| | | | | |
|---|---|---|---|---|
| MT118835.1 | MT121215.1 | MT123290.1 | MT12329(1–3).2 | MT126808.1 |
| MT13504(1–4).1 | MT152824.1 | MT1597(05–22).2 | MT1637(16–21).1 | MT18490(7–9).2 |
| MT18491(0–3).1 | MT186683.1 | MT1883(39–41).1 | MT192759.1 | MT192765.1 |
| MT19277(2,3).1 | MT198651.1 | MT198652.2 | MT198653.1 | MT21519(4,5).1 |
| MT226610.1 | MT230904.1 | MT2335(19–23).1 | MT233526.1 | MT240479.1 |
| MT2464(49–90).1 | MT246667.1 | MT2519(72–80).1 | MT2526(77–91).1 | MT252(693–710).1 |
| MT2527(12–33).1 | MT2527(35–50).1 | MT2527(52–64).1 | MT2527(67–70).1 | MT2527(72–89).1 |
| MT252(791–807).1 | MT2528(09–20).1 | MT25282(2,3).1 | MT253(696–710).1 | MT25691(7,8).1 |
| MT256924.2 | MT2583(77–83).1 | MT2592(26–31).1 | MT2592(35–87).1 | MT262(896–916).1 |
| MT262993.1 | MT263074.1 | MT263(381–465).1 | MT26346(7–9).1 | MT2701(01–14).1 |
| MT27081(4,5).1 | MT27632(3,4).1 | MT27632(5,6).2 | MT276327.1 | MT276328.2 |
| MT276329.1 | MT276330.2 | MT27659(7,8).1 | MT276600.1 | MT281577.1 |
| MT2918(26–34).1 | MT291835.2 | MT291836.1 | MT2925(69–75).1 | MT293156.1 |
| MT2931(58–92).1 | MT293(194–202).1 | MT2932(04–13).1 | MT29321(5,6).1 | MT2932(18–20).1 |
| MT293222.1 | MT29322(4,5).1 | MT29546(4,5).1 | MT300186.2 | MT3044(74–91).1 |
| MT30870(2–4).1 | MT318827.1 | MT320538.2 | MT320891.2 | MT32239(4–8).1 |
| MT3224(00–24).1 | MT324062.1 | MT324684.1 | MT325(561–640).1 | MT32602(3,4).1 |
| MT3260(27–30).1 | MT32603(5,6).1 | MT3260(39–42).1 | MT326046.1 | MT32604(8,9).1 |
| MT32605(1–6).1 | MT326058.1 | MT326061.1 | MT326063.1 | MT3260(65–71).1 |
| MT32607(4–6).1 | MT326078.1 | MT3260(81–97).1 | MT326(099,100).1 | MT32610(3–6).1 |
| MT32611(1–3).1 | MT3261(16–21).1 | MT32612(3–5).1 | MT3261(27–30).1 | MT32613(2–8).1 |
| MT326140.1 | MT3261(48–54).1 | MT3261(58–62).1 | MT326164.1 | MT32616(6,7).1 |
| MT326169.1 | MT326173.1 | MT326178.1 | MT32618(0,1).1 | MT32618(4,5).1 |
| MT326187.1 | MT32619(0,1).1 | MT327745.1 | MT32803(2–5).1 | MT334523.1 |
| MT334526.1 | MT334529.1 | MT334531.1 | MT3345(33–42).1 | MT334544.1 |
| MT33454(6,7).1 | MT334557.1 | MT334561.1 | MT3390(39–41).1 | MT339043.1 |
| MT3449(44–63).1 | MT34579(8,9).1 | MT34580(1–3).1 | MT34580(5,6).1 | MT34580(8,9).1 |
| MT34581(1,2).1 | MT34581(4–7).1 | MT345819.1 | MT3458(21–30).1 | MT34583(2–6).1 |

(continued on next page)

(Table S1 continued)

| | | | | |
|---|---|---|---|---|
| MT345838.1 | MT34584(0–4).1 | MT3458(46–62).1 | MT3458(65–88).1 | MT3502(36–57).1 |
| MT3502(63–80).1 | MT350282.1 | MT35840(1,2).1 | MT3586(37–75).1 | MT3586(77–85).1 |
| MT358(687–703).1 | MT3587(05–26).1 | MT3587(28–48).1 | MT35986(5,6).1 | MT3650(25–33).1 |
| MT370516.1 | MT370518.1 | MT370831.1 | MT37083(3–9).1 | MT37084(1–6).1 |
| MT3708(48–54).1 | MT3708(56–66).1 | MT3708(68–72).1 | MT37087(4–8).1 | MT37088(0–3).1 |
| MT37088(5,6).1 | MT3708(88–92).1 | MT37089(4–9).1 | MT37090(1–9).1 | MT37091(1–7).1 |
| MT3709(19,20).1 | MT370925.1 | MT370927.1 | MT37093(0–5).1 | MT370937.1 |
| MT3709(39,40).1 | MT370942.1 | MT370944.1 | MT3709(46–50).1 | MT37095(2–4).1 |
| MT37095(6–8).1 | MT37096(0–2).1 | MT37096(4–7).1 | MT37097(0–2).1 | MT37097(5–7).1 |
| MT3709(79–81).1 | MT37098(4–8).1 | MT370991.1 | MT37099(3,4).1 | MT37099(6–8).1 |
| MT37100(0–2).1 | MT37100(6–9).1 | MT37101(2,3).1 | MT37101(5–9).1 | MT37102(2,3).1 |
| MT37102(6–9).1 | MT37103(2–6).1 | MT371038.1 | MT3710(47–50).1 | MT3715(68–74).1 |
| MT37248(0–2).1 | MT3741(01–16).1 | MT3754(28–38).1 | MT3754(40–63).1 | MT3754(65–72).1 |
| MT375474.1 | MT37547(6–8).1 | MT37548(0–3).1 | MT3807(28–34).1 | MT3854(14–79).1 |
| MT385482.1 | MT385485.1 | MT3945(28–31).1 | MT396241.1 | MT396266.1 |
| MT4076(49–59).1 | MT412134.1 | MT412155.1 | MT412170.1 | MT412173.1 |
| MT41217(6–8).1 | MT4122(26–30).1 | MT4122(33–59).1 | MT41226(1–4).1 | MT41226(7–9).1 |
| MT4122(71–81).1 | MT412(283–314).1 | MT41231(6–9).1 | MT4123(21–34).1 | MT41233(6,7).1 |
| MT41532(0–3).1 | MT415(894–912).1 | MT41672(5,6).2 | MT4188(80–92).1 | MT419810.1 |
| MT419812.1 | MT41981(4–6).1 | MT419818.1 | MT419820.1 | MT4198(29–32).1 |
| MT41983(4–7).1 | MT419839.1 | MT41984(1,2).1 | MT41984(6,7).1 | MT419851.1 |
| MT41985(3,4).1 | MT419860.1 | MT42280(6,7).1 | MT42855(1–4).1 | MT4291(83–91).1 |
| MT432195.1 | MT434(782–817).1 | MT4350(79–86).1 | MT4387(18–25).1 | MT4387(27–32).1 |
| MT4387(34–44).1 | MT4387(46–60).1 | MT43923(3,4).1 | MT439237.1 | MT439239.1 |
| MT439242.1 | MT43924(4,5).1 | MT439247.1 | MT4392(49–54).1 | MT439256.1 |
| MT439258.1 | MT43926(0–3).1 | MT439265.1 | MT43926(7,8).1 | MT43927(0,1).1 |
| MT439273.1 | MT4392(79,80).1 | MT43928(2–4).1 | MT4392(87–93).1 | MT43929(5–7).1 |
| MT439300.1 | MT43930(2,3).1 | MT439307.1 | MT43931(0–2).1 | MT43931(4–8).1 |

(continued on next page)

(Table S1 continued)

| | | | | |
|---|---|---|---|---|
| MT444148.1 | MT444540.1 | MT444619.1 | MT44462(4–6).1 | MT444633.1 |
| MT444635.1 | MT446312.1 | MT446339.1 | MT4463(42–50).1 | MT44635(2–4).1 |
| MT44635(6–9).1 | MT44715(4–9).1 | MT44716(1–3).1 | MT44716(5,6).1 | MT447169.1 |
| MT44717(1–7).1 | MT44963(6,7).1 | MT449639.1 | MT449641.1 | MT449644.1 |
| MT449647.1 | MT449651.1 | MT449656.1 | MT44966(4,5).1 | MT449670.1 |
| MT44967(2,3).1 | MT4508(72–87).1 | MT450(889–907).1 | MT4509(09–15).1 | MT45091(7,8).1 |
| MT450932.1 | MT45093(5,6).1 | MT450938.1 | MT45094(1–4).1 | MT450946.1 |
| MT4509(48–51).1 | MT45095(4–7).1 | MT45096(0–3).1 | MT45096(5,6).1 | MT4509(68–74).1 |
| MT450977.1 | MT45098(0–5).1 | MT45(0988–1000).1 | MT451002.1 | MT45100(4–9).1 |
| MT451013.1 | MT451015.1 | MT4510(17–22).1 | MT451024.1 | MT451026.1 |
| MT451033.1 | MT4510(35–42).1 | MT45104(4–9).1 | MT45105(1,2).1 | MT4510(56–62).1 |
| MT45106(5,6).1 | MT451072.1 | MT451074.1 | MT45107(6,7).1 | MT451079.1 |
| MT451081.1 | MT45108(6–8).1 | MT451090.1 | MT45109(2–4).1 | MT451098.1 |
| MT45110(0–5).1 | MT45110(8,9).1 | MT45111(4–9).1 | MT4511(21–92).1 | MT45119(4–9).1 |
| MT45120(1–4).1 | MT4512(07–20).1 | MT45122(4–7).1 | MT4512(29–33).1 | MT4512(35–42).1 |
| MT45124(4–8).1 | MT451253.1 | MT45125(5–9).1 | MT451261.1 | MT45126(3–5).1 |
| MT451269.1 | MT451272.1 | MT451274.1 | MT451276.1 | MT45127(8,9).1 |
| MT451281.1 | MT45128(3,4).1 | MT451324.1 | MT45135(4–6).1 | MT451358.1 |
| MT451366.1 | MT451371.1 | MT45137(5,6).1 | MT451386.1 | MT451389.1 |
| MT45139(1,2).1 | MT45141(0–2).1 | MT451415.1 | MT451434.1 | MT451436.1 |
| MT451439.1 | MT45144(1,2).1 | MT45144(7–9).1 | MT451452.1 | MT451456.1 |
| MT451464.1 | MT451466.1 | MT451470.1 | MT451476.1 | MT451478.1 |
| MT451488.1 | MT451496.1 | MT451549.1 | MT451569.1 | MT451578.1 |
| MT45158(0–2).1 | MT451584.1 | MT451586.1 | MT451590.1 | MT451592.1 |
| MT451597.1 | MT451599.1 | MT45160(1–4).1 | MT45160(7–9).1 | MT451611.1 |
| MT451614.1 | MT451616.1 | MT451641.1 | MT451646.1 | MT451651.1 |
| MT451653.1 | MT451655.1 | MT451657.1 | MT45166(6–8).1 | MT45167(4,5).1 |
| MT451680.1 | MT451684.1 | MT451692.1 | MT451694.1 | MT451696.1 |

(continued on next page)

(Table S1 continued)

| | | | | |
|---|---|---|---|---|
| MT451704.1 | MT451708.1 | MT45171(0–3).1 | MT451716.1 | MT451718.1 |
| MT451723.1 | MT451726.1 | MT451728.1 | MT451730.1 | MT45173(3,4).1 |
| MT45173(6,7).1 | MT451740.1 | MT451742.1 | MT45174(8,9).1 | MT451751.1 |
| MT45175(4,5).1 | MT451762.1 | MT451771.1 | MT45177(5–7).1 | MT451779.1 |
| MT451783.1 | MT451786.1 | MT451874.1 | MT45187(6–8).1 | MT4518(80–90).1 |
| MT45257(4–6).1 | MT457(390–403).1 | MT459(832–917).1 | MT4599(19–25).1 | MT459979.1 |
| MT4599(85–90).1 | MT4600(89–92).1 | MT46011(3–7).1 | MT4601(19–21).1 | MT46012(4,5).1 |
| MT4601(27–30).1 | MT46013(2,3).1 | MT4601(35–40).1 | MT4616(03–60).1 | MT46166(2–4).1 |
| MT46166(6–8).1 | MT466071.1 | MT466583.1 | MT4672(37–63).1 | MT4701(00–79).1 |
| MT470219.1 | MT47262(1–7).1 | MT4741(26–30).1 | MT47413(2–8).1 | MT47638(4,5).1 |
| MT4778(35–62).1 | MT477885.1 | MT47790(3,4).1 | MT479216.1 | MT47922(3–7).1 |
| MT481(895–909).1 | MT4819(27–36).1 | MT481992.1 | MT48211(0,1).1 | MT48211(3–9).1 |
| MT4821(21–49).1 | MT4835(53–60).1 | MT483563.1 | MT483702.1 | MT483867.1 |
| MT4838(74–81).1 | MT49697(2–7).1 | MT4969(79–97).1 | MT499(171–210).1 | MT499215.1 |
| MT4992(17–20).1 | MT500122.1 | MT502774.1 | MT506201.1 | MT506203.1 |
| MT506205.1 | MT506209.1 | MT506221.1 | MT50653(0–5).1 | MT506537.1 |
| MT5065(39–41).1 | MT506544.1 | MT506546.1 | MT50656(5–8).1 | MT50657(0–3).1 |
| MT506581.1 | MT50664(2,3).1 | MT5066(48–52).1 | MT506654.1 | MT5066(58–60).1 |
| MT50666(3,4).1 | MT506666.1 | MT506668.1 | MT506670.1 | MT50667(4,5).1 |
| MT506677.1 | MT5066(79–81).1 | MT506683.1 | MT506685.1 | MT506687.1 |
| MT5066(89,90).1 | MT50669(3–7).1 | MT50670(0,1).1 | MT506707.1 | MT506(885–907).1 |
| MT506981.1 | MT507018.1 | MT507032.2 | MT5072(71–82).1 | MT50779(3–6).1 |
| MT5094(52–68).1 | MT50947(1,2).1 | MT50947(4,5).1 | MT509477.1 | MT509480.1 |
| MT509486.1 | MT509(488–512).1 | MT5096(49–51).1 | MT50965(6–9).1 | MT509669.1 |
| MT509674.1 | MT509683.1 | MT509685.1 | MT50970(2,3).1 | MT50995(8,9).1 |
| MT510(690–703).1 | MT51071(8,9).1 | MT51072(1,2).1 | MT51072(5–8).1 | MT510999.1 |
| MT511059.1 | MT5110(65–84).1 | MT5124(15–31).1 | MT5124(33–47).1 | MT51264(4,5).1 |
| MT513758.1 | MT5174(20–34).1 | MT51743(6,7).1 | MT517443.1 | MT52017(2–4).1 |

(continued on next page)

(Table S1 continued)

| | | | | |
|---|---|---|---|---|
| MT52017(6–8).1 | MT52018(0–9).1 | MT520(191–201).1 | MT5202(03–10).1 | MT5202(12–25).1 |
| MT5202(28–40).1 | MT5202(43–66).1 | MT5202(69–73).1 | MT5202(76–81).1 | MT52028(3–5).1 |
| MT5202(87–92).1 | MT52029(5,6).1 | MT520(298–302).1 | MT5203(04–10).1 | MT5203(12–21).1 |
| MT52032(3–5).1 | MT5203(27–33).1 | MT5203(36–53).1 | MT5203(55–77).1 | MT5203(79–92).1 |
| MT520394.1 | MT520(396–424).1 | MT52042(6–9).1 | MT52043(1–7).1 | MT5204(39–41).1 |
| MT52044(3–8).1 | MT52045(1–5).1 | MT5204(57–61).1 | MT52046(3–7).1 | MT520469.1 |
| MT52047(1–5).1 | MT52047(7–9).1 | MT5204(81–90).1 | MT52049(2–8).1 | MT5205(00–12).1 |
| MT5205(15–23).1 | MT52052(5,6).1 | MT5205(28–30).1 | MT5205(32–44).1 | MT52054(6,7).1 |
| MT52285(2–5).1 | MT525950.1 | MT527178.1 | MT527184.1 | MT528235.1 |
| MT52823(7–9).1 | MT528(598–631).1 | MT531537.2 | MT53243(5–8).1 | MT532557.1 |
| MT533(197–206).1 | MT533208.1 | MT533210.1 | MT533212.1 | MT533215.1 |
| MT53321(7–9).1 | MT53322(1–3).1 | MT5332(25–30).1 | MT53323(2–8).1 | MT534285.1 |
| MT53430(4–9).1 | MT53431(1–8).1 | MT534320.1 | MT5343(22–39).1 | MT534630.1 |
| MT535472.1 | MT535481.1 | MT5354(86–97).1 | MT535(499–503).1 | MT535505.1 |
| MT53550(8,9).1 | MT5361(76–90).1 | MT536956.1 | MT53696(1,2).1 | MT53696(4,5).1 |
| MT53697(0–2).1 | MT53697(4–7).1 | MT5391(58–60).1 | MT5391(62–76).1 | MT539726.1 |
| MT539729.1 | MT547814.1 | MT55160(4–9).1 | MT554052.1 | MT5575(68–70).1 |
| MT558(647–707).1 | MT55903(7,8).1 | MT55980(0–3).1 | MT560525.1 | MT56053(0,1).1 |
| MT56065(6,7).1 | MT5606(67–75).1 | MT560677.1 | MT5606(79,80).1 | MT560682.1 |
| MT5606(89,90).1 | MT56069(2–4).1 | MT56070(4–6).1 | MT560827.1 | MT56549(5–9).1 |
| MT56643(5–7).1 | MT5686(34–41).1 | MT568645.1 | MT56947(0–4).1 | MT5760(25–35).1 |
| MT5760(37–49).1 | MT5760(52–61).1 | MT5765(29–32).1 | MT576584.1 | MT5766(39–45).1 |
| MT576689.1 | MT5770(09–15).1 | MT577359.1 | MT577627.1 | MT57801(5–7).1 |
| MT58245(3,4).1 | MT582456.1 | MT5824(58–63).1 | MT58246(5–9).1 | MT5824(71–88).1 |
| MT582491.1 | MT582494.1 | MT582496.1 | MT58249(8,9).1 | MT584985.1 |
| MT58499(7,8).1 | MT585003.1 | MT58501(0,1).1 | MT585014.1 | MT585019.1 |
| MT585027.1 | MT585032.1 | MT585036.1 | MT585039.1 | MT585042.1 |
| MT58504(4,5).1 | MT585048.1 | MT585050.1 | MT585052.1 | MT5850(58–61).1 |

(continued on next page)

(Table S1 continued)

| | | | | |
|---|---|---|---|---|
| MT585064.1 | MT58506(6,7).1 | MT585071.1 | MT585077.1 | MT585(082–104).1 |
| MT590(598–600).1 | MT594109.1 | MT594112.1 | MT59440(1,2).1 | MT59817(1–8).1 |
| MT59863(3,4).1 | MT598636.1 | MT5986(39–42).1 | MT6012(75–95).1 | MT60581(5–8).1 |
| MT6072(44–55).1 | MT6076(01–21).1 | MT608648.1 | MT609(558–600).1 | MT609918.1 |
| MT610913.1 | MT61144(0–4).1 | MT6114(48–51).1 | MT611527.1 | MT611530.1 |
| MT61153(6–9).1 | MT611968.1 | MT61434(7–9).1 | MT61435(6,7).1 | MT614447.1 |
| MT6144(49,50).1 | MT614(452–510).1 | MT61451(2–7).1 | MT6145(19–61).1 | MT614563.1 |
| MT614(595–601).1 | MT62076(6,7).1 | MT6207(69–72).1 | MT621560.1 | MT622319.1 |
| MT622321.1 | MT624728.1 | MT624733.1 | MT627213.1 | MT62721(6,7).1 |
| MT627219.1 | MT627221.1 | MT627223.1 | MT62722(5–9).1 | MT62723(1–8).1 |
| MT627(240–300).1 | MT6273(02–12).1 | MT62731(4–8).1 | MT627325.1 | MT6273(89–96).1 |
| MT6274(04–37).1 | MT627607.1 | MT627609.1 | MT62761(2–6).1 | MT6276(18–38).1 |
| MT6277(45–52).1 | MT6277(54–67).1 | MT6279(20–32).1 | MT6280(37–71).1 | MT6280(73–82).1 |
| MT62808(4,5).1 | MT6280(87–93).1 | MT628095.1 | MT62809(7,8).1 | MT62810(0–2).1 |
| MT62810(4,5).1 | MT628110.1 | MT628113.1 | MT628116.1 | MT62811(8,9).1 |
| MT628121.1 | MT6281(23–43).1 | MT6281(45–50).1 | MT62815(2–8).1 | MT628160.1 |
| MT62816(3–7).1 | MT6281(69–78).1 | MT62818(0,1).1 | MT628184.1 | MT62818(6,7).1 |
| MT628(190–225).1 | MT62822(7,8).1 | MT62823(0–4).1 | MT628236.1 | MT628238.1 |
| MT6282(41–76).1 | MT6282(79,80).1 | MT6304(21–32).1 | MT6317(84–95).1 | MT631797.1 |
| MT631(799,800).1 | MT63180(2,3).1 | MT631805.1 | MT63180(7–9).1 | MT631813.1 |
| MT6318(15–25).1 | MT6318(27–31).1 | MT631833.1 | MT6318(37–42).1 | MT632491.1 |
| MT632(493–529).1 | MT6325(31–67).1 | MT6325(69–86).1 | MT6325(88–91).1 | MT632(593–616).1 |
| MT6326(18–27).1 | MT6326(29–32).1 | MT632766.1 | MT6327(69–73).1 | MT632(775–808).1 |
| MT6328(10–61).1 | MT6328(63–72).1 | MT6328(74–90).1 | MT63289(2–8).1 | MT6329(00–14).1 |
| MT63(2916–3011).1 | MT6330(13–39).1 | MT633041.1 | MT63304(4–6).1 | MT635255.1 |
| MT6352(69–72).1 | MT635328.1 | MT635339.1 | MT63539(1–3).1 | MT635397.1 |
| MT6354(03–10).1 | MT635445.1 | MT635641.1 | MT635643.1 | MT6356(45–55).1 |
| MT635672.1 | MT63585(5–8).1 | MT637143.1 | MT64148(5,6).1 | MT6414(88–90).1 |

(continued on next page)

(Table S1 continued)

| | | | | |
|---|---|---|---|---|
| MT641(492–506).1 | MT6415(08–30).1 | MT6415(32–96).1 | MT641(598–615).1 | MT6416(17–36).1 |
| MT6416(39–46).1 | MT641648.1 | MT641651.1 | MT641656.1 | MT64166(1–7).1 |
| MT641669.1 | MT641674.1 | MT64167(6–9).1 | MT64168(1,2).1 | MT641686.1 |
| MT641689.1 | MT64169(2,3).1 | MT641697.1 | MT641699.1 | MT641708.1 |
| MT641710.1 | MT64172(0–4).1 | MT6417(28–31).1 | MT641733.1 | MT6417(35–46).1 |
| MT641748.1 | MT64175(0–4).1 | MT6417(57–64).1 | MT64176(6–8).1 | MT641774.1 |
| MT64177(6–8).1 | MT64206(8,9).1 | MT64207(2–4).1 | MT6420(78–83).1 | MT64208(5–9).1 |
| MT642(091–107).1 | MT642109.1 | MT6421(11–34).1 | MT642(136–207).1 | MT6422(09–35).1 |
| MT6422(38–73).1 | MT642(275–334).1 | MT6423(36–53).1 | MT6423(55–67).1 | MT6423(69–75).1 |
| MT6423(77–85).1 | MT642(387–435).1 | MT644268.1 | MT64605(3,4).1 | MT64606(4,5).1 |
| MT6460(68–84).1 | MT6460(86–94).1 | MT64609(6–8).1 | MT646101.1 | MT646106.1 |
| MT646112.1 | MT64611(6,7).1 | MT646120.1 | MT648676.1 | MT648830.1 |
| MT653094.1 | MT653097.1 | MT653099.1 | MT65513(1–5).1 | MT65574(2–6).1 |
| MT65575(0,1).1 | MT655948.1 | MT65601(0,1).1 | MT65625(6,7).1 | MT65659(0,1).1 |
| MT657271.1 | MT657275.1 | MT657958.1 | MT658264.1 | MT65850(0–5).1 |
| MT65850(7–9).1 | MT661524.1 | MT66410(5–7).1 | MT664109.1 | MT66411(7,8).1 |
| MT664143.1 | MT664161.1 | MT6641(69–72).1 | MT664175.1 | MT664197.1 |
| MT66420(1–3).1 | MT664205.1 | MT664209.1 | MT664727.1 | MT664729.1 |
| MT664774.1 | MT664793.1 | MT66479(5,6).1 | MT66480(7,8).1 | MT664822.1 |
| MT664986.1 | MT664990.1 | MT665006.1 | MT665028.1 | MT665970.1 |
| MT665972.1 | MT665974.1 | MT666042.1 | MT666068.1 | MT666099.1 |
| MT667351.1 | MT668716.1 | MT66932(1,2).1 | MT670008.1 | MT67001(3–5).1 |
| MT67001(7–9).1 | MT67002(1–3).1 | MT6718(10–28).1 | MT675289.1 | MT675517.1 |
| MT675933.1 | MT6759(37–45).1 | MT67595(0–2).1 | MT67595(4–9).1 | MT67601(3,4).1 |
| MT6762(88–91).1 | MT67636(6–9).1 | MT6763(88–90).1 | MT676397.1 | MT678839.1 |
| MT6791(55–96).1 | MT679198.1 | MT67920(1,2).1 | MT67920(5,6).1 | MT679208.1 |
| MT6802(17–42).1 | MT682732.1 | MT6833(86–95).1 | MT683397.1 | MT683399.1 |
| MT68340(3,4).1 | MT6834(06–13).1 | MT68341(6–8).1 | MT703677.1 | MT70388(3,4).1 |

(continued on next page)

(Table S1 continued)

| | | | | |
|---|---|---|---|---|
| MT703958.1 | MT703963.1 | MT703969.1 | MT703973.1 | MT70397(5–9).1 |
| MT70398(2–5).1 | MT70(3989–4001).1 | MT70400(3–9).1 | MT70401(1,2).1 | MT704014.1 |
| MT70401(7–9).1 | MT70402(2–6).1 | MT7040(29,30).1 | MT704033.1 | MT7040(37–47).1 |
| MT7040(49–52).1 | MT70405(5,6).1 | MT70406(0,1).1 | MT7040(63–75).1 | MT704077.1 |
| MT70408(0,1).1 | MT70408(4,5).1 | MT704088.1 | MT70409(4,5).1 | MT70410(0–2).1 |
| MT70411(1,2).1 | MT70411(4–7).1 | MT7041(19–29).1 | MT70413(1–6).1 | MT70431(0–7).1 |
| MT704812.1 | MT7048(16–25).1 | MT70520(5,6).1 | MT706050.1 | MT70613(2–8).1 |
| MT70614(0–6).1 | MT706(148–208).1 | MT706210.1 | MT70621(2–6).1 | MT7062(18–41).1 |
| MT7062(43–56).1 | MT7062(58–60).1 | MT70626(2–6).1 | MT7062(68–73).1 | MT70627(5–7).1 |
| MT7062(79–82).1 | MT706(284–315).1 | MT7063(17–76).1 | MT7063(79–87).1 | MT706(389–444).1 |
| MT7064(46–51).1 | MT706453.1 | MT70910(4,5).1 | MT710714.1 | MT71187(2,3).1 |
| MT711880.1 | MT712859.1 | MT712861.1 | MT712863.1 | MT730002.1 |
| MT731285.1 | MT731292.1 | MT731327.1 | MT731346.1 | MT731468.1 |
| MT7316(53–69).1 | MT731673.1 | MT7317(33–44).1 | MT731746.1 | MT731764.1 |
| MT73193(1–8).1 | MT733120.1 | MT734046.1 | MT734055.1 | MT738101.1 |
| MT738173.1 | MT7394(25–43).1 | MT739444.2 | MT73944(5,6).1 | MT7394(48–58).1 |
| MT7394(60–77).1 | MT74038(1–4).1 | MT74039(0–7).1 | MT74040(1–4).1 | MT74043(1,2).1 |
| MT74043(4–6).1 | MT7404(38–41).1 | MT740444.1 | MT7404(76–80).1 | MT740483.1 |
| MT740485.1 | MT74061(7–9).1 | MT740684.1 | MT740686.1 | MT740688.1 |
| MT740706.1 | MT740708.1 | MT740717.1 | MT74072(3,4).1 | MT740820.1 |
| MT740896.1 | MT740923.1 | MT741096.1 | MT741490.1 | MT7420(87–92).1 |
| MT74278(2–4).1 | MT745584.1 | MT745601.1 | MT745605.1 | MT745608.1 |
| MT74561(0,1).1 | MT7456(29–31).1 | MT7456(39,40).1 | MT745642.1 | MT74564(4–6).1 |
| MT7456(48–74).1 | MT745(676–749).1 | MT745767.1 | MT745836.1 | MT745875.1 |
| MT747438.1 | MT747978.1 | MT748758.1 | MT7500(20–67).1 | MT7500(69–75).1 |
| MT7500(78–81).1 | MT75008(3,4).1 | MT750(087–116).1 | MT75012(1,2).1 | MT750124.1 |
| MT75013(1,2).1 | MT75013(5,6).1 | MT750139.1 | MT75014(2–5).1 | MT75014(8,9).1 |
| MT7501(51–82).1 | MT7501(86–92).1 | MT75019(4,5).1 | MT750(197–201).1 | MT7503(33–44).1 |

(continued on next page)

(Table S1 continued)

| | | | | |
|---|---|---|---|---|
| MT75034(6–8).1 | MT75035(0–9).1 | MT7503(61–74).1 | MT7503(77–81).1 | MT75038(3–7).1 |
| MT7503(89,90).1 | MT75039(3–7).1 | MT75040(1,2).1 | MT7504(04–27).1 | MT7504(29–34).1 |
| MT75043(6,7).1 | MT7504(39–44).1 | MT75044(6–8).1 | MT75045(0–3).1 | MT7504(55–60).1 |
| MT7504(62–88).1 | MT75049(0–3).1 | MT750495.1 | MT75049(7,8).1 | MT75050(0–7).1 |
| MT7505(09,10).1 | MT755582.1 | MT755585.1 | MT755587.1 | MT75559(3,4).1 |
| MT755(599,600).1 | MT755602.1 | MT7558(27–38).1 | MT75585(7,8).1 | MT755(883–901).1 |
| MT75591(6,7).1 | MT75595(5,6).1 | MT755973.1 | MT757006.1 | MT75701(2,3).1 |
| MT757016.1 | MT7570(18–22).1 | MT75702(4,5).1 | MT757027.1 | MT7570(29,30).1 |
| MT75703(2,3).1 | MT7570(37–41).1 | MT75704(3,4).1 | MT757047.1 | MT757049.1 |
| MT757051.1 | MT757054.1 | MT757056.1 | MT757058.1 | MT75706(3,4).1 |
| MT75706(6–8).1 | MT7570(70–83).1 | MT757085.1 | MT757087.1 | MT757094.1 |
| MT757096.1 | MT757100.1 | MT75710(4,5).1 | MT75711(0,1).1 | MT75711(5–7).1 |
| MT757119.1 | MT75712(1–3).1 | MT758193.1 | MT758197.1 | MT75820(2,3).1 |
| MT758213.1 | MT758215.1 | MT758240.1 | MT758314.1 | MT758346.1 |
| MT758375.1 | MT758402.1 | MT758421.1 | MT758436.1 | MT758439.1 |
| MT758471.1 | MT758683.1 | MT758701.1 | MT759582.1 | MT759588.1 |
| MT759714.1 | MT759742.1 | MT759820.1 | MT759844.1 | MT760790.1 |
| MT76239(6–8).1 | MT76294(2–4).1 | MT764166.1 | MT764170.1 | MT764172.1 |
| MT764174.1 | MT7641(77–81).1 | MT76690(7,8).1 | MT7720(85–91).1 | MT772210.1 |
| MT772215.1 | MT772237.1 | MT772240.1 | MT77225(2,3).1 | MT772256.1 |
| MT772271.1 | MT77229(3,4).1 | MT772297.1 | MT77240(3–5).1 | MT7724(07–12).1 |
| MT77241(6–9).1 | MT77242(1–3).1 | MT77242(5–8).1 | MT772430.1 | MT77243(2–8).1 |
| MT77244(0,1).1 | MT77244(3–5).1 | MT7724(47–50).1 | MT77245(2–5).1 | MT7724(57–67).1 |
| MT772469.1 | MT77247(1–7).1 | MT7724(79–91).1 | MT772(494–507).1 | MT7725(09,10).1 |
| MT772512.1 | MT77251(4–6).1 | MT7725(18–29).1 | MT77253(1–9).1 | MT772541.1 |
| MT7725(43–59).1 | MT7725(61–71).1 | MT77257(3–5).1 | MT7725(77–80).1 | MT77313(3,4).1 |
| MT7758(26–33).1 | MT7764(01–27).1 | MT776904.1 | MT777677.1 | MT77882(7–9).1 |
| MT78141(4,5).1 | MT782348.1 | MT782361.1 | MT782364.1 | MT7823(69–71).1 |

(continued on next page)

(Table S1 continued)

| | | | | |
|---|---|---|---|---|
| MT78237(7,8).1 | MT78238(1,2).1 | MT786327.1 | MT786329.1 | MT786331.1 |
| MT78633(4–7).1 | MT786422.1 | MT786438.1 | MT786442.1 | MT786444.1 |
| MT786455.1 | MT786(798–802).1 | MT786804.1 | MT7868(09–15).1 | MT7868(17–40).1 |
| MT786842.1 | MT7868(44–51).1 | MT786859.1 | MT786865.1 | MT786868.1 |
| MT786941.1 | MT786943.1 | MT786946.1 | MT787473.1 | MT787486.1 |
| MT787500.1 | MT787503.1 | MT787553.1 | MT787564.1 | MT787577.1 |
| MT78764(6–8).1 | MT787650.1 | MT787742.1 | MT78774(5,6).1 | MT787749.1 |
| MT789688.1 | MT789690.1 | MT789712.1 | MT790522.1 | MT791312.1 |
| MT79184(4,5).1 | MT79185(0–2).1 | MT79185(4,5).1 | MT79185(7,8).1 | MT79186(1,2).1 |
| MT791864.1 | MT7918(68–70).1 | MT791878.1 | MT791880.1 | MT791883.1 |
| MT791885.1 | MT791891.1 | MT79189(5,6).1 | MT791899.1 | MT791907.1 |
| MT79191(4,5).1 | MT791920.1 | MT791923.1 | MT791926.1 | MT7919(36–50).1 |
| MT7919(53–63).1 | MT79587(0–6).1 | MT7958(78–84).1 | MT7958(87–96).1 | MT795(898–902).1 |
| MT79590(4–7).1 | MT7959(09–13).1 | MT7977(58–62).1 | MT79776(4,5).1 | MT79776(8,9).1 |
| MT79777(1,2).1 | MT798592.1 | MT799967.1 | MT7999(69,70).1 | MT799972.1 |
| MT799974.1 | MT799976.1 | MT799991.1 | MT799998.1 | MT800002.1 |
| MT800005.1 | MT800010.1 | MT80001(3–6).1 | MT800021.1 | MT80002(3,4).1 |
| MT8000(29–31).1 | MT80003(5–7).1 | MT800041.1 | MT800043.1 | MT800045.1 |
| MT800047.1 | MT800050.1 | MT800105.1 | MT800282.1 | MT800284.1 |
| MT800505.1 | MT800515.1 | MT800542.1 | MT800753.1 | MT800758.1 |
| MT800794.1 | MT800816.1 | MT800818.1 | MT800820.1 | MT800845.1 |
| MT800873.1 | MT80089(5,6).1 | MT80092(3–5).1 | MT800928.1 | MT800934.1 |
| MT800936.1 | MT8009(39,40).1 | MT800942.1 | MT80097(7,8).1 | MT80099(4,5).1 |
| MT800997.1 | MT80(0999–1004).1 | MT801006.1 | MT801043.1 | MT801048.1 |
| MT801051.1 | MT806104.1 | MT806176.1 | MT80619(1,2).1 | MT8067(56–83).1 |
| MT806(785–809).1 | MT806884.1 | MT807936.1 | MT808071.1 | MT810084.1 |
| MT81008(6–9).1 | MT8104(77–81).1 | MT810483.1 | MT81048(5–9).1 | MT810(491–514).1 |
| MT810516.1 | MT81051(8,9).1 | MT8105(21–42).1 | MT8105(44–68).1 | MT81057(0–5).1 |

(continued on next page)

(Table S1 continued)

| | | | | |
|---|---|---|---|---|
| MT8105(77–82).1 | MT810(584–621).1 | MT8106(23–52).1 | MT81065(4,5).1 | MT8106(57–73).1 |
| MT8106(75–88).1 | MT81069(0,1).1 | MT81069(3–5).1 | MT810(697–710).1 | MT81071(2–4).1 |
| MT8107(16–25).1 | MT81072(7,8).1 | MT81073(0,1).1 | MT810758.1 | MT810761.1 |
| MT81076(4–9).1 | MT81077(1–6).1 | MT8107(78–84).1 | MT81078(6–9).1 | MT81081(4–6).1 |
| MT810819.1 | MT8108(21–33).1 | MT81083(5–9).1 | MT81084(1–3).1 | MT81084(6–8).1 |
| MT81085(0–2).1 | MT81085(5–9).1 | MT810861.1 | MT810863.1 | MT81086(5–7).1 |
| MT8108(69,70).1 | MT81087(2–7).1 | MT810879.1 | MT81088(1–6).1 | MT810(888–903).1 |
| MT8109(05–11).1 | MT810913.1 | MT81091(5–7).1 | MT8109(19,20).1 | MT810922.1 |
| MT810924.1 | MT8109(26–30).1 | MT81093(2,3).1 | MT810936.1 | MT81094(1–3).1 |
| MT81094(6,7).1 | MT8109(49–53).1 | MT810955.1 | MT81095(7,8).1 | MT81096(1–4).1 |
| MT81096(6,7).1 | MT810969.1 | MT81097(1–3).1 | MT8109(75–98).1 | MT81100(0–3).1 |
| MT81100(6–8).1 | MT8110(10–22).1 | MT811162.1 | MT81116(4–6).1 | MT811169.1 |
| MT811171.1 | MT811173.1 | MT81117(5–7).1 | MT8111(79–88).1 | MT81119(0,1).1 |
| MT811194.1 | MT811(196–210).1 | MT811212.1 | MT8112(15–22).1 | MT811224.1 |
| MT8112(26–31).1 | MT81123(3–8).1 | MT81124(0–2).1 | MT8112(44–59).1 | MT8112(61–72).1 |
| MT8112(74–83).1 | MT811286.1 | MT8112(88–90).1 | MT811292.1 | MT811(294–318).1 |
| MT81132(0–6).1 | MT8113(28–41).1 | MT8113(43–69).1 | MT81137(1,2).1 | MT81137(4,5).1 |
| MT811378.1 | MT811380.1 | MT8113(82–90).1 | MT81139(2–5).1 | MT811(397–401).1 |
| MT8114(03–14).1 | MT811416.1 | MT81142(0–4).1 | MT811426.1 | MT8114(28–30).1 |
| MT8114(32–43).1 | MT81144(5–8).1 | MT811451.1 | MT81145(3,4).1 | MT811456.1 |
| MT8114(58–61).1 | MT811(463–523).1 | MT8115(25–35).1 | MT81154(4–6).1 | MT8115(48–58).1 |
| MT81156(0–2).1 | MT81156(4–9).1 | MT811639.1 | MT811641.1 | MT811649.1 |
| MT81165(4,5).1 | MT8116(57–60).1 | MT811662.1 | MT811664.1 | MT811667.1 |
| MT8116(69–73).1 | MT8116(75–83).1 | MT81168(8,9).1 | MT811693.1 | MT811(695–703).1 |
| MT8117(05–16).1 | MT8117(18–29).1 | MT81173(1,2).1 | MT811734.1 | MT81173(7,8).1 |
| MT81174(0,1).1 | MT81174(3–5).1 | MT81174(7–9).1 | MT81175(1–3).1 | MT811755.1 |
| MT811758.1 | MT814636.1 | MT814698.1 | MT8201(27–32).1 | MT8204(61–84).1 |
| MT82048(6–9).1 | MT8215(30–43).1 | MT8215(45–75).1 | MT8215(77–82).1 | MT821(584–632).1 |

(continued on next page)

(Table S1 continued)

| | | | | |
|---|---|---|---|---|
| MT8216(34–48).1 | MT821650.1 | MT82165(2–6).1 | MT8216(58–66).1 | MT8216(68–71).1 |
| MT82167(3,4).1 | MT8216(76–96).1 | MT821(698–705).1 | MT821707.1 | MT8217(09–11).1 |
| MT82171(3–8).1 | MT8217(20–35).1 | MT8217(37–50).1 | MT82175(2–6).1 | MT8217(58–78).1 |
| MT8217(80–93).1 | MT821795.1 | MT825091.1 | MT82707(4,5).1 | MT827190.1 |
| MT82720(2,3).1 | MT8272(06–53).1 | MT8278(09–13).1 | MT8278(15–29).1 | MT82783(1–7).1 |
| MT8278(39–43).1 | MT827845.1 | MT827872.1 | MT827940.1 | MT83112(2–5).1 |
| MT8311(27–31).1 | MT83113(3–5).1 | MT8311(37–45).1 | MT8311(47–54).1 | MT8311(56–84).1 |
| MT831(186–202).1 | MT8312(04–27).1 | MT8312(29–41).1 | MT83124(3–6).1 | MT831(248–316).1 |
| MT8313(18–82).1 | MT83138(5–7).1 | MT831(389–429).1 | MT8314(31–65).1 | MT8314(67–80).1 |
| MT83148(2–6).1 | MT8315(31–50).1 | MT83155(2–9).1 | MT8315(61–91).1 | MT83159(3–9).1 |
| MT8316(01–21).1 | MT83162(3–7).1 | MT8316(29–46).1 | MT8316(48–50).1 | MT831(652–711).1 |
| MT83171(3,4).1 | MT8317(16–57).1 | MT8317(60–96).1 | MT831(798–800).1 | MT831802.1 |
| MT83180(4–9).1 | MT831811.1 | MT831813.1 | MT83181(5,6).1 | MT8318(18–30).1 |
| MT8318(32–48).1 | MT8318(50–63).1 | MT8318(65–70).1 | MT8318(72–81).1 | MT8318(83–92).1 |
| MT831(894–900).1 | MT833958.1 | MT83396(0–2).1 | MT83396(4–9).1 | MT833972.1 |
| MT8339(74–82).1 | MT83(3985–4017).1 | MT8340(19–50).1 | MT83405(2–5).1 | MT8340(57–62).1 |
| MT8340(64–80).1 | MT83408(2–4).1 | MT8340(86–97).1 | MT834(099–121).1 | MT8341(23–30).1 |
| MT83413(2–9).1 | MT83414(1–8).1 | MT8341(50–78).1 | MT8341(80–93).1 | MT83419(5–9).1 |
| MT834201.1 | MT83420(3,4).1 | MT8342(06–14).1 | MT8342(16–39).1 | MT83424(1–7).1 |
| MT8342(49–56).1 | MT8342(58–78).1 | MT83428(0–2).1 | MT8342(84–93).1 | MT834(295–313).1 |
| MT8343(15–31).1 | MT83433(5–9).1 | MT834341.1 | MT83434(3–5).1 | MT83434(7,8).1 |
| MT834350.1 | MT8343(52–60).1 | MT83460(7–9).1 | MT834611.1 | MT83461(3–7).1 |
| MT8346(19,20).1 | MT83462(2–6).1 | MT834628.1 | MT83463(0–2).1 | MT8346(34–49).1 |
| MT83465(1–3).1 | MT83465(6,7).1 | MT83466(0–2).1 | MT834664.1 | MT83466(7,8).1 |
| MT83467(1–6).1 | MT8346(78–82).1 | MT8346(84–97).1 | MT834699.1 | MT834715.1 |
| MT83473(0,1).1 | MT83473(3–5).1 | MT8348(49,50).1 | MT83502(6,7).1 | MT835383.1 |
| MT8432(85–95).1 | MT843(297–301).1 | MT84330(3–7).1 | MT8433(09–20).1 | MT8437(37–59).1 |
| MT8440(21–49).1 | MT84408(8,9).1 | MT84587(7,8).1 | MT8460(04–15).1 | MT846413.1 |

(continued on next page)

(Table S1 continued)

| | | | | |
|---|---|---|---|---|
| MT846415.1 | MT84641(7,8).1 | MT84642(0–3).1 | MT84642(5,6).1 | MT8464(28–36).1 |
| MT8464(38–40).1 | MT84644(2–6).1 | MT8464(49–52).1 | MT84645(4–7).1 | MT84646(0–5).1 |
| MT8464(67–71).1 | MT846473.1 | MT84647(5–8).1 | MT84648(0,1).1 | MT84648(3–6).1 |
| MT8464(88–92).1 | MT846494.1 | MT846(499–515).1 | MT8465(17–23).1 | MT84652(5–9).1 |
| MT846531.1 | MT84653(3–5).1 | MT8465(37–41).1 | MT846545.1 | MT846547.1 |
| MT8465(53–61).1 | MT84656(3–5).1 | MT8465(67–81).1 | MT8465(83–98).1 | MT8466(00–16).1 |
| MT8466(18–23).1 | MT84662(6–9).1 | MT8466(31–55).1 | MT8466(57–60).1 | MT8466(62–71).1 |
| MT84667(3–7).1 | MT846679.1 | MT84668(1,2).1 | MT846684.1 | MT84668(6,7).1 |
| MT8466(89–91).1 | MT8472(08–28).1 | MT85645(2–9).1 | MT8566(78–80).1 | MT8566(82–91).1 |
| MT87249(2–9).1 | MT87250(1,2).1 | MT87305(0–3).1 | MT8730(55–73).1 | MT87307(5,6).1 |
| MT873(078–100).1 | MT87310(2–5).1 | MT8731(08–21).1 | MT8731(23–37).1 | MT8731(39–41).1 |
| MT873143.1 | MT8731(45–54).1 | MT8731(56–61).1 | MT873163.1 | MT87316(5–9).1 |
| MT8731(72–85).1 | MT8731(87–92).1 | MT873(194–213).1 | MT87321(6–8).1 | MT87322(0–6).1 |
| MT873228.1 | MT87323(0,1).1 | MT873233.1 | MT8732(35–40).1 | MT87324(2–5).1 |
| MT873247.1 | MT8732(49–51).1 | MT87325(3–9).1 | MT8732(61–70).1 | MT87327(2–5).1 |
| MT8732(77–87).1 | MT8732(89,90).1 | MT87329(2–6).1 | MT873(298–309).1 | MT87331(1,2).1 |
| MT873314.1 | MT8733(16–21).1 | MT87332(3,4).1 | MT8733(26–40).1 | MT8733(42–53).1 |
| MT87335(5,6).1 | MT8733(59–70).1 | MT87337(2,3).1 | MT8733(75–85).1 | MT873387.1 |
| MT8733(89–93).1 | MT87339(5–9).1 | MT8734(01–14).1 | MT8734(16–22).1 | MT873424.1 |
| MT873426.1 | MT8734(28–31).1 | MT87343(3,4).1 | MT8734(36–48).1 | MT8734(50–73).1 |
| MT87347(5–7).1 | MT8734(79–82).1 | MT8734(84–91).1 | MT87349(3,4).1 | MT873(496–501).1 |
| MT87350(3–7).1 | MT8735(09,10).1 | MT873801.1 | MT873803.1 | MT873807.1 |
| MT873812.1 | MT873821.1 | MT873827.1 | MT873890.1 | MT873892.1 |
| MT873898.1 | MT873923.1 | MT8753(54–60).1 | MT87643(1–3).1 | MT87652(5–7).1 |
| MT87654(6–8).1 | MT87655(4–6).1 | MT87657(1,2).1 | MT87659(8,9).1 | MT87660(6,7).1 |
| MT877256.1 | MT877296.1 | MT877(298–300).1 | MT877302.1 | MT87730(4–6).1 |
| MT877309.1 | MT87731(5,6).1 | MT877320.1 | MT877324.1 | MT8773(28–31).1 |
| MT877333.1 | MT877335.1 | MT877338.1 | MT877340.1 | MT877357.1 |

(continued on next page)

(Table S1 continued)

| | | | | |
|---|---|---|---|---|
| MT8773(59–61).1 | MT87736(5,6).1 | MT877379.1 | MT877381.1 | MT877383.1 |
| MT877409.1 | MT877418.1 | MT879619.1 | MT8808(73–88).1 | MT8823(10–25).1 |
| MT883(499–505).1 | MT8862(88–90).1 | MT886(292–318).1 | MT8863(21–60).1 | MT8863(62–73).1 |
| MT8863(75–86).1 | MT886407.1 | MT8864(09,10).1 | MT886413.1 | MT889692.1 |
| MT89021(0,1).1 | MT890230.1 | MT89023(5,6).1 | MT890238.1 | MT890240.1 |
| MT890243.1 | MT890277.1 | MT8902(79–81).1 | MT89028(5,6).1 | MT8902(88–90).1 |
| MT89029(2–6).1 | MT89030(1,2).1 | MT8903(09–11).1 | MT890313.1 | MT890316.1 |
| MT89032(0–2).1 | MT890330.1 | MT89033(5–8).1 | MT890340.1 | MT89034(4–9).1 |
| MT89035(2,3).1 | MT89035(5–8).1 | MT890462.1 | MT890669.1 | MT89(2963–3019).1 |
| MT893659.1 | MT89366(1,2).1 | MT893664.1 | MT8936(67–75).1 | MT893677.1 |
| MT8936(79–81).1 | MT89368(3–6).1 | MT893688.1 | MT893692.1 | MT89726(0,1).1 |
| MT8974(59–66).1 | MT9054(16–31).1 | MT90751(5–9).1 | MT907520.2 | MT911534.1 |
| MT9115(38–68).1 | MT911(796–801).1 | MT9118(03–25).1 | MT91182(7,8).1 | MT9130(38–59).1 |
| MT9130(61–71).1 | MT9130(73–90).1 | MT91309(2,3).1 | MT91309(5,6).1 | MT913(098–107).1 |
| MT9131(09–14).1 | MT91311(6–9).1 | MT9131(22–31).1 | MT91313(3–5).1 | MT9131(37–52).1 |
| MT9195(25–36).1 | MT9197(68–90).1 | MT9(19994–20040).1 | MT921009.1 | MT921159.1 |
| MT92116(5,6).1 | MT92598(0–9).1 | MT92599(1–7).1 | MT92896(0–8).1 | MT92897(0–6).1 |
| MT9289(79–85).1 | MT9289(87–92).1 | MT92899(4–6).1 | MT92(8998–9021).1 | MT9290(23–33).1 |
| MT9290(35–45).1 | MT929047.1 | MT9290(49–67).1 | MT9290(69–83).1 | MT929(085–101).1 |
| MT92910(4–7).1 | MT92911(0–2).1 | MT92911(5,6).1 | MT92911(8,9).1 | MT92912(3,4).1 |
| MT929128.1 | MT929130.1 | MT92913(4,5).1 | MT929148.1 | MT929150.1 |
| MT92915(3,4).1 | MT92915(6–9).1 | MT9373(17–26).1 | MT937808.1 | MT93781(0–8).1 |
| MT93782(0–7).1 | MT940449.1 | MT94045(1,2).1 | MT9404(58–62).1 | MT940464.1 |
| MT94047(2–9).1 | MT940(481–505).1 | MT9405(07–26).1 | MT941047.1 | MT941052.1 |
| MT94105(5,6).1 | MT9410(59–61).1 | MT941067.1 | MT941272.1 | MT941274.1 |
| MT941276.1 | MT941403.1 | MT9468(69–73).1 | MT9468(84–96).1 | MT9470(74–84).1 |
| MT9471(53–63).1 | MT9471(65–75).1 | MT9475(46–66).1 | MT947568.1 | MT9475(70–84).1 |
| MT94758(6–8).1 | MT94759(1–3).1 | MT947595.1 | MT9501(87–93).1 | MT950195.1 |

(continued on next page)

(Table S1 continued)

| | | | | |
|---|---|---|---|---|
| MT95023(0,1).1 | MT95034(8,9).1 | MT950532.1 | MT950684.1 | MT950725.1 |
| MT950761.1 | MT951157.1 | MT951170.1 | MT951177.1 | MT951181.1 |
| MT951208.1 | MT95139(2,3).1 | MT95192(4–7).1 | MT9519(29–32).1 | MT9519(34–42).1 |
| MT9519(44–53).1 | MT9519(55–78).1 | MT951980.1 | MT952134.1 | MT9526(03–10).1 |
| MT9526(12–40).1 | MT9526(42–56).1 | MT9526(63–73).1 | MT952675.1 | MT9526(77–85).1 |
| MT95268(7,8).1 | MT95269(0–8).1 | MT95270(0–2).1 | MT953868.1 | MT9538(74–80).1 |
| MT953887.1 | MT95389(0,1).1 | MT95389(3–5).1 | MT953925.1 | MT953978.1 |
| MT95398(0,1).1 | MT953984.1 | MT953991.1 | MT953996.1 | MT954060.1 |
| MT954065.1 | MT954067.1 | MT954069.1 | MT95407(1,2).1 | MT954103.1 |
| MT954414.1 | MT954416.1 | MT954593.1 | MT954959.1 | MT954971.1 |
| MT955114.1 | MT955161.1 | MT9551(68–74).1 | MT955326.1 | MT955347.1 |
| MT955354.1 | MT955360.1 | MT956642.1 | MT956684.1 | MT95668(6–8).1 |
| MT956693.1 | MT956695.1 | MT956(698–704).1 | MT956706.1 | MT9567(09,10).1 |
| MT9567(12–24).1 | MT95672(6,7).1 | MT9567(29–32).1 | MT95673(5–8).1 | MT9567(40–62).1 |
| MT9567(67–70).1 | MT95677(2,3).1 | MT95677(5–7).1 | MT9567(83–91).1 | MT95679(3–9).1 |
| MT95680(1–6).1 | MT95681(3,4).1 | MT95691(1–8).1 | MT9582(05–63).1 | MT965767.1 |
| MT965769.1 | MT96607(6,7).1 | MT966079.1 | MT966081.1 | MT96608(3,4).1 |
| MT96608(6,7).1 | MT966089.1 | MT966091.1 | MT96609(4–7).1 | MT966(099–102).1 |
| MT96610(4–8).1 | MT966110.1 | MT966113.1 | MT966126.1 | MT966129.1 |
| MT966133.1 | MT966138.1 | MT96614(0–2).1 | MT96614(4–6).1 | MT96614(8,9).1 |
| MT9661(54–65).1 | MT966167.1 | MT966169.1 | MT966171.1 | MT966174.1 |
| MT9661(76–81).1 | MT966183.1 | MT96618(5,6).1 | MT96619(1–3).1 | MT966195.1 |
| MT966197.1 | MT966199.1 | MT966201.1 | MT96620(4–6).1 | MT9662(09–14).1 |
| MT96621(6–9).1 | MT9662(23–31).1 | MT96623(3,4).1 | MT96623(7–9).1 | MT966241.1 |
| MT96624(3–9).1 | MT96625(1–7).1 | MT966262.1 | MT967(899–910).1 | MT9679(18–84).1 |
| MT96805(1–5).1 | MT9680(57–77).1 | MT969352.1 | MT96935(4,5).1 | MT96935(7–9).1 |
| MT969361.1 | MT96936(3,4).1 | MT9693(66–91).1 | MT969393.1 | MT969395.1 |
| MT969(397–405).1 | MT9694(08–19).1 | MT9694(21–34).1 | MT9694(36–41).1 | MT969443.1 |

(continued on next page)

(Table S1 continued)

| | | | | |
|---|---|---|---|---|
| MT9694(45–60).1 | MT9694(62–70).1 | MT969473.1 | MT9694(75–85).1 | MT969(488–526).1 |
| MT969528.1 | MT96953(0–4).1 | MT969(536–627).1 | MT9696(29–32).1 | MT96963(5–8).1 |
| MT9696(40–83).1 | MT969(685–721).1 | MT9697(23–34).1 | MT9697(36–97).1 | MT969(799–842).1 |
| MT96984(4,5).1 | MT9698(47–58).1 | MT96986(0–4).1 | MT96986(6,7).1 | MT9698(69–72).1 |
| MT969(874–903).1 | MT9699(05–79).1 | MT9(69981–70013).1 | MT9700(56–64).1 | MT970078.1 |
| MT97008(1,2).1 | MT970084.1 | MT970086.1 | MT97008(8,9).1 | MT970091.1 |
| MT970112.1 | MT9701(49–55).1 | MT970(158–265).1 | MT9702(67–91).1 | MT970(293–315).1 |
| MT97031(7,8).1 | MT97032(1–3).1 | MT970328.1 | MT9703(30–50).1 | MT97035(5–7).1 |
| MT9703(59,60).1 | MT970362.1 | MT970367.1 | MT97037(0–4).1 | MT970376.1 |
| MT9703(79,80).1 | MT970382.1 | MT97038(7,8).1 | MT970390.1 | MT970393.1 |
| MT970396.1 | MT970403.1 | MT970406.1 | MT970408.1 | MT97041(0–2).1 |
| MT970414.1 | MT970416.1 | MT9704(19,20).1 | MT97042(2,3).1 | MT970(439–591).1 |
| MT970(593–708).1 | MT970(710–804).1 | MT9708(06–66).1 | MT9708(68–76).1 | MT9708(78–89).1 |
| MT970(891–932).1 | MT9709(34–44).1 | MT9709(46–81).1 | MT97(0983–1002).1 | MT97100(4–8).1 |
| MT9710(10–71).1 | MT971(074–144).1 | MT9711(46–53).1 | MT971(155–235).1 | MT9712(37–52).1 |
| MT971(254–413).1 | MT97141(5,6).1 | MT9714(18–24).1 | MT9714(26–47).1 | MT9714(49–56).1 |
| MT9714(58–69).1 | MT97147(1–8).1 | MT9714(80–93).1 | MT971(495–503).1 | MT97150(5–8).1 |
| MT97151(0–2).1 | MT97151(4,5).1 | MT9715(17–33).1 | MT9715(35–51).1 | MT97155(3–8).1 |
| MT97156(0–2).1 | MT971564.1 | MT971566.1 | MT97156(8,9).1 | MT97157(2–4).1 |
| MT97157(6,7).1 | MT971579.1 | MT97158(1–6).1 | MT9715(88–91).1 | MT97159(4–6).1 |
| MT971(599,600).1 | MT9716(02–13).1 | MT97161(5–9).1 | MT97162(1,2).1 | MT971627.1 |
| MT9716(29–32).1 | MT9716(34–47).1 | MT9716(49–67).1 | MT97167(0–3).1 | MT9716(75–89).1 |
| MT97169(1–8).1 | MT97170(0–7).1 | MT9717(09–16).1 | MT9717(18–31).1 | MT971(733–816).1 |
| MT9718(18–24).1 | MT9718(26–48).1 | MT9718(51–68).1 | MT9718(70–85).1 | MT971(887–903).1 |
| MT9719(05–10).1 | MT9719(12–21).1 | MT97192(5–7).1 | MT9719(30–40).1 | MT97194(2,3).1 |
| MT9719(45–54).1 | MT9719(57–79).1 | MT97(1981–2020).1 | MT97202(2–4).1 | MT9720(26–42).1 |
| MT9720(44–54).1 | MT9720(56–73).1 | MT9720(75–81).1 | MT972083.1 | MT9720(85–96).1 |
| MT972(098–100).1 | MT9721(11–23).1 | MT972125.1 | MT9721(28–33).1 | MT97213(5–7).1 |

(continued on next page)

(Table S1 continued)

| | | | | |
|---|---|---|---|---|
| MT9721(39–45).1 | MT9721(47–53).1 | MT972155.1 | MT9721(58–66).1 | MT972168.1 |
| MT97217(1–8).1 | MT97218(1,2).1 | MT97218(4–9).1 | MT972(191–207).1 | MT9722(09–17).1 |
| MT97222(0,1).1 | MT97222(3–8).1 | MT97223(1–9).1 | MT9722(41–53).1 | MT972(255–308).1 |
| MT9723(10–20).1 | MT9723(22–33).1 | MT97233(5–8).1 | MT9723(40–53).1 | MT972355.1 |
| MT9723(57–95).1 | MT972(397–400).1 | MT9724(02–18).1 | MT97242(0,1).1 | MT9724(23–38).1 |
| MT97244(0–3).1 | MT9724(45–98).1 | MT972500.1 | MT9725(02–30).1 | MT9725(32–44).1 |
| MT972546.1 | MT9725(48–68).1 | MT97257(0,1).1 | MT97257(3–7).1 | MT972580.1 |
| MT972582.1 | MT9725(84–92).1 | MT97259(4–6).1 | MT972599.1 | MT972602.1 |
| MT97260(4,5).1 | MT9726(07–12).1 | MT97261(4–6).1 | MT97261(8,9).1 | MT97262(1,2).1 |
| MT97262(5–7).1 | MT9726(29–32).1 | MT9726(34–40).1 | MT97264(2,3).1 | MT97264(7–9).1 |
| MT97265(1–9).1 | MT972662.1 | MT9726(64–72).1 | MT972675.1 | MT9726(77–81).1 |
| MT97268(3,4).1 | MT9726(86–92).1 | MT972695.1 | MT972697.1 | MT9727(00–23).1 |
| MT972725.1 | MT9727(27–30).1 | MT97273(2–4).1 | MT9727(36–46).1 | MT9727(48–50).1 |
| MT97275(2–4).1 | MT972756.1 | MT972758.1 | MT97276(0–5).1 | MT9727(67–90).1 |
| MT97279(2–6).1 | MT972(799–814).1 | MT972816.1 | MT9728(18–51).1 | MT97285(3,4).1 |
| MT9728(56–74).1 | MT97287(6–9).1 | MT972881.1 | MT9728(83–96).1 | MT972(898–904).1 |
| MT9729(06–13).1 | MT97291(5–7).1 | MT9729(19–40).1 | MT972942.1 | MT972944.1 |
| MT97294(6–9).1 | MT97295(1,2).1 | MT97295(4,5).1 | MT972957.1 | MT9729(59,60).1 |
| MT972962.1 | MT97296(4–7).1 | MT9729(69,70).1 | MT97297(3–8).1 | MT97298(0,1).1 |
| MT97298(4–6).1 | MT97298(8,9).1 | MT97299(1–8).1 | MT97300(0–6).1 | MT97300(8,9).1 |
| MT973011.1 | MT9730(13–20).1 | MT9730(22–35).1 | MT9730(37–46).1 | MT9730(48–57).1 |
| MT973059.1 | MT97306(1–3).1 | MT9730(65–75).1 | MT973077.1 | MT97308(0–6).1 |
| MT9730(88–93).1 | MT97309(5–8).1 | MT97310(0,1).1 | MT9731(03–21).1 | MT973123.1 |
| MT9731(25–30).1 | MT97313(2–7).1 | MT973139.1 | MT97314(1–8).1 | MT9731(50–87).1 |
| MT973(189–214).1 | MT97321(6–8).1 | MT97322(0–2).1 | MT97322(5–8).1 | MT97323(0–7).1 |
| MT973239.1 | MT97324(1,2).1 | MT97324(4–7).1 | MT9732(49–60).1 | MT97326(2–7).1 |
| MT9732(69–76).1 | MT9732(78–86).1 | MT9732(88–92).1 | MT973(294–302).1 | MT97330(4–8).1 |
| MT9733(10–24).1 | MT9733(26–39).1 | MT97334(1,2).1 | MT973344.1 | MT97334(7,8).1 |

(continued on next page)

(Table S1 continued)

| | | | | |
|---|---|---|---|---|
| MT973353.1 | MT973356.1 | MT973358.1 | MT973366.1 | MT973371.1 |
| MT973373.1 | MT973376.1 | MT973378.1 | MT97338(0,1).1 | MT973389.1 |
| MT973391.1 | MT97339(5,6).1 | MT973402.1 | MT973409.1 | MT973419.1 |
| MT973421.1 | MT973424.1 | MT97342(6,7).1 | MT973431.1 | MT97343(5,6).1 |
| MT973438.1 | MT973448.1 | MT973454.1 | MT973484.1 | MT974069.1 |
| MT974071.1 | MT981(086–108).1 | MT9814(18–41).1 | MT9814(49–57).1 | MT9814(59–67).1 |
| MT981483.1 | MT981485.1 | MT9814(87–90).1 | MT9904(49,50).1 | MT9927(23–51).1 |
| MT993864.1 | MT993866.1 | MT993868.1 | MT9938(70–87).1 | MT994286.1 |
| MT99439(3–5).1 | MT994632.1 | MT994849.1 | MT994881.1 | MT994884.1 |
| MT9949(01–89).1 | MT99720(3–6).1 | MT9972(08–10).1 | MT99721(2–6).1 | MT9972(18–30).1 |
| MT99723(2,3).1 | MT99723(5–7).1 | MT99724(0–3).1 | MT997(685–747).1 | MT9977(62–71).1 |
| MT9979(49–94).1 | MT99844(2–4).1 | MW0003(54–61).1 | MW00036(3–6).1 | MW000369.1 |
| MW000371.1 | MW0012(32–88).1 | MW0016(67–75).1 | MW00169(2,3).1 | MW001(880–931).1 |
| MW00280(6–9).1 | MW00281(1–3).1 | MW00281(5–8).1 | MW00282(2,3).1 | MW00282(5–9).1 |
| MW002831.1 | MW00283(3,4).1 | MW00283(6,7).1 | MW004168.1 | MW00655(5–7).1 |
| MW0065(59–61).1 | MW00656(3–5).1 | MW006567.1 | MW006569.1 | MW006571.1 |
| MW00657(3–9).1 | MW006583.1 | MW00658(5,6).1 | MW00659(0–6).1 | MW006(598–601).1 |
| MW00660(3–7).1 | MW008562.1 | MW0085(65–71).1 | MW0085(73–85).1 | MW008587.1 |
| MW008590.1 | MW009055.1 | MW009107.1 | MW009255.1 | MW01002(8,9).1 |
| MW01023(5,6).1 | MW0102(45–55).1 | MW01176(2–8).1 | MW0122(66–79).1 | MW014808.1 |
| MW018448.1 | MW0200(61–86).1 | MW0214(54–61).1 | MW02146(3–6).1 | MW0214(68–70).1 |
| MW02151(2–4).1 | MW0233(88–94).1 | MW023404.1 | MW023409.1 | MW02341(5,6).1 |
| MW023418.1 | MW023424.1 | MW02342(8,9).1 | MW02343(5–7).1 | MW0234(39,40).1 |
| MW02344(2,3).1 | MW0234(46–50).1 | MW02345(2,3).1 | MW02345(6,7).1 | MW0234(59–62).1 |
| MW023466.1 | MW02347(0,1).1 | MW023474.1 | MW023476.1 | MW023478.1 |
| MW023480.1 | MW02348(2,3).1 | MW02348(6,7).1 | MW02349(4,5).1 | MW023498.1 |
| MW02350(0–2).1 | MW023511.1 | MW02351(6,7).1 | MW02352(2,3).1 | MW023525.1 |
| MW023532.1 | MW023542.1 | MW023547.1 | MW023550.1 | MW023552.1 |

(continued on next page)

(Table S1 continued)

| | | | | |
|---|---|---|---|---|
| MW023555.1 | MW023558.1 | MW023562.1 | MW023564.1 | MW023570.1 |
| MW023576.1 | MW0235(79,80).1 | MW023584.1 | MW023592.1 | MW023935.1 |
| MW030(193–278).1 | MW030985.1 | MW03098(8,9).1 | MW03099(3,4).1 | MW030996.1 |
| MW030998.1 | MW03100(0,1).1 | MW03100(3,4).1 | MW0310(09–11).1 | MW03101(4–8).1 |
| MW03102(1–3).1 | MW031025.1 | MW0310(27–30).1 | MW03103(3–5).1 | MW0310(37–40).1 |
| MW0310(43–56).1 | MW0310(58–61).1 | MW0310(63–74).1 | MW0310(77–80).1 | MW031082.1 |
| MW031084.1 | MW031086.1 | MW0310(88–93).1 | MW031(799–803).1 | MW03537(0,1).1 |
| MW03537(3,4).1 | MW0353(76–82).1 | MW03538(4–6).1 | MW03538(8,9).1 | MW03539(1–7).1 |
| MW035(399–416).1 | MW0354(18–20).1 | MW03542(2,3).1 | MW0354(25–56).1 | MW0354(58–66).1 |
| MW03546(8,9).1 | MW03547(1–6).1 | MW035484.1 | MW035(495–507).1 | MW035509.1 |
| MW03551(1,2).1 | MW0355(14–22).1 | MW0355(24–43).1 | MW03554(5,6).1 | MW0355(48–62).1 |
| MW0355(64–79).1 | MW0359(18–60).1 | MW03596(2–8).1 | MW03597(0–4).1 | MW035976.1 |
| MW035984.1 | MW03598(7,8).1 | MW035990.1 | MW035992.1 | MW035997.1 |
| MW03(5999,6000).1 | MW036003.1 | MW03600(6–8).1 | MW03601(0,1).1 | MW03601(3,4).1 |
| MW0360(16–24).1 | MW03602(6–9).1 | MW03603(1,2).1 | MW03603(4,5).1 | MW03603(8,9).1 |
| MW03604(1–4).1 | MW03604(6,7).1 | MW0360(49–51).1 | MW03605(3–7).1 | MW036060.1 |
| MW03606(2,3).1 | MW0360(67–74).1 | MW03608(0,1).1 | MW03608(3,4).1 | MW036086.1 |
| MW03923(4,5).1 | MW039450.1 | MW040606.1 | MW040617.1 | MW04062(1–5).1 |
| MW0406(27–36).1 | MW0406(38–55).1 | MW0406(59,60).1 | MW04066(4–9).1 | MW040673.1 |
| MW040675.1 | MW04067(8,9).1 | MW040681.1 | MW040683.1 | MW04068(6,7).1 |
| MW040690.1 | MW04069(3–7).1 | MW041156.1 | MW042923.1 | MW042925.1 |
| MW04701(6–8).1 | MW047085.1 | MW0473(07–18).1 | MW04(8997–9003).1 | MW0490(05–23).1 |
| MW052620.1 | MW05262(4,5).1 | MW052629.1 | MW0526(31–93).1 | MW053(297–319).1 |
| MW053708.1 | MW053750.1 | MW053767.1 | MW053869.1 | MW05387(4–6).1 |
| MW053883.1 | MW053885.1 | MW053921.1 | MW053925.1 | MW053929.1 |
| MW053935.1 | MW053938.1 | MW053950.1 | MW053957.1 | MW053961.1 |
| MW053976.1 | MW05398(0,1).1 | MW054091.1 | MW054103.1 | MW05410(8,9).1 |
| MW054137.1 | MW054154.1 | MW05603(2–4).1 | MW056078.1 | MW056086.1 |

(continued on next page)

(Table S1 continued)

| | | | | |
|---|---|---|---|---|
| MW05609(2,3).1 | MW056097.1 | MW05610(1,2).1 | MW05610(4,5).1 | MW056107.1 |
| MW0561(09–12).1 | MW05611(4–8).1 | MW056122.1 | MW05612(4–8).1 | MW056131.1 |
| MW05613(5–8).1 | MW056141.1 | MW05614(6,7).1 | MW056149.1 | MW05615(1,2).1 |
| MW05615(5,6).1 | MW056159.1 | MW056161.1 | MW0561(66–77).1 | MW0643(14–29).1 |
| MW064331.1 | MW0643(33–83).1 | MW064(385–430).1 | MW0644(32–70).1 | MW0644(72–94).1 |
| MW064(496–535).1 | MW06453(7–9).1 | MW06454(1–9).1 | MW064551.1 | MW06455(3–8).1 |
| MW0645(60–79).1 | MW0645(81–94).1 | MW064(597–681).1 | MW06468(3–5).1 | MW0646(87–92).1 |
| MW06469(4–7).1 | MW064699.1 | MW06470(1–5).1 | MW0647(07–17).1 | MW064719.1 |
| MW0647(21–30).1 | MW0647(32–67).1 | MW0647(69–72).1 | MW0647(74–80).1 | MW064782.1 |
| MW064(784–810).1 | MW0648(12–21).1 | MW0648(23–32).1 | MW06483(4,5).1 | MW0648(37–54).1 |
| MW0648(56–60).1 | MW0648(62–86).1 | MW064(888–938).1 | MW0649(40–82).1 | MW06498(4–7).1 |
| MW0649(89,90).1 | MW06(4992–5044).1 | MW0650(46–74).1 | MW065(076–101).1 | MW06510(3,4).1 |
| MW0651(06–30).1 | MW0651(32–44).1 | MW0651(46–56).1 | MW065158.1 | MW0651(60–98).1 |
| MW06520(0,1).1 | MW06520(3–8).1 | MW06521(0,1).1 | MW0652(13–24).1 | MW0652(26–40).1 |
| MW06524(2–5).1 | MW0652(47–73).1 | MW065(275–310).1 | MW0653(12–27).1 | MW0653(29–55).1 |
| MW065(357–422).1 | MW0654(24–83).1 | MW0676(75–82).1 | MW067(684–749).1 | MW06775(1–3).1 |
| MW0677(57–66).1 | MW0677(68–71).1 | MW06777(3–5).1 | MW067(777–800).1 | MW06780(2–6).1 |
| MW0678(09–18).1 | MW06782(0–3).1 | MW06782(5–7).1 | MW070033.1 | MW0700(35–46).1 |
| MW0700(48–51).1 | MW070054.1 | MW070057.1 | MW0700(59–82).1 | MW0700(84–93).1 |
| MW07009(6–9).1 | MW07010(1–3).1 | MW0701(05–13).1 | MW07573(2,3).1 | MW075735.1 |
| MW0757(37–42).1 | MW07574(4,5).1 | MW07574(7–9).1 | MW07575(1–8).1 | MW07576(0–8).1 |
| MW07577(7–9).1 | MW0757(81–91).1 | MW07579(3–9).1 | MW07580(1–4).1 | MW075808.1 |
| MW0774(66–83).1 | MW07748(5–7).1 | MW077489.1 | MW07749(1–8).1 | MW07750(0–2).1 |
| MW07941(8,9).1 | MW07942(1,2).1 | MW07942(8,9).1 | MW0798(19–42).1 | NC_045512.2 |

Table S2: List of SARS-CoV-1 genome accession codes.

| | | |
|---|---|---|
| AY278487.3 | AY304486.1 | AY338174.1 |
| AY348314.1 | AY394850.2 | AY572034.1 |
| AY772062.1 | DQ412042.1 | FJ882938.1 |
| FJ882954.1 | | |

Table S3: List of coronavirus (non-SARS) genome accession codes.

| | | | | |
|---|---|---|---|---|
| AF304460.1 | AY278487.3 | AY304486.1 | AY338174.1 | AY348314.1 |
| AY391777.1 | AY394850.2 | AY567487.2 | AY572034.1 | AY58522(8,9).1 |
| AY597011.2 | AY772062.1 | AY884001.1 | AY9034(59,60).1 | DJ009246.1 |
| DQ339101.1 | DQ412042.1 | DQ415(896–914).1 | DQ44591(1,2).1 | FJ415324.1 |
| FJ882938.1 | FJ882954.1 | HM034837.1 | JN12983(4,5).1 | JQ7655(63–75).1 |
| JX104161.1 | JX50306(0,1).1 | JX504050.1 | JX524171.1 | KF430201.1 |
| KF514430.1 | KF51443(2,3).1 | KF53006(0,1).1 | KF5300(63–92).1 | KF53009(4–9).1 |
| KF5301(04–14).1 | KF68634(0–4).1 | KF686346.1 | KF745068.1 | KF917527.1 |
| KF923(886–925).1 | KF958702.1 | KJ36150(0,1).1 | KJ477102.1 | KJ556336.1 |
| KJ95821(8,9).1 | KP19861(0,1).1 | KP71992(7,8).1 | KT121572.1 | KT266906.1 |
| KT381875.1 | KT77955(5,6).1 | KU131570.1 | KU291448.1 | KU521535.1 |
| KX179500.1 | KX344031.1 | KX5389(64–79).1 | KY01428(1,2).1 | KY36990(5–8).1 |
| KY369909.2 | KY36991(0–4).1 | KY5549(67–75).1 | KY621348.1 | KY6749(14–21).1 |
| KY67494(1–3).1 | KY6847(59,60).1 | KY829118.1 | KY9673(56–61).1 | KY983583.1 |
| KY98358(5–8).1 | KY996417.1 | LG07610(3–5).1 | MF314143.1 | MF374983.2 |
| MF374984.1 | MF374985.2 | MF542265.1 | MG1977(09–23).1 | MG428699.1 |
| MG42870(1–6).1 | MG772808.1 | MG97744(4,5).1 | MG977447.1 | MG977449.1 |
| MG97745(1,2).1 | MH121121.1 | MH940245.1 | MK167038.1 | MK3036(19–25).1 |
| MK33404(3–7).1 | MN026164.1 | MN306018.1 | MN306036.1 | MN30604(0–3).1 |
| MN306046.1 | MN306053.1 | MN310476.1 | MN310478.1 | MN369046.1 |
| MT438(696–700).1 | NC_002645.1 | NC_005831.2 | NC_006213.1 | NC_006577.2 |

Table S4: Summary of primers and probes for each set. We only considered new sequences from each publication, that is, those which to our knowledge have not been reported previously.

| Name of Set | Primer/probe | Sequence $5' \rightarrow 3'$ |
|---|---|---|
| Bhadra et al. [1] | E-PF | TCGGAAGAGACAGGTACGTT |
| Bhadra et al. [1] | E-Probe | CATCCTTACTGCGCTTCGAT |
| Bhadra et al. [1] | E-RP | CAATATTGCAGCAGTACGCA |
| Bhadra et al. [1] | M-PF | CTGACCAGACCGCTTCTAG |
| Bhadra et al. [1] | M-Probe | GACCTGCCTAAAGAAATCACT |
| Bhadra et al. [1] | M-RP | GAAAGCGTTCGTGATGTAGC |
| CDC | 2019-nCoV_N1-F | GACCCCAAAATCAGCGAAAT |
| CDC | 2019-nCoV_N1-P | ACCCCGCATTACGTTTGGTGGACC |
| CDC | 2019-nCoV_N1-R | TCTGGTTACTGCCAGTTGAATCTG |
| CDC | 2019-nCoV_N2-F | TTACAAACATTGGCCGCAAA |
| CDC | 2019-nCoV_N2-P | ACAATTTGCCCCCAGCGCTTCAG |
| CDC | 2019-nCoV_N2-R | GCGCGACATTCCGAAGAA |
| Corman et al. [2] | E_Sarbeco_F | ACAGGTACGTTAATAGTTAATAGCGT |
| Corman et al. [2] | E_Sarbeco_P1 | ACACTAGCCATCCTTACTGCGCTTCG |
| Corman et al. [2] | E_Sarbeco_R | ATATTGCAGCAGTACGCACACA |
| Corman et al. [2] | N_Sarbeco_F | CACATTGGCACCCGCAATC |
| Corman et al. [2] | N_Sarbeco_P | ACTTCCTCAAGGAACAACATTGCCA |
| Corman et al. [2] | N_Sarbeco_R | GAGGAACGAGAAGAGGCTTG |

(continued on next page)

(Table S4 continued)

| Name of Set | Primer/probe | Sequence 5′ → 3′ |
|---|---|---|
| Corman et al. [2] | RdRp_SARSr-FA | GTGAAATGGTCATGTGTGGCGG |
| Corman et al. [2] | RdRp_SARSr-FG | GTGAGATGGTCATGTGTGGCGG |
| Corman et al. [2] | RdRp_SARSr-P1-1 | CCAGGTGGAACGTCATCAGGTGATGC |
| Corman et al. [2] | RdRp_SARSr-P1-2 | CCAGGTGGAACGTCATCCGGTGATGC |
| Corman et al. [2] | RdRp_SARSr-P1-3 | CCAGGTGGAACATCATCAGGTGATGC |
| Corman et al. [2] | RdRp_SARSr-P1-4 | CCAGGTGGAACATCATCCGGTGATGC |
| Corman et al. [2] | RdRp_SARSr-P1-5 | CCAGGTGGTACGTCATCAGGTGATGC |
| Corman et al. [2] | RdRp_SARSr-P1-6 | CCAGGTGGTACGTCATCCGGTGATGC |
| Corman et al. [2] | RdRp_SARSr-P1-7 | CCAGGTGGTACATCATCAGGTGATGC |
| Corman et al. [2] | RdRp_SARSr-P1-8 | CCAGGTGGTACATCATCCGGTGATGC |
| Corman et al. [2] | RdRp_SARSr-P2 | CAGGTGGAACCTCATCAGGAGATGC |
| Corman et al. [2] | RdRp_SARSr-R1 | CAAATGTTAAACACACTATTAGCATA |
| Corman et al. [2] | RdRp_SARSr-R2 | CAAATGTTAAAGACACTATTAGCATA |
| Corman et al. [2] | RdRp_SARSr-R3 | CAGATGTTAAACACACTATTAGCATA |
| Corman et al. [2] | RdRp_SARSr-R4 | CAGATGTTAAAGACACTATTAGCATA |
| Davda et al. [3] | MF1 | TGGCAGATTCCAACGG |
| Davda et al. [3] | MF2 | CCTTGAACAATGGAACCTAG |
| Davda et al. [3] | MR1 | ACAGCGTCCTAGATGGTG |
| Davda et al. [3] | MR2 | AGCAATACGAAGATGTCCAC |
| Davda et al. [3] | N1F1 | AGGAAATTTTGGGGACCAGGAA |
| Davda et al. [3] | N1F2 | GATTACAAACATTGGCCGCAAATTG |

(continued on next page)

(Table S4 continued)

| Name of Set | Primer/probe | Sequence 5′ → 3′ |
|---|---|---|
| Davda et al. [3] | N1R1 | CTGCTCATGGATTGTTGCAA |
| Davda et al. [3] | N1R2 | GAAATCATCCAAATCTGCAGCAG |
| Davda et al. [3] | N2F1 | GCCTCGGCAAAAACGTAC |
| Davda et al. [3] | N2F2 | GTAACACAAGCTTTCGGC |
| Davda et al. [3] | N2R1 | CTTGTGTGGTCTGCATGAG |
| Davda et al. [3] | N2R2 | TGAGTTGAGTCAGCACTG |
| Davda et al. [3] | Orf1abF1 | GGGAAATTGTTAAATTTATCTCAACCTG |
| Davda et al. [3] | Orf1abF2 | AATTGTCGGTGGACAAATTGTC |
| Davda et al. [3] | Orf1abR1 | CTGGTGTACCAACCAATGGAGC |
| Davda et al. [3] | Orf1abR2 | ACTAGTAGGTTGTTCTAATGGTTG |
| Grant et al. [4] | NgeneProbe | CTGTCACTAAGAAATCTGCTGCTGAGGC |
| Grant et al. [4] | NgeneTaq1 | TCTGGTAAAGGCCAACAACAA |
| Grant et al. [4] | NgeneTaq2 | TGTATGCTTTAGTGGCAGTACG |
| Hirotsu et al. [5] | NIID-N2-F | AAATTTTGGGGACCAGGAAC |
| Hirotsu et al. [5] | NIID-N2-P | ATGTCGCGCATTGGCATGGA |
| Hirotsu et al. [5] | NIID-N2-R | TGGCAGCTGTGTAGGTCAAC |
| Lanza et al. [6] | UFRN_1_F | GGGCATACACTCGCTATGTC |
| Lanza et al. [6] | UFRN_1_P | TCTGTGGCCCTGATGGCTACCCT |
| Lanza et al. [6] | UFRN_1_R | GCATGAAGCTTTACCAGCAC |
| Lanza et al. [6] | UFRN_2_F | GGCTACTAACAATGCCATGC |
| Lanza et al. [6] | UFRN_2_P | GGGTGGTAGTTGTGTTTTAAGCGG |

(continued on next page)

(Table S4 continued)

| Name of Set | Primer/probe | Sequence 5′ → 3′ |
|---|---|---|
| Lanza et al. [6] | UFRN_2_R | TAACATTTGGGCCGACAACA |
| Lanza et al. [6] | UFRN_3_F | TTCATGTTGTCGGCCCAAAT |
| Lanza et al. [6] | UFRN_3_P | GAAGACATTCAACTTCTTAAGAGTGC |
| Lanza et al. [6] | UFRN_3_R | TGGTGCAAGTAGAACTTCGT |
| Lanza et al. [6] | UFRN_4_F | TGGTGCTAGGAGAGTGTGG |
| Lanza et al. [6] | UFRN_4_P | CTTATGAATGTCTTGACACTCGTTTATA |
| Lanza et al. [6] | UFRN_4_R | CCCACATGGAAATGGCTTGAT |
| Lanza et al. [6] | UFRN_5_F | AGGGCACACTAGAACCAGAA |
| Lanza et al. [6] | UFRN_5_P | GGTCCAGACATGTTCCTCGGAACT |
| Lanza et al. [6] | UFRN_5_R | CAATTTCAGCAGGACAACGC |
| Lanza et al. [6] | UFRN_6_F | TCTTCACGACATTGGTAACCC |
| Lanza et al. [6] | UFRN_6_P | TACCTCAAGCTGATGTAGAATGGAAG |
| Lanza et al. [6] | UFRN_6_R | TCACTACAAGGCTGTGCATC |
| Lanza et al. [6] | UFRN_7_F | CTTCACGACATTGGTAACCCT |
| Lanza et al. [6] | UFRN_7_P | GTGTACCTCAAGCTGATGTAGAATGG |
| Lanza et al. [6] | UFRN_7_R | GTCACTACAAGGCTGTGCAT |
| Lanza et al. [6] | UFRN_8_F | GGCACAGGTGTTCTTACTGA |
| Lanza et al. [6] | UFRN_8_P | CCAACAATTTGGCAGAGACATTGC |
| Lanza et al. [6] | UFRN_8_R | TCAAGTGTCTGTGGATCACG |
| Lanza et al. [6] | UFRN_9_F | AGGCACAGGTGTTCTTACTG |
| Lanza et al. [6] | UFRN_9_P | TCCAACAATTTGGCAGAGACATTGC |

(continued on next page)

(Table S4 continued)

| Name of Set | Primer/probe | Sequence 5′ → 3′ |
|---|---|---|
| Lanza et al. [6] | UFRN_9_R | TCACGGACAGCATCAGTAGT |
| Li et al. [7] | S-1 | TGTACTTGGACAATCAAAAAGAGTTGAT |
| Li et al. [7] | S-2 | AGGAGCAGTTGTGAAGTTCTTTTC |
| Lu et al. [8] | ORF1a-F | AGAAGATTGGTTAGATGATGATAGT |
| Lu et al. [8] | ORF1a-P | TCCTCACTGCCGTCTTGTTGACCA |
| Lu et al. [8] | ORF1a-R | TTCCATCTCTAATTGAGGTTGAACC |
| Luminex | China_N_F | GGGGAACTTCTCCTGCTAGAAT |
| Luminex | China_N_P | TTGCTGCTGCTTGACAGATT |
| Luminex | China_N_R | CAGACATTTTGCTCTCAAGCTG |
| Luminex | China_ORF_F | CCCTGTGGGTTTTACACTTAA |
| Luminex | China_ORF_P | CCGTCTGCGGTATGTGGAAAGGTTATGG |
| Luminex | China_ORF_R | ACGATTGTGCATCAGCTGA |
| Munnink et al. [9] | SARS-CoV-2_1_LEFT | ACCAACCAACTTTCGATCTCTTGT |
| Munnink et al. [9] | SARS-CoV-2_1_RIGHT | CGAGCATCCGAACGTTTGATGA |
| Munnink et al. [9] | SARS-CoV-2_2_LEFT | TCGTACGTGGCTTTGGAGACTC |
| Munnink et al. [9] | SARS-CoV-2_2_RIGHT | ATGCACTCAAGAGGGTAGCCAT |
| Munnink et al. [9] | SARS-CoV-2_3_LEFT | ACGAGCTTGGCACTGATCCTTA |
| Munnink et al. [9] | SARS-CoV-2_3_RIGHT | GGTTGCATTCATTTGGTGACGC |
| Munnink et al. [9] | SARS-CoV-2_4_LEFT | ACACCTTCAATGGGGAATGTCC |
| Munnink et al. [9] | SARS-CoV-2_4_RIGHT | AGGCACACTTGTTATGGCAACC |
| Munnink et al. [9] | SARS-CoV-2_5_LEFT | TGTCCAGCATGTCACAATTCAGA |

(continued on next page)

(Table S4 continued)

| Name of Set | Primer/probe | Sequence 5′ → 3′ |
|---|---|---|
| Munnink et al. [9] | SARS-CoV-2_5_RIGHT | TACAACACGAGCAGCCTCTGAT |
| Munnink et al. [9] | SARS-CoV-2_6_LEFT | TGTGAAAGGTTTGGATTATAAAGCATTCA |
| Munnink et al. [9] | SARS-CoV-2_6_RIGHT | ACAGGTGACAATTTGTCCACCG |
| Munnink et al. [9] | SARS-CoV-2_7_LEFT | TGGCACTGTTTATGAAAAACTCAAACC |
| Munnink et al. [9] | SARS-CoV-2_7_RIGHT | TTTCGAGCAACATAAGCCCGTT |
| Munnink et al. [9] | SARS-CoV-2_8_LEFT | AGGGAGAAACACTTCCCACAGA |
| Munnink et al. [9] | SARS-CoV-2_8_RIGHT | AATCAATGCCCAGTGGTGTAAGT |
| Munnink et al. [9] | SARS-CoV-2_9_LEFT | TCACTTTTGAACTTGATGAAAGGATTGA |
| Munnink et al. [9] | SARS-CoV-2_9_RIGHT | GATTGTCCTCACTGCCGTCTTG |
| Munnink et al. [9] | SARS-CoV-2_11_LEFT | AAACATGGAGGAGGTGTTGCAG |
| Munnink et al. [9] | SARS-CoV-2_11_RIGHT | TCTTGTTTTCTCTGTTCAACTGAAGGT |
| Munnink et al. [9] | SARS-CoV-2_12_RIGHT | CCTGACCCGGGTAAGTGGTTAT |
| Munnink et al. [9] | SARS-CoV-2_13_LEFT | GCTCCATATATAGTGGGTGATGTTGT |
| Munnink et al. [9] | SARS-CoV-2_13_RIGHT | AGCCATGTGTTACATAGCCAAGTG |
| Munnink et al. [9] | SARS-CoV-2_14_LEFT | GTTTCAACTATACAGCGTAAATATAAGGGT |
| Munnink et al. [9] | SARS-CoV-2_14_RIGHT | CGTGTGGAGGTTAATGTTGTCTACT |
| Munnink et al. [9] | SARS-CoV-2_15_LEFT | CTATTCTGGACAATCTACACAACTAGGT |
| Munnink et al. [9] | SARS-CoV-2_15_RIGHT | AGCATCTTGTAGAGCAGGTGGA |
| Munnink et al. [9] | SARS-CoV-2_16_LEFT | AGGTACATGTCAGCATTAAATCACACT |
| Munnink et al. [9] | SARS-CoV-2_16_RIGHT | AGTTCATACTGAGCAGGTGGTG |
| Munnink et al. [9] | SARS-CoV-2_17_LEFT | GCTGTTATGTACATGGGCACACT |

(continued on next page)

(Table S4 continued)

| Name of Set | Primer/probe | Sequence 5′ → 3′ |
|---|---|---|
| Munnink et al. [9] | SARS-CoV-2_17_RIGHT | GCTTGCGTTTGGATATGGTTGG |
| Munnink et al. [9] | SARS-CoV-2_18_LEFT | CAGTTACACAACAACCATAAAACCAGT |
| Munnink et al. [9] | SARS-CoV-2_18_RIGHT | GATTATCCATTCCCTGCGCGTC |
| Munnink et al. [9] | SARS-CoV-2_19_LEFT | TGTTACATAAACCTATTGTTTGGCATGT |
| Munnink et al. [9] | SARS-CoV-2_19_RIGHT | TCCCAAGGGACACTATTAACAGCA |
| Munnink et al. [9] | SARS-CoV-2_20_LEFT | TACAGAAGAGGTTGGCCACACA |
| Munnink et al. [9] | SARS-CoV-2_21_LEFT | AGCAAAGAATACTGTTAAGAGTGTCGG |
| Munnink et al. [9] | SARS-CoV-2_21_RIGHT | TCGGGGCCATTTGTACAAGATT |
| Munnink et al. [9] | SARS-CoV-2_22_LEFT | AGTTGCAGAGTGGTTTTTGGCA |
| Munnink et al. [9] | SARS-CoV-2_22_RIGHT | ACTGTAGTGACAAGTCTCTCGCA |
| Munnink et al. [9] | SARS-CoV-2_23_LEFT | ACTATTGTTAATGGTGTTAGAAGGTCCT |
| Munnink et al. [9] | SARS-CoV-2_23_RIGHT | GCAACTTCCGCACTATCACCAA |
| Munnink et al. [9] | SARS-CoV-2_24_LEFT | AGCTAATAACACTAAAGGTTCATTGCCT |
| Munnink et al. [9] | SARS-CoV-2_24_RIGHT | TGACTTTTTGCTACCTGCGCAT |
| Munnink et al. [9] | SARS-CoV-2_25_LEFT | TGTCTTAAATTGTCACATCAATCTGACAT |
| Munnink et al. [9] | SARS-CoV-2_25_RIGHT | TCACGAGTGACACCACCATCAA |
| Munnink et al. [9] | SARS-CoV-2_26_LEFT | TGGTTGAAGCAGTTAATTAAAGTTACACT |
| Munnink et al. [9] | SARS-CoV-2_26_RIGHT | TTCAGCAGCCAAAACACAAGCT |
| Munnink et al. [9] | SARS-CoV-2_27_LEFT | CTGGTTTGCCTGGCACGATATT |
| Munnink et al. [9] | SARS-CoV-2_27_RIGHT | TCTACACCACAGAAAACTCCTGGT |
| Munnink et al. [9] | SARS-CoV-2_28_LEFT | CCTTGAAGGTTCTGTTAGAGTGGT |

(continued on next page)

(Table S4 continued)

| Name of Set | Primer/probe | Sequence 5′ → 3′ |
|---|---|---|
| Munnink et al. [9] | SARS-CoV-2_28_RIGHT | AGGTGTGAACATAACCATCCACTG |
| Munnink et al. [9] | SARS-CoV-2_29_LEFT | ACAGTCATGTAGTTGCCTTTAATACTTTAC |
| Munnink et al. [9] | SARS-CoV-2_29_RIGHT | GAGCCTTTGCGAGATGACAACA |
| Munnink et al. [9] | SARS-CoV-2_30_LEFT | AGAAATGTATCTAAAGTTGCGTAGTGATG |
| Munnink et al. [9] | SARS-CoV-2_30_RIGHT | CCCTGAGTTGAACATTACCAGCC |
| Munnink et al. [9] | SARS-CoV-2_31_LEFT | AACGGTCTTTGGCTTGATGACG |
| Munnink et al. [9] | SARS-CoV-2_31_RIGHT | ACCTTCTAAGTCTGTGCCAGCA |
| Munnink et al. [9] | SARS-CoV-2_32_LEFT | TACCAATGTGCTATGAGGCCCA |
| Munnink et al. [9] | SARS-CoV-2_32_RIGHT | GCACTACCCAATATGGTACGTCC |
| Munnink et al. [9] | SARS-CoV-2_33_LEFT | TGTGGCTATGAAGTACAATTATGAACCT |
| Munnink et al. [9] | SARS-CoV-2_33_RIGHT | AGCTACAGTGGCAAGAGAAGGT |
| Munnink et al. [9] | SARS-CoV-2_34_LEFT | GTCCAGAGTACTCAATGGTCTTTGT |
| Munnink et al. [9] | SARS-CoV-2_34_RIGHT | ACCTCTGGCCAAAAACATGACA |
| Munnink et al. [9] | SARS-CoV-2_35_LEFT | TGGTGCTAGGAGAGTGTGGACA |
| Munnink et al. [9] | SARS-CoV-2_35_RIGHT | GGCTACTTTGATACAAGGTTTGCC |
| Munnink et al. [9] | SARS-CoV-2_36_LEFT | CGCTACTTTAGACTGACTCTTGGTG |
| Munnink et al. [9] | SARS-CoV-2_36_RIGHT | ATCACCATTAGCAACAGCCTGC |
| Munnink et al. [9] | SARS-CoV-2_37_LEFT | CTTTCCATGCAGGGTGCTGTAG |
| Munnink et al. [9] | SARS-CoV-2_37_RIGHT | GTTTGGCTGCTGTTGTAAGAGGT |
| Munnink et al. [9] | SARS-CoV-2_38_LEFT | AGATCTGAGGACAAGAGGGCAA |
| Munnink et al. [9] | SARS-CoV-2_38_RIGHT | TGTCATCAGTGCAAGCAGTTTGT |

(continued on next page)

(Table S4 continued)

| Name of Set | Primer/probe | Sequence 5′ → 3′ |
|---|---|---|
| Munnink et al. [9] | SARS-CoV-2_39_LEFT | ACAATTCACCTAATTTAGCATGGCCT |
| Munnink et al. [9] | SARS-CoV-2_39_RIGHT | TTGGTTGTCCCCCACTAGCTAG |
| Munnink et al. [9] | SARS-CoV-2_40_LEFT | GGTATGGTACTTGGTAGTTTAGCTGC |
| Munnink et al. [9] | SARS-CoV-2_40_RIGHT | ACGATTGTGCATCAGCTGACTG |
| Munnink et al. [9] | SARS-CoV-2_41_LEFT | AGTATGTACAAATACCTACAACTTGTGCT |
| Munnink et al. [9] | SARS-CoV-2_41_RIGHT | AGCATAGACGAGGTCTGCCATT |
| Munnink et al. [9] | SARS-CoV-2_42_LEFT | TCTCTAACTACCAACATGAAGAAACAATTT |
| Munnink et al. [9] | SARS-CoV-2_42_RIGHT | GCAGTTAAAGCCCTGGTCAAGG |
| Munnink et al. [9] | SARS-CoV-2_43_LEFT | TGGTATTGTTGGTGTACTGACATTAGA |
| Munnink et al. [9] | SARS-CoV-2_43_RIGHT | AGGGTCAGCAGCATACACAAGT |
| Munnink et al. [9] | SARS-CoV-2_44_LEFT | TGGACCACTAGTGAGAAAAATATTTGTTG |
| Munnink et al. [9] | SARS-CoV-2_44_RIGHT | ACAGCCACCATCGTAACAATCA |
| Munnink et al. [9] | SARS-CoV-2_45_LEFT | CACTTCTTCTTTGCTCAGGATGGT |
| Munnink et al. [9] | SARS-CoV-2_45_RIGHT | AACATGTTGTGCCAACCACCAT |
| Munnink et al. [9] | SARS-CoV-2_46_LEFT | TGCAAAGAATAGAGCTCGCACC |
| Munnink et al. [9] | SARS-CoV-2_46_RIGHT | TGCATTAACATTGGCCGTGACA |
| Munnink et al. [9] | SARS-CoV-2_47_LEFT | TGTAGCTTGTCACACCGTTTCT |
| Munnink et al. [9] | SARS-CoV-2_47_RIGHT | AGTAAGGTCAGTCTCAGTCCAACA |
| Munnink et al. [9] | SARS-CoV-2_48_LEFT | CTCTCTGACGATGCTGTTGTGT |
| Munnink et al. [9] | SARS-CoV-2_48_RIGHT | TGCGGTGTGTACATAGCCTCAT |
| Munnink et al. [9] | SARS-CoV-2_49_LEFT | AGGAGTATGCTGATGTCTTTCATTTGT |

(continued on next page)

(Table S4 continued)

| Name of Set | Primer/probe | Sequence 5′ → 3′ |
|---|---|---|
| Munnink et al. [9] | SARS-CoV-2_49_RIGHT | CACCAGCATTTGTCCAGTCACA |
| Munnink et al. [9] | SARS-CoV-2_50_LEFT | AGGAGGTATGAGCTATTATTGTAAATCACA |
| Munnink et al. [9] | SARS-CoV-2_50_RIGHT | GTTGTACCTCGGTAAACAACAGCA |
| Munnink et al. [9] | SARS-CoV-2_51_LEFT | TCTTTCATGGGAAGTTGGTAAACCT |
| Munnink et al. [9] | SARS-CoV-2_51_RIGHT | TCACATAGTGCATCAACAGCGG |
| Munnink et al. [9] | SARS-CoV-2_52_LEFT | TGCAAATTATCAAAAGGTTGGTATGCA |
| Munnink et al. [9] | SARS-CoV-2_52_RIGHT | CCGAGGAACATGTCTGGACCTA |
| Munnink et al. [9] | SARS-CoV-2_53_RIGHT | CAAGAGTGAGCTGTTTCAGTGGT |
| Munnink et al. [9] | SARS-CoV-2_54_LEFT | TGGAGAAAAGCTGTCTTTATTTCACCT |
| Munnink et al. [9] | SARS-CoV-2_54_RIGHT | GCTTCTTCGCGGGTGATAAACA |
| Munnink et al. [9] | SARS-CoV-2_55_LEFT | TGTTGACACTAAATTCAAAACTGAAGGT |
| Munnink et al. [9] | SARS-CoV-2_55_RIGHT | TGTCAACTCAAAGCCATGTGCC |
| Munnink et al. [9] | SARS-CoV-2_56_LEFT | ACCACCGCCTGGAGATCAATTT |
| Munnink et al. [9] | SARS-CoV-2_56_RIGHT | CGCTTAACAAAGCACTCGTGGA |
| Munnink et al. [9] | SARS-CoV-2_57_LEFT | CGTTTATGATTGATGTTCAACAATGGGG |
| Munnink et al. [9] | SARS-CoV-2_57_RIGHT | ACAACCAGGCAAGTTAAGGTTAGA |
| Munnink et al. [9] | SARS-CoV-2_58_LEFT | GCCTTGTAGTGACAAAGCTTATAAAATAGA |
| Munnink et al. [9] | SARS-CoV-2_58_RIGHT | AAACCCACAAGCTAAAGCCAGC |
| Munnink et al. [9] | SARS-CoV-2_59_LEFT | AGTCTCATGGAAAACAAGTAGTGTCA |
| Munnink et al. [9] | SARS-CoV-2_59_RIGHT | ATTAGCAGCAATGTCCACACCC |
| Munnink et al. [9] | SARS-CoV-2_60_LEFT | CAGGGTGAAGTACCAGTTTCTATCATT |

(continued on next page)

(Table S4 continued)

| Name of Set | Primer/probe | Sequence 5′ → 3′ |
|---|---|---|
| Munnink et al. [9] | SARS-CoV-2_60_RIGHT | GAGTAAAGTAAGTTTCAGGTAATTGTTGG |
| Munnink et al. [9] | SARS-CoV-2_61_LEFT | AGAAATGCCCGTAATGGTGTTCT |
| Munnink et al. [9] | SARS-CoV-2_61_RIGHT | TGAACCTGTTTGCGCATCTGTT |
| Munnink et al. [9] | SARS-CoV-2_62_LEFT | TCGTTTATGGAGATTTTAGTCATAGTCAGT |
| Munnink et al. [9] | SARS-CoV-2_62_RIGHT | TTGCGACATTCATCATTATGCCTTT |
| Munnink et al. [9] | SARS-CoV-2_63_LEFT | TGGCCATGTAGAAACATTTTACCCA |
| Munnink et al. [9] | SARS-CoV-2_63_RIGHT | ATAGCCACGGAACCTCCAAGAG |
| Munnink et al. [9] | SARS-CoV-2_64_LEFT | CTGTACATACAGCTAATAAATGGGATCTCA |
| Munnink et al. [9] | SARS-CoV-2_64_RIGHT | TTTGACCTTCTTTTAAAGACATAACAGCA |
| Munnink et al. [9] | SARS-CoV-2_65_RIGHT | ACCTCTTAGTACCATTGGTCCCA |
| Munnink et al. [9] | SARS-CoV-2_66_RIGHT | ACCCTGTTTTCCTTCAAGGTCC |
| Munnink et al. [9] | SARS-CoV-2_67_LEFT | CAATTTTGTAATGATCCATTTTTGGGTGT |
| Munnink et al. [9] | SARS-CoV-2_67_RIGHT | GGTCAAGTGCACAGTCTACAGC |
| Munnink et al. [9] | SARS-CoV-2_68_LEFT | ACATAGAAGTTATTTGACTCCTGGTGA |
| Munnink et al. [9] | SARS-CoV-2_68_RIGHT | CCCTGGAGCGATTTGTCTGACT |
| Munnink et al. [9] | SARS-CoV-2_69_LEFT | ACTGTGTTGCTGATTATTCTGTCCT |
| Munnink et al. [9] | SARS-CoV-2_69_RIGHT | TAGGTCCACAAACAGTTGCTGG |
| Munnink et al. [9] | SARS-CoV-2_70_LEFT | CCGGTAGCACACCTTGTAATGG |
| Munnink et al. [9] | SARS-CoV-2_70_RIGHT | CCCCTATTAAACAGCCTGCACG |
| Munnink et al. [9] | SARS-CoV-2_71_LEFT | ACTTCTAACCAGGTTGCTGTTCTT |
| Munnink et al. [9] | SARS-CoV-2_71_RIGHT | CAGCTATTCCAGTTAAAGCACGGT |

(continued on next page)

(Table S4 continued)

| Name of Set | Primer/probe | Sequence 5′ → 3′ |
|---|---|---|
| Munnink et al. [9] | SARS-CoV-2_72_LEFT | TGTTACCACAGAAATTCTACCAGTGT |
| Munnink et al. [9] | SARS-CoV-2_72_RIGHT | TACCCGCTAACAGTGCAGAAGT |
| Munnink et al. [9] | SARS-CoV-2_73_LEFT | CTTGCAGATGCTGGCTTCATCA |
| Munnink et al. [9] | SARS-CoV-2_73_RIGHT | TGCACTTCAGCCTCAACTTTGT |
| Munnink et al. [9] | SARS-CoV-2_74_LEFT | GTGCACTTGGAAAACTTCAAGATGT |
| Munnink et al. [9] | SARS-CoV-2_74_RIGHT | TGTTACAAACCAGTGTGTGCCA |
| Munnink et al. [9] | SARS-CoV-2_75_LEFT | CTTCCCTCAGTCAGCACCTCAT |
| Munnink et al. [9] | SARS-CoV-2_75_RIGHT | CAAGCCAGCTATAAAACCTAGCCA |
| Munnink et al. [9] | SARS-CoV-2_76_LEFT | GTTGATTTAGGTGACATCTCTGGCA |
| Munnink et al. [9] | SARS-CoV-2_76_RIGHT | AGCGCTCTGAAAAACAGCAAGA |
| Munnink et al. [9] | SARS-CoV-2_77_LEFT | TGGAACTGTAACTTTGAAGCAAGGT |
| Munnink et al. [9] | SARS-CoV-2_77_RIGHT | GACTTGTTGTGCCATCACCTGA |
| Munnink et al. [9] | SARS-CoV-2_78_LEFT | CTTTGGCTTTGCTGGAAATGCC |
| Munnink et al. [9] | SARS-CoV-2_78_RIGHT | GTGCTTACAAAGGCACGCTAGT |
| Munnink et al. [9] | SARS-CoV-2_79_LEFT | GGTGTTGAACATGTTACCTTCTTCAT |
| Munnink et al. [9] | SARS-CoV-2_79_RIGHT | GTACCGTTGGAATCTGCCATGG |
| Munnink et al. [9] | SARS-CoV-2_80_LEFT | ACGTGAGTCTTGTAAAACCTTCTTTTT |
| Munnink et al. [9] | SARS-CoV-2_80_RIGHT | AATGACCACATGGAACGCGTAC |
| Munnink et al. [9] | SARS-CoV-2_81_LEFT | CTTGTTTTGTGCTTGCTGCTGT |
| Munnink et al. [9] | SARS-CoV-2_81_RIGHT | ACTGCTACTGGAATGGTCTGTGT |
| Munnink et al. [9] | SARS-CoV-2_82_LEFT | GGACCTGCCTAAAGAAATCACTGT |

(continued on next page)

(Table S4 continued)

| Name of Set | Primer/probe | Sequence 5′ → 3′ |
|---|---|---|
| Munnink et al. [9] | SARS-CoV-2_82_RIGHT | TGCCCTCGTATGTTCCAGAAGA |
| Munnink et al. [9] | SARS-CoV-2_83_LEFT | TGAAGAGCAACCAATGGAGATTGA |
| Munnink et al. [9] | SARS-CoV-2_83_RIGHT | TGTTCGTTTAGGCGTGACAAGT |
| Munnink et al. [9] | SARS-CoV-2_84_LEFT | CTTCACACTCAAAAGAAAGACAGAATGA |
| Munnink et al. [9] | SARS-CoV-2_84_RIGHT | ACGAACAACGCACTACAAGACT |
| Munnink et al. [9] | SARS-CoV-2_85_LEFT | AGCACCTTTAATTGAATTGTGCGTG |
| Munnink et al. [9] | SARS-CoV-2_85_RIGHT | CGTCTGGTAGCTCTTCGGTAGT |
| Munnink et al. [9] | SARS-CoV-2_86_LEFT | GGCCCCAAGGTTTACCCAATAA |
| Munnink et al. [9] | SARS-CoV-2_86_RIGHT | CTGTTGCGACTACGTGATGAGG |
| Munnink et al. [9] | SARS-CoV-2_87_LEFT | AAAAGATCACATTGGCACCCGC |
| Munnink et al. [9] | SARS-CoV-2_87_RIGHT | CGACATTCCGAAGAACGCTGAA |
| Munnink et al. [9] | SARS-CoV-2_88_LEFT | TAACACAAGCTTTCGGCAGACG |
| Munnink et al. [9] | SARS-CoV-2_88_RIGHT | GTGGTCTGCATGAGTTTAGGCC |
| Munnink et al. [9] | SARS-CoV-2_89_LEFT | AGGCTGATGAAACTCAAGCCTT |
| Munnink et al. [9] | SARS-CoV-2_89_RIGHT | AAAATCACATGGGGATAGCACTACT |
| Nalla et al. [10] | nCoV_2019-F | CAAATTCTATGGTGGTTGGCACA |
| Nalla et al. [10] | nCoV_2019-P1 | ATAATCCCAACCCATAAG |
| Nalla et al. [10] | nCoV_2019-P2 | ATAATCCCAACCCATGAG |
| Nalla et al. [10] | nCoV_2019-R | GGCATGGCTCTATCACATTTAGG |
| Niu et al. [11] | E-F | TTCTTGCTTTCGTGGTATTC |
| Niu et al. [11] | E-P | GTTACACTAGCCATCCTTACTGCGCTTCGA |

(continued on next page)

(Table S4 continued)

| Name of Set | Primer/probe | Sequence 5′ → 3′ |
|---|---|---|
| Niu et al. [11] | E-R | CACGTTAACAATATTGCAGC |
| Niu et al. [11] | RdRp-F | GGTCATGTGTGGCGGCTC |
| Niu et al. [11] | RdRp-P | CTATATGTTAAACCAGGTGGAAC |
| Niu et al. [11] | RdRp-R | GCTGTAACAGCTTGACAAATGAAAG |
| Park et al. [12] | SARS-CoV-2_IBS_E2-F | TTCGGAAGAGACAGGTACGTTA |
| Park et al. [12] | SARS-CoV-2_IBS_E2-R | AGCAGTACGCACACAATCG |
| Park et al. [12] | SARS-CoV-2_IBS_N1-F | CAATGCTGCAATCGTGCTAC |
| Park et al. [12] | SARS-CoV-2_IBS_N1-R | GTTGCGACTACGTGATGAGG |
| Park et al. [12] | SARS-CoV-2_IBS_RdRP2-F | AGAATAGAGCTCGCACCGTA |
| Park et al. [12] | SARS-CoV-2_IBS_RdRP2-R | CTCCTCTAGTGGCGGCTATT |
| Park et al. [12] | SARS-CoV-2_IBS_S2-F | GCTGGTGCTGCAGCTTATTA |
| Park et al. [12] | SARS-CoV-2_IBS_S2-R | AGGGTCAAGTGCACAGTCTA |
| Park et al. [12] | SARS-CoV-2_IBS_m_N1-F | AAGGAAATTTTGGGGACCAG |
| Park et al. [12] | SARS-CoV-2_IBS_m_N1-R | GAGTCAGCACTGCTCATGGA |
| Park et al. [12] | SARS-CoV-2_IBS_m_N2-F | AAAGGCCAACAACAACAAGG |
| Park et al. [12] | SARS-CoV-2_IBS_m_N2-R | GCTCTGTTGGTGGGAATGTT |
| Park et al. [12] | SARS-CoV-2_IBS_m_RdRP1-F | GCTCGCAAACATACAACGTG |
| Park et al. [12] | SARS-CoV-2_IBS_m_RdRP1-R | CATTAACATTGGCCGTGACA |
| Park et al. [12] | SARS-CoV-2_IBS_m_RdRP2-F | TGAAATCAATAGCCGCCACT |
| Park et al. [12] | SARS-CoV-2_IBS_m_RdRP2-R | TGTTTGCGAGCAAGAACAAG |
| Park et al. [12] | SARS-CoV-2_IBS_m_S1-F | CAGATGCTGGCTTCATCAAA |

(continued on next page)

(Table S4 continued)

| Name of Set | Primer/probe | Sequence 5′ → 3′ |
|---|---|---|
| Park et al. [12] | SARS-CoV-2_IBS_m_S1-R | GGTTGGCAATCAATTTTTGG |
| Park et al. [12] | SARS-CoV-2_IBS_m_S2-F | ACTGTTTTGCCACCTTTGCT |
| Park et al. [12] | SARS-CoV-2_IBS_m_S2-R | AGCTTGTGCATTTTGGTTGA |
| Rahman et al. [13] | M-F | CAAGGACCTGCCTAAAGAAATCAC |
| Rahman et al. [13] | M-P | TGTTGCTACATCACGAACGCTTTC |
| Rahman et al. [13] | M-R | ACGCTGCGAAGCTCCCAAT |
| Rahman et al. [13] | ORF1b-F | CATGGTGGACAGCCTTTGTTAC |
| Rahman et al. [13] | ORF1b-P | AATGTGAATGCGTCATCATCTGAAGCA |
| Rahman et al. [13] | ORF1b-R | TCGCGTGGTTTGCCAAGAT |
| Toptan et al. [14] | M-475-F | TGTGACATCAAGGACCTGCC |
| Toptan et al. [14] | M-507-P | TGTTGCTACATCACGAACGC |
| Toptan et al. [14] | M-574-R | CTGAGTCACCTGCTACACGC |
| Toptan et al. [14] | S-1850-F | GCACAGAAGTCCCTGTTGCT |
| Toptan et al. [14] | S-1884-P | ACTTACTCCTACTTGGCGTGT |
| Toptan et al. [14] | S-1949-R | AAACAGCCTGCACGTGTTTG |
| Vogels et al. [15] | HKU-N-F | TAATCAGACAAGGAACTGATTA |
| Vogels et al. [15] | HKU-N-P | GCAAATTGTGCAATTTGCGG |
| Vogels et al. [15] | HKU-N-R | CGAAGGTGTGACTTCCATG |
| Vogels et al. [15] | HKU-ORF1-F1 | TGGGGCTTTACAGGTAACCT |
| Vogels et al. [15] | HKU-ORF1-F2 | TGGGGCTTTACGGGTAACCT |
| Vogels et al. [15] | HKU-ORF1-F3 | TGGGGTTTTACAGGTAACCT |

(continued on next page)

(Table S4 continued)

| Name of Set | Primer/probe | Sequence 5′ → 3′ |
|---|---|---|
| Vogels et al. [15] | HKU-ORF1-F4 | TGGGGTTTTACGGGTAACCT |
| Vogels et al. [15] | HKU-ORF1-P1 | TAGTTGTGATGCAATCATGACTAG |
| Vogels et al. [15] | HKU-ORF1-P2 | TAGTTGTGATGCTATCATGACTAG |
| Vogels et al. [15] | HKU-ORF1-R1 | AACACGCTTAACAAAGCACTC |
| Vogels et al. [15] | HKU-ORF1-R2 | AACGCGCTTAACAAAGCACTC |
| WHO | NIID_WH-1_F24381 | TCAAGACTCACTTTCTTCCAC |
| WHO | NIID_WH-1_F501 | TTCGGATGCTCGAACTGCACC |
| WHO | NIID_WH-1_F509 | CTCGAACTGCACCTCATGG |
| WHO | NIID_WH-1_R24873 | ATTTGAAACAAAGACACCTTCAC |
| WHO | NIID_WH-1_R854 | CAGAAGTTGTTATCGACATAGC |
| WHO | NIID_WH-1_R913 | CTTTACCAGCACGTGCTAGAAGG |
| WHO | NIID_WH-1_Seq_F24383 | AAGACTCACTTTCTTCCACAG |
| WHO | NIID_WH-1_Seq_F519 | ACCTCATGGTCATGTTATGG |
| WHO | NIID_WH-1_Seq_R24865 | CAAAGACACCTTCACGAGG |
| WHO | NIID_WH-1_Seq_R840 | GACATAGCGAGTGTATGCC |
| WHO | WH-NICN-F | CGTTTGGTGGACCCTCAGAT |
| WHO | WH-NICN-P | CAACTGGCAGTAACCA |
| WHO | WH-NICN-R | CCCCACTGCGTTCTCCATT |
| WHO | WuhanCoV-spk1-f | TTGGCAAAATTCAAGACTCACTTT |
| WHO | WuhanCoV-spk2-r | TGTGGTTCATAAAAATTCCTTTGTG |
| WHO | nCoV_IP2-12669Fw | ATGAGCTTAGTCCTGTTG |

(continued on next page)

(Table S4 continued)

| Name of Set | Primer/probe | Sequence $5' \rightarrow 3'$ |
|---|---|---|
| WHO | nCoV_IP2-12696bProbe | AGATGTCTTGTGCTGCCGGTA |
| WHO | nCoV_IP2-12759Rv | CTCCCTTTGTTGTGTTGT |
| WHO | nCoV_IP4-14059Fw | GGTAACTGGTATGATTTCG |
| WHO | nCoV_IP4-14084Probe | TCATACAAACCACGCCAGG |
| WHO | nCoV_IP4-14146Rv | CTGGTCAAGGTTAATATAGG |
| Yip et al. [16] | nsp2f | ATGCATTTGCATCAGAGGCT |
| Yip et al. [16] | nsp2r | TTGTTATAGCGGCCTTCTGT |

Table S5: Reference melting temperatures and coverages for SARS-CoV-2.

| Set | Primer/probe | $T_{\text{ref.}}$ (°C) | $C_{\text{strict}}$ (%) | $C_{\text{part.}}$ (%) | $N_{imp}$ |
|---|---|---|---|---|---|
| Bhadra et al. [1] | E-PF | 62.6 | 99.3 | 99.4 | 1 |
| Bhadra et al. [1] | E-Probe | 64.2 | 99.3 | 99.3 | 1 |
| Bhadra et al. [1] | E-RP | 61.4 | 99.3 | 99.3 | 0 |
| Bhadra et al. [1] | M-PF | 62.2 | 99.3 | 99.3 | 0 |
| Bhadra et al. [1] | M-Probe | 61.3 | 99.4 | 99.4 | 0 |
| Bhadra et al. [1] | M-RP | 63.3 | 98.7 | 98.7 | 0 |
| CDC | 2019-nCoV_N1-F | 62.4 | 99.3 | 99.3 | 0 |
| CDC | 2019-nCoV_N1-P | 75.5 | 98.4 | 99.4 | 223 |
| CDC | 2019-nCoV_N1-R | 69.0 | 99.2 | 99.4 | 37 |
| CDC | 2019-nCoV_N2-F | 61.1 | 99.0 | 99.0 | 0 |
| CDC | 2019-nCoV_N2-P | 74.7 | 98.9 | 99.1 | 32 |
| CDC | 2019-nCoV_N2-R | 63.3 | 99.3 | 99.3 | 1 |
| Corman et al. [2] | E_Sarbeco_F | 65.6 | 99.4 | 99.4 | 1 |
| Corman et al. [2] | E_Sarbeco_P1 | 78.2 | 99.3 | 99.3 | 12 |
| Corman et al. [2] | E_Sarbeco_R | 65.9 | 99.3 | 99.3 | 5 |
| Corman et al. [2] | N_Sarbeco_F | 64.6 | 99.2 | 99.2 | 1 |
| Corman et al. [2] | N_Sarbeco_P | 71.9 | 98.8 | 99.1 | 60 |
| Corman et al. [2] | N_Sarbeco_R | 65.1 | 97.6 | 97.6 | 1 |
| Corman et al. [2] | RdRp_SARSr-FA | 69.1 | 99.3 | 99.4 | 32 |

(continued on next page)

(Table S5, SARS-Cov-2 coverage, continued)

| Set | Primer/probe | $T_{\text{ref.}}$ (°C) | $C_{\text{strict}}$ (%) | $C_{\text{part.}}$ (%) | $N_{imp}$ |
|---|---|---|---|---|---|
| Corman et al. [2] | RdRp_SARSr-FG | 70.7 | 0.000 | 99.3 | 21522 |
| Corman et al. [2] | RdRp_SARSr-P1-1 | 79.2 | 0.000 | 99.4 | 21540 |
| Corman et al. [2] | RdRp_SARSr-P1-2 | 81.7 | 0.000 | 99.3 | 21521 |
| Corman et al. [2] | RdRp_SARSr-P1-3 | 77.3 | 0.000 | 99.4 | 21540 |
| Corman et al. [2] | RdRp_SARSr-P1-4 | 79.9 | 0.000 | 99.3 | 21521 |
| Corman et al. [2] | RdRp_SARSr-P1-5 | 78.2 | 0.000 | 99.4 | 21538 |
| Corman et al. [2] | RdRp_SARSr-P1-6 | 80.7 | 0.000 | 0.000 | 0 |
| Corman et al. [2] | RdRp_SARSr-P1-7 | 76.2 | 0.000 | 99.4 | 21539 |
| Corman et al. [2] | RdRp_SARSr-P1-8 | 78.8 | 0.000 | 0.0692 | 15 |
| Corman et al. [2] | RdRp_SARSr-P2 | 76.3 | 99.3 | 99.4 | 26 |
| Corman et al. [2] | RdRp_SARSr-R1 | 61.7 | 0.000 | 0.000 | 0 |
| Corman et al. [2] | RdRp_SARSr-R2 | 62.1 | 0.000 | 0.000 | 0 |
| Corman et al. [2] | RdRp_SARSr-R3 | 63.5 | 0.000 | 0.000 | 0 |
| Corman et al. [2] | RdRp_SARSr-R4 | 63.8 | 0.000 | 0.000 | 0 |
| Davda et al. [3] | MF1 | 56.5 | 98.7 | 98.7 | 0 |
| Davda et al. [3] | MF2 | 58.6 | 99.4 | 99.4 | 0 |
| Davda et al. [3] | MR1 | 58.9 | 99.2 | 99.2 | 0 |
| Davda et al. [3] | MR2 | 60.2 | 99.3 | 99.3 | 0 |
| Davda et al. [3] | N1F1 | 66.7 | 99.3 | 99.4 | 10 |
| Davda et al. [3] | N1F2 | 70.3 | 99.0 | 99.1 | 24 |
| Davda et al. [3] | N1R1 | 62.2 | 99.3 | 99.3 | 0 |

(continued on next page)

(Table S5, SARS-Cov-2 coverage, continued)

| Set | Primer/probe | $T_{\text{ref.}}$ (°C) | $C_{\text{strict}}$ (%) | $C_{\text{part.}}$ (%) | $N_{imp}$ |
|---|---|---|---|---|---|
| Davda et al. [3] | N1R2 | 68.0 | 99.1 | 99.1 | 5 |
| Davda et al. [3] | N2F1 | 60.3 | 99.3 | 99.3 | 0 |
| Davda et al. [3] | N2F2 | 57.6 | 99.2 | 99.2 | 0 |
| Davda et al. [3] | N2R1 | 59.9 | 95.3 | 95.3 | 0 |
| Davda et al. [3] | N2R2 | 57.0 | 99.4 | 99.4 | 0 |
| Davda et al. [3] | Orf1abF1 | 69.7 | 99.1 | 99.4 | 54 |
| Davda et al. [3] | Orf1abF2 | 63.2 | 99.0 | 99.0 | 0 |
| Davda et al. [3] | Orf1abR1 | 67.4 | 99.0 | 99.0 | 12 |
| Davda et al. [3] | Orf1abR2 | 61.7 | 98.9 | 98.9 | 0 |
| Grant et al. [4] | NgeneProbe | 79.9 | 99.3 | 99.5 | 32 |
| Grant et al. [4] | NgeneTaq1 | 62.4 | 97.5 | 97.5 | 0 |
| Grant et al. [4] | NgeneTaq2 | 63.8 | 99.2 | 99.2 | 0 |
| Hirotsu et al. [5] | NIID-N2-F | 60.6 | 99.2 | 99.2 | 0 |
| Hirotsu et al. [5] | NIID-N2-P | 68.8 | 99.4 | 99.4 | 7 |
| Hirotsu et al. [5] | NIID-N2-R | 63.9 | 0.000 | 0.000 | 0 |
| Lanza et al. [6] | UFRN_1_F | 63.7 | 99.0 | 99.1 | 11 |
| Lanza et al. [6] | UFRN_1_P | 75.0 | 99.3 | 99.4 | 36 |
| Lanza et al. [6] | UFRN_1_R | 62.9 | 99.3 | 99.3 | 0 |
| Lanza et al. [6] | UFRN_2_F | 62.2 | 99.4 | 99.4 | 0 |
| Lanza et al. [6] | UFRN_2_P | 69.7 | 98.5 | 98.8 | 65 |
| Lanza et al. [6] | UFRN_2_R | 62.0 | 99.1 | 99.1 | 0 |

(continued on next page)

(Table S5, SARS-Cov-2 coverage, continued)

| Set | Primer/probe | $T_{\text{ref.}}$ (°C) | $C_{\text{strict}}$ (%) | $C_{\text{part.}}$ (%) | $N_{imp}$ |
|---|---|---|---|---|---|
| Lanza et al. [6] | UFRN_3_F | 62.9 | 99.0 | 99.0 | 2 |
| Lanza et al. [6] | UFRN_3_P | 69.1 | 99.0 | 99.1 | 21 |
| Lanza et al. [6] | UFRN_3_R | 60.5 | 99.0 | 99.0 | 0 |
| Lanza et al. [6] | UFRN_4_F | 62.0 | 99.3 | 99.3 | 0 |
| Lanza et al. [6] | UFRN_4_P | 69.0 | 99.0 | 99.5 | 107 |
| Lanza et al. [6] | UFRN_4_R | 64.9 | 99.3 | 99.3 | 2 |
| Lanza et al. [6] | UFRN_5_F | 62.2 | 99.2 | 99.2 | 0 |
| Lanza et al. [6] | UFRN_5_P | 73.0 | 98.8 | 98.9 | 25 |
| Lanza et al. [6] | UFRN_5_R | 63.7 | 98.9 | 98.9 | 0 |
| Lanza et al. [6] | UFRN_6_F | 63.2 | 98.9 | 98.9 | 0 |
| Lanza et al. [6] | UFRN_6_P | 71.3 | 99.0 | 99.2 | 42 |
| Lanza et al. [6] | UFRN_6_R | 62.4 | 99.2 | 99.2 | 0 |
| Lanza et al. [6] | UFRN_7_F | 62.5 | 98.9 | 98.9 | 0 |
| Lanza et al. [6] | UFRN_7_P | 71.7 | 99.1 | 99.2 | 36 |
| Lanza et al. [6] | UFRN_7_R | 61.5 | 99.2 | 99.2 | 0 |
| Lanza et al. [6] | UFRN_8_F | 61.3 | 99.4 | 99.4 | 0 |
| Lanza et al. [6] | UFRN_8_P | 70.5 | 99.4 | 99.4 | 11 |
| Lanza et al. [6] | UFRN_8_R | 62.0 | 99.4 | 99.4 | 0 |
| Lanza et al. [6] | UFRN_9_F | 60.5 | 99.4 | 99.4 | 0 |
| Lanza et al. [6] | UFRN_9_P | 72.6 | 99.4 | 99.5 | 24 |
| Lanza et al. [6] | UFRN_9_R | 62.4 | 99.2 | 99.2 | 0 |

(continued on next page)

(Table S5, SARS-Cov-2 coverage, continued)

| Set | Primer/probe | $T_{\text{ref.}}$ (°C) | $C_{\text{strict}}$ (%) | $C_{\text{part.}}$ (%) | $N_{imp}$ |
|---|---|---|---|---|---|
| Li et al. [7] | S-1 | 70.4 | 98.2 | 98.3 | 5 |
| Li et al. [7] | S-2 | 67.5 | 99.2 | 99.2 | 5 |
| Lu et al. [8] | ORF1a-F | 64.0 | 99.4 | 99.4 | 0 |
| Lu et al. [8] | ORF1a-P | 74.7 | 98.6 | 99.3 | 150 |
| Lu et al. [8] | ORF1a-R | 68.8 | 99.3 | 99.4 | 20 |
| Luminex | China_N_F | 67.1 | 64.9 | 65.1 | 54 |
| Luminex | China_N_P | 63.1 | 99.2 | 99.2 | 0 |
| Luminex | China_N_R | 65.6 | 96.4 | 96.4 | 4 |
| Luminex | China_ORF_F | 59.4 | 99.5 | 99.5 | 0 |
| Luminex | China_ORF_P | 81.0 | 98.9 | 99.0 | 9 |
| Luminex | China_ORF_R | 61.5 | 99.1 | 99.1 | 0 |
| Munnink et al. [9] | SARS-CoV-2_1_LEFT | 67.9 | 45.9 | 46.0 | 8 |
| Munnink et al. [9] | SARS-CoV-2_1_RIGHT | 69.9 | 97.4 | 98.9 | 313 |
| Munnink et al. [9] | SARS-CoV-2_2_LEFT | 69.4 | 98.9 | 99.0 | 25 |
| Munnink et al. [9] | SARS-CoV-2_2_RIGHT | 67.7 | 99.4 | 99.4 | 6 |
| Munnink et al. [9] | SARS-CoV-2_3_LEFT | 68.7 | 99.3 | 99.3 | 9 |
| Munnink et al. [9] | SARS-CoV-2_3_RIGHT | 68.4 | 98.9 | 99.3 | 89 |
| Munnink et al. [9] | SARS-CoV-2_4_LEFT | 66.8 | 70.0 | 70.0 | 1 |
| Munnink et al. [9] | SARS-CoV-2_4_RIGHT | 66.6 | 99.1 | 99.2 | 21 |
| Munnink et al. [9] | SARS-CoV-2_5_LEFT | 67.7 | 98.5 | 98.5 | 5 |
| Munnink et al. [9] | SARS-CoV-2_5_RIGHT | 68.2 | 99.0 | 99.0 | 9 |

(continued on next page)

(Table S5, SARS-Cov-2 coverage, continued)

| Set | Primer/probe | $T_{\mathrm{ref.}}$ (°C) | $C_{\mathrm{strict}}$ (%) | $C_{\mathrm{part.}}$ (%) | $N_{imp}$ |
|---|---|---|---|---|---|
| Munnink et al. [9] | SARS-CoV-2_6_LEFT | 73.2 | 98.3 | 99.3 | 233 |
| Munnink et al. [9] | SARS-CoV-2_6_RIGHT | 66.3 | 99.3 | 99.3 | 1 |
| Munnink et al. [9] | SARS-CoV-2_7_LEFT | 71.3 | 99.1 | 99.2 | 27 |
| Munnink et al. [9] | SARS-CoV-2_7_RIGHT | 67.8 | 99.1 | 99.2 | 10 |
| Munnink et al. [9] | SARS-CoV-2_8_LEFT | 67.2 | 98.9 | 99.0 | 5 |
| Munnink et al. [9] | SARS-CoV-2_8_RIGHT | 65.9 | 98.1 | 98.3 | 36 |
| Munnink et al. [9] | SARS-CoV-2_9_LEFT | 72.3 | 99.0 | 99.1 | 25 |
| Munnink et al. [9] | SARS-CoV-2_9_RIGHT | 69.0 | 98.6 | 98.7 | 22 |
| Munnink et al. [9] | SARS-CoV-2_11_LEFT | 66.9 | 99.2 | 99.3 | 14 |
| Munnink et al. [9] | SARS-CoV-2_11_RIGHT | 71.3 | 98.0 | 99.3 | 280 |
| Munnink et al. [9] | SARS-CoV-2_12_RIGHT | 67.2 | 99.3 | 99.3 | 4 |
| Munnink et al. [9] | SARS-CoV-2_13_LEFT | 70.0 | 99.3 | 99.3 | 19 |
| Munnink et al. [9] | SARS-CoV-2_13_RIGHT | 68.2 | 99.1 | 99.4 | 57 |
| Munnink et al. [9] | SARS-CoV-2_14_LEFT | 72.3 | 99.3 | 99.4 | 6 |
| Munnink et al. [9] | SARS-CoV-2_14_RIGHT | 68.1 | 99.1 | 99.4 | 62 |
| Munnink et al. [9] | SARS-CoV-2_15_LEFT | 70.9 | 99.2 | 99.3 | 37 |
| Munnink et al. [9] | SARS-CoV-2_15_RIGHT | 67.9 | 99.0 | 99.1 | 26 |
| Munnink et al. [9] | SARS-CoV-2_16_LEFT | 69.9 | 98.3 | 98.4 | 35 |
| Munnink et al. [9] | SARS-CoV-2_16_RIGHT | 65.4 | 99.3 | 99.3 | 1 |
| Munnink et al. [9] | SARS-CoV-2_17_LEFT | 66.7 | 99.3 | 99.4 | 22 |
| Munnink et al. [9] | SARS-CoV-2_17_RIGHT | 67.9 | 98.3 | 98.4 | 26 |

(continued on next page)

(Table S5, SARS-Cov-2 coverage, continued)

| Set | Primer/probe | $T_{\text{ref.}}$ (°C) | $C_{\text{strict}}$ (%) | $C_{\text{part.}}$ (%) | $N_{imp}$ |
|---|---|---|---|---|---|
| Munnink et al. [9] | SARS-CoV-2_18_LEFT | 68.0 | 99.2 | 99.3 | 27 |
| Munnink et al. [9] | SARS-CoV-2_18_RIGHT | 71.0 | 98.9 | 99.0 | 21 |
| Munnink et al. [9] | SARS-CoV-2_19_LEFT | 69.6 | 99.4 | 99.5 | 31 |
| Munnink et al. [9] | SARS-CoV-2_19_RIGHT | 69.6 | 99.0 | 99.2 | 45 |
| Munnink et al. [9] | SARS-CoV-2_20_LEFT | 67.0 | 98.9 | 98.9 | 2 |
| Munnink et al. [9] | SARS-CoV-2_21_LEFT | 72.1 | 98.5 | 98.7 | 32 |
| Munnink et al. [9] | SARS-CoV-2_21_RIGHT | 66.8 | 99.2 | 99.2 | 6 |
| Munnink et al. [9] | SARS-CoV-2_22_LEFT | 66.6 | 98.9 | 98.9 | 7 |
| Munnink et al. [9] | SARS-CoV-2_22_RIGHT | 67.7 | 99.4 | 99.4 | 1 |
| Munnink et al. [9] | SARS-CoV-2_23_LEFT | 70.0 | 83.3 | 99.4 | 3496 |
| Munnink et al. [9] | SARS-CoV-2_23_RIGHT | 68.1 | 99.2 | 99.3 | 6 |
| Munnink et al. [9] | SARS-CoV-2_24_LEFT | 72.0 | 99.2 | 99.3 | 39 |
| Munnink et al. [9] | SARS-CoV-2_24_RIGHT | 67.5 | 98.4 | 98.4 | 0 |
| Munnink et al. [9] | SARS-CoV-2_25_LEFT | 71.7 | 99.2 | 99.3 | 30 |
| Munnink et al. [9] | SARS-CoV-2_25_RIGHT | 67.5 | 99.0 | 99.0 | 2 |
| Munnink et al. [9] | SARS-CoV-2_26_LEFT | 70.5 | 99.0 | 99.1 | 8 |
| Munnink et al. [9] | SARS-CoV-2_26_RIGHT | 67.5 | 99.3 | 99.3 | 3 |
| Munnink et al. [9] | SARS-CoV-2_27_LEFT | 68.0 | 98.9 | 98.9 | 13 |
| Munnink et al. [9] | SARS-CoV-2_27_RIGHT | 68.2 | 99.2 | 99.3 | 19 |
| Munnink et al. [9] | SARS-CoV-2_28_LEFT | 67.5 | 99.0 | 99.1 | 32 |
| Munnink et al. [9] | SARS-CoV-2_28_RIGHT | 67.4 | 99.3 | 99.3 | 7 |

(continued on next page)

(Table S5, SARS-Cov-2 coverage, continued)

| Set | Primer/probe | $T_{\text{ref.}}$ (°C) | $C_{\text{strict}}$ (%) | $C_{\text{part.}}$ (%) | $N_{imp}$ |
|---|---|---|---|---|---|
| Munnink et al. [9] | SARS-CoV-2_29_LEFT | 71.6 | 99.2 | 99.5 | 49 |
| Munnink et al. [9] | SARS-CoV-2_29_RIGHT | 69.1 | 97.6 | 97.9 | 53 |
| Munnink et al. [9] | SARS-CoV-2_30_LEFT | 72.3 | 99.4 | 99.5 | 10 |
| Munnink et al. [9] | SARS-CoV-2_30_RIGHT | 69.1 | 98.2 | 98.6 | 84 |
| Munnink et al. [9] | SARS-CoV-2_31_LEFT | 68.4 | 99.2 | 99.2 | 3 |
| Munnink et al. [9] | SARS-CoV-2_31_RIGHT | 67.4 | 99.0 | 99.4 | 81 |
| Munnink et al. [9] | SARS-CoV-2_32_LEFT | 68.1 | 99.4 | 99.4 | 2 |
| Munnink et al. [9] | SARS-CoV-2_32_RIGHT | 67.8 | 99.2 | 99.3 | 14 |
| Munnink et al. [9] | SARS-CoV-2_33_LEFT | 71.5 | 99.0 | 99.3 | 58 |
| Munnink et al. [9] | SARS-CoV-2_33_RIGHT | 66.9 | 99.3 | 99.3 | 3 |
| Munnink et al. [9] | SARS-CoV-2_34_LEFT | 68.6 | 99.4 | 99.4 | 2 |
| Munnink et al. [9] | SARS-CoV-2_34_RIGHT | 66.1 | 99.0 | 99.1 | 6 |
| Munnink et al. [9] | SARS-CoV-2_35_LEFT | 68.7 | 99.3 | 99.3 | 10 |
| Munnink et al. [9] | SARS-CoV-2_35_RIGHT | 68.7 | 99.3 | 99.4 | 9 |
| Munnink et al. [9] | SARS-CoV-2_36_LEFT | 70.5 | 99.5 | 99.7 | 45 |
| Munnink et al. [9] | SARS-CoV-2_36_RIGHT | 68.0 | 99.3 | 99.3 | 2 |
| Munnink et al. [9] | SARS-CoV-2_37_LEFT | 68.5 | 98.9 | 99.0 | 9 |
| Munnink et al. [9] | SARS-CoV-2_37_RIGHT | 67.6 | 99.4 | 99.4 | 16 |
| Munnink et al. [9] | SARS-CoV-2_38_LEFT | 68.5 | 99.3 | 99.4 | 9 |
| Munnink et al. [9] | SARS-CoV-2_38_RIGHT | 67.3 | 99.3 | 99.5 | 51 |
| Munnink et al. [9] | SARS-CoV-2_39_LEFT | 70.8 | 99.3 | 99.4 | 27 |

(continued on next page)

(Table S5, SARS-Cov-2 coverage, continued)

| Set | Primer/probe | $T_{\text{ref.}}$ (°C) | $C_{\text{strict}}$ (%) | $C_{\text{part.}}$ (%) | $N_{imp}$ |
|---|---|---|---|---|---|
| Munnink et al. [9] | SARS-CoV-2_39_RIGHT | 67.6 | 99.2 | 99.3 | 11 |
| Munnink et al. [9] | SARS-CoV-2_40_LEFT | 70.7 | 99.1 | 99.4 | 52 |
| Munnink et al. [9] | SARS-CoV-2_40_RIGHT | 67.9 | 99.0 | 99.1 | 10 |
| Munnink et al. [9] | SARS-CoV-2_41_LEFT | 70.2 | 99.2 | 99.3 | 14 |
| Munnink et al. [9] | SARS-CoV-2_41_RIGHT | 68.4 | 99.3 | 99.4 | 2 |
| Munnink et al. [9] | SARS-CoV-2_42_LEFT | 72.3 | 99.2 | 99.3 | 20 |
| Munnink et al. [9] | SARS-CoV-2_42_RIGHT | 68.8 | 98.9 | 99.4 | 102 |
| Munnink et al. [9] | SARS-CoV-2_43_LEFT | 69.1 | 99.4 | 99.5 | 9 |
| Munnink et al. [9] | SARS-CoV-2_43_RIGHT | 66.7 | 99.3 | 99.3 | 4 |
| Munnink et al. [9] | SARS-CoV-2_44_LEFT | 72.3 | 99.3 | 99.4 | 17 |
| Munnink et al. [9] | SARS-CoV-2_44_RIGHT | 65.7 | 99.5 | 99.5 | 3 |
| Munnink et al. [9] | SARS-CoV-2_45_LEFT | 69.7 | 98.9 | 98.9 | 7 |
| Munnink et al. [9] | SARS-CoV-2_45_RIGHT | 65.0 | 99.3 | 99.3 | 0 |
| Munnink et al. [9] | SARS-CoV-2_46_LEFT | 69.3 | 99.3 | 99.4 | 17 |
| Munnink et al. [9] | SARS-CoV-2_46_RIGHT | 67.3 | 99.3 | 99.3 | 0 |
| Munnink et al. [9] | SARS-CoV-2_47_LEFT | 64.8 | 99.4 | 99.4 | 0 |
| Munnink et al. [9] | SARS-CoV-2_47_RIGHT | 67.9 | 99.3 | 99.5 | 24 |
| Munnink et al. [9] | SARS-CoV-2_48_LEFT | 66.7 | 98.8 | 98.9 | 29 |
| Munnink et al. [9] | SARS-CoV-2_48_RIGHT | 67.3 | 98.2 | 98.3 | 1 |
| Munnink et al. [9] | SARS-CoV-2_49_LEFT | 71.4 | 99.2 | 99.4 | 50 |
| Munnink et al. [9] | SARS-CoV-2_49_RIGHT | 66.8 | 99.2 | 99.3 | 16 |

(continued on next page)

(Table S5, SARS-Cov-2 coverage, continued)

| Set | Primer/probe | $T_{\text{ref.}}$ (°C) | $C_{\text{strict}}$ (%) | $C_{\text{part.}}$ (%) | $N_{imp}$ |
|---|---|---|---|---|---|
| Munnink et al. [9] | SARS-CoV-2_50_LEFT | 73.8 | 99.3 | 99.4 | 36 |
| Munnink et al. [9] | SARS-CoV-2_50_RIGHT | 68.1 | 99.2 | 99.3 | 16 |
| Munnink et al. [9] | SARS-CoV-2_51_LEFT | 68.9 | 99.2 | 99.3 | 17 |
| Munnink et al. [9] | SARS-CoV-2_51_RIGHT | 67.8 | 99.5 | 99.5 | 1 |
| Munnink et al. [9] | SARS-CoV-2_52_LEFT | 71.3 | 99.2 | 99.3 | 9 |
| Munnink et al. [9] | SARS-CoV-2_52_RIGHT | 68.5 | 99.0 | 99.0 | 6 |
| Munnink et al. [9] | SARS-CoV-2_53_RIGHT | 67.3 | 99.3 | 99.4 | 19 |
| Munnink et al. [9] | SARS-CoV-2_54_LEFT | 72.2 | 98.9 | 99.1 | 37 |
| Munnink et al. [9] | SARS-CoV-2_54_RIGHT | 69.3 | 99.2 | 99.3 | 38 |
| Munnink et al. [9] | SARS-CoV-2_55_LEFT | 70.0 | 99.1 | 99.3 | 40 |
| Munnink et al. [9] | SARS-CoV-2_55_RIGHT | 68.1 | 99.4 | 99.5 | 7 |
| Munnink et al. [9] | SARS-CoV-2_56_LEFT | 68.9 | 99.3 | 99.4 | 17 |
| Munnink et al. [9] | SARS-CoV-2_56_RIGHT | 68.4 | 99.1 | 99.4 | 48 |
| Munnink et al. [9] | SARS-CoV-2_57_LEFT | 74.2 | 99.3 | 99.4 | 33 |
| Munnink et al. [9] | SARS-CoV-2_57_RIGHT | 66.7 | 99.0 | 99.1 | 23 |
| Munnink et al. [9] | SARS-CoV-2_58_LEFT | 73.8 | 99.1 | 99.2 | 22 |
| Munnink et al. [9] | SARS-CoV-2_58_RIGHT | 67.9 | 98.7 | 98.7 | 2 |
| Munnink et al. [9] | SARS-CoV-2_59_LEFT | 68.9 | 98.1 | 98.2 | 13 |
| Munnink et al. [9] | SARS-CoV-2_59_RIGHT | 67.0 | 99.1 | 99.1 | 3 |
| Munnink et al. [9] | SARS-CoV-2_60_LEFT | 71.0 | 98.2 | 99.4 | 250 |
| Munnink et al. [9] | SARS-CoV-2_60_RIGHT | 70.4 | 99.1 | 99.3 | 26 |

(continued on next page)

(Table S5, SARS-Cov-2 coverage, continued)

| Set | Primer/probe | $T_{\text{ref.}}$ (°C) | $C_{\text{strict}}$ (%) | $C_{\text{part.}}$ (%) | $N_{imp}$ |
|---|---|---|---|---|---|
| Munnink et al. [9] | SARS-CoV-2_61_LEFT | 67.2 | 98.9 | 99.0 | 9 |
| Munnink et al. [9] | SARS-CoV-2_61_RIGHT | 67.3 | 98.8 | 98.8 | 16 |
| Munnink et al. [9] | SARS-CoV-2_62_LEFT | 74.0 | 99.1 | 99.1 | 6 |
| Munnink et al. [9] | SARS-CoV-2_62_RIGHT | 70.0 | 99.3 | 99.3 | 1 |
| Munnink et al. [9] | SARS-CoV-2_63_LEFT | 69.5 | 99.1 | 99.1 | 4 |
| Munnink et al. [9] | SARS-CoV-2_63_RIGHT | 69.1 | 99.0 | 99.0 | 10 |
| Munnink et al. [9] | SARS-CoV-2_64_LEFT | 74.9 | 99.3 | 99.4 | 27 |
| Munnink et al. [9] | SARS-CoV-2_64_RIGHT | 72.4 | 98.4 | 98.7 | 48 |
| Munnink et al. [9] | SARS-CoV-2_65_RIGHT | 66.8 | 98.9 | 99.0 | 19 |
| Munnink et al. [9] | SARS-CoV-2_66_RIGHT | 66.6 | 98.9 | 98.9 | 4 |
| Munnink et al. [9] | SARS-CoV-2_67_LEFT | 71.8 | 99.0 | 99.2 | 56 |
| Munnink et al. [9] | SARS-CoV-2_67_RIGHT | 67.2 | 95.6 | 95.6 | 8 |
| Munnink et al. [9] | SARS-CoV-2_68_LEFT | 70.1 | 99.0 | 99.3 | 50 |
| Munnink et al. [9] | SARS-CoV-2_68_RIGHT | 69.7 | 99.3 | 99.3 | 20 |
| Munnink et al. [9] | SARS-CoV-2_69_LEFT | 68.3 | 99.2 | 99.3 | 16 |
| Munnink et al. [9] | SARS-CoV-2_69_RIGHT | 66.0 | 98.7 | 98.7 | 0 |
| Munnink et al. [9] | SARS-CoV-2_70_LEFT | 67.2 | 83.0 | 83.0 | 1 |
| Munnink et al. [9] | SARS-CoV-2_70_RIGHT | 68.6 | 99.4 | 99.4 | 4 |
| Munnink et al. [9] | SARS-CoV-2_71_LEFT | 66.9 | 99.4 | 99.4 | 1 |
| Munnink et al. [9] | SARS-CoV-2_71_RIGHT | 69.3 | 99.2 | 99.2 | 5 |
| Munnink et al. [9] | SARS-CoV-2_72_LEFT | 67.9 | 98.6 | 98.6 | 6 |

(continued on next page)

(Table S5, SARS-Cov-2 coverage, continued)

| Set | Primer/probe | $T_{\text{ref.}}$ (°C) | $C_{\text{strict}}$ (%) | $C_{\text{part.}}$ (%) | $N_{imp}$ |
|---|---|---|---|---|---|
| Munnink et al. [9] | SARS-CoV-2_72_RIGHT | 67.1 | 99.1 | 99.1 | 2 |
| Munnink et al. [9] | SARS-CoV-2_73_LEFT | 69.6 | 99.2 | 99.3 | 17 |
| Munnink et al. [9] | SARS-CoV-2_73_RIGHT | 66.3 | 99.3 | 99.4 | 3 |
| Munnink et al. [9] | SARS-CoV-2_74_LEFT | 68.7 | 99.2 | 99.3 | 7 |
| Munnink et al. [9] | SARS-CoV-2_74_RIGHT | 64.4 | 98.7 | 98.7 | 1 |
| Munnink et al. [9] | SARS-CoV-2_75_LEFT | 68.9 | 99.5 | 99.5 | 3 |
| Munnink et al. [9] | SARS-CoV-2_75_RIGHT | 69.8 | 99.2 | 99.3 | 23 |
| Munnink et al. [9] | SARS-CoV-2_76_LEFT | 70.7 | 99.4 | 99.5 | 17 |
| Munnink et al. [9] | SARS-CoV-2_76_RIGHT | 68.4 | 60.4 | 60.5 | 16 |
| Munnink et al. [9] | SARS-CoV-2_77_LEFT | 68.8 | 98.2 | 99.3 | 251 |
| Munnink et al. [9] | SARS-CoV-2_77_RIGHT | 67.0 | 97.2 | 97.6 | 89 |
| Munnink et al. [9] | SARS-CoV-2_78_LEFT | 69.3 | 98.7 | 99.1 | 86 |
| Munnink et al. [9] | SARS-CoV-2_78_RIGHT | 66.8 | 99.3 | 99.4 | 5 |
| Munnink et al. [9] | SARS-CoV-2_79_LEFT | 68.4 | 99.0 | 99.0 | 17 |
| Munnink et al. [9] | SARS-CoV-2_79_RIGHT | 68.7 | 98.7 | 99.0 | 66 |
| Munnink et al. [9] | SARS-CoV-2_80_LEFT | 68.9 | 99.1 | 99.3 | 36 |
| Munnink et al. [9] | SARS-CoV-2_80_RIGHT | 67.1 | 99.4 | 99.4 | 1 |
| Munnink et al. [9] | SARS-CoV-2_81_LEFT | 65.9 | 97.8 | 99.3 | 324 |
| Munnink et al. [9] | SARS-CoV-2_81_RIGHT | 66.9 | 99.3 | 99.3 | 11 |
| Munnink et al. [9] | SARS-CoV-2_82_LEFT | 68.9 | 99.4 | 99.4 | 0 |
| Munnink et al. [9] | SARS-CoV-2_82_RIGHT | 68.7 | 99.1 | 99.3 | 31 |

(continued on next page)

(Table S5, SARS-Cov-2 coverage, continued)

| Set | Primer/probe | $T_{\text{ref.}}$ (°C) | $C_{\text{strict}}$ (%) | $C_{\text{part.}}$ (%) | $N_{imp}$ |
|---|---|---|---|---|---|
| Munnink et al. [9] | SARS-CoV-2_83_LEFT | 70.0 | 99.3 | 99.4 | 21 |
| Munnink et al. [9] | SARS-CoV-2_83_RIGHT | 65.7 | 99.0 | 99.0 | 4 |
| Munnink et al. [9] | SARS-CoV-2_84_LEFT | 72.3 | 98.4 | 98.7 | 69 |
| Munnink et al. [9] | SARS-CoV-2_84_RIGHT | 64.6 | 99.0 | 99.0 | 0 |
| Munnink et al. [9] | SARS-CoV-2_85_LEFT | 69.8 | 97.5 | 99.1 | 340 |
| Munnink et al. [9] | SARS-CoV-2_85_RIGHT | 67.8 | 99.3 | 99.4 | 15 |
| Munnink et al. [9] | SARS-CoV-2_86_LEFT | 67.0 | 99.4 | 99.5 | 11 |
| Munnink et al. [9] | SARS-CoV-2_86_RIGHT | 68.3 | 97.7 | 98.7 | 217 |
| Munnink et al. [9] | SARS-CoV-2_87_LEFT | 69.1 | 99.0 | 99.4 | 77 |
| Munnink et al. [9] | SARS-CoV-2_87_RIGHT | 69.8 | 99.2 | 99.3 | 20 |
| Munnink et al. [9] | SARS-CoV-2_88_LEFT | 68.4 | 99.2 | 99.2 | 10 |
| Munnink et al. [9] | SARS-CoV-2_88_RIGHT | 68.5 | 97.4 | 98.3 | 182 |
| Munnink et al. [9] | SARS-CoV-2_89_LEFT | 66.9 | 98.3 | 98.6 | 60 |
| Munnink et al. [9] | SARS-CoV-2_89_RIGHT | 68.3 | 64.7 | 64.9 | 32 |
| Nalla et al. [10] | nCoV_2019-F | 66.3 | 99.4 | 99.4 | 1 |
| Nalla et al. [10] | nCoV_2019-P1 | 51.8 | 99.5 | 99.5 | 0 |
| Nalla et al. [10] | nCoV_2019-P2 | 54.8 | 0.000 | 0.000 | 0 |
| Nalla et al. [10] | nCoV_2019-R | 68.0 | 99.5 | 99.5 | 1 |
| Niu et al. [11] | E-F | 59.2 | 99.1 | 99.1 | 0 |
| Niu et al. [11] | E-P | 84.2 | 99.2 | 99.3 | 13 |
| Niu et al. [11] | E-R | 57.6 | 99.3 | 99.3 | 0 |

(continued on next page)

(Table S5, SARS-Cov-2 coverage, continued)

| Set | Primer/probe | $T_{\text{ref.}}$ (°C) | $C_{\text{strict}}$ (%) | $C_{\text{part.}}$ (%) | $N_{imp}$ |
|---|---|---|---|---|---|
| Niu et al. [11] | RdRp-F | 64.6 | 0.000 | 0.000 | 0 |
| Niu et al. [11] | RdRp-P | 60.8 | 99.4 | 99.4 | 0 |
| Niu et al. [11] | RdRp-R | 68.4 | 0.000 | 0.000 | 0 |
| Park et al. [12] | SARS-CoV-2_IBS_E2-F | 65.2 | 99.3 | 99.3 | 1 |
| Park et al. [12] | SARS-CoV-2_IBS_E2-R | 61.3 | 99.3 | 99.3 | 0 |
| Park et al. [12] | SARS-CoV-2_IBS_N1-F | 63.3 | 99.1 | 99.1 | 0 |
| Park et al. [12] | SARS-CoV-2_IBS_N1-R | 63.9 | 98.0 | 98.1 | 11 |
| Park et al. [12] | SARS-CoV-2_IBS_RdRP2-F | 63.8 | 99.3 | 99.3 | 2 |
| Park et al. [12] | SARS-CoV-2_IBS_RdRP2-R | 64.5 | 99.4 | 99.4 | 0 |
| Park et al. [12] | SARS-CoV-2_IBS_S2-F | 63.7 | 94.9 | 94.9 | 0 |
| Park et al. [12] | SARS-CoV-2_IBS_S2-R | 61.6 | 95.6 | 95.6 | 0 |
| Park et al. [12] | SARS-CoV-2_IBS_m_N1-F | 60.8 | 99.3 | 99.3 | 0 |
| Park et al. [12] | SARS-CoV-2_IBS_m_N1-R | 65.4 | 99.3 | 99.4 | 5 |
| Park et al. [12] | SARS-CoV-2_IBS_m_N2-F | 60.3 | 98.8 | 98.8 | 0 |
| Park et al. [12] | SARS-CoV-2_IBS_m_N2-R | 62.4 | 98.7 | 98.7 | 0 |
| Park et al. [12] | SARS-CoV-2_IBS_m_RdRP1-F | 62.7 | 99.3 | 99.3 | 6 |
| Park et al. [12] | SARS-CoV-2_IBS_m_RdRP1-R | 60.8 | 99.3 | 99.3 | 0 |
| Park et al. [12] | SARS-CoV-2_IBS_m_RdRP2-F | 62.9 | 99.3 | 99.3 | 0 |
| Park et al. [12] | SARS-CoV-2_IBS_m_RdRP2-R | 62.2 | 98.8 | 98.8 | 0 |
| Park et al. [12] | SARS-CoV-2_IBS_m_S1-F | 62.5 | 99.3 | 99.3 | 0 |
| Park et al. [12] | SARS-CoV-2_IBS_m_S1-R | 59.3 | 99.2 | 99.2 | 0 |

(continued on next page)

(Table S5, SARS-Cov-2 coverage, continued)

| Set | Primer/probe | $T_{ref.}$ (°C) | $C_{strict}$ (%) | $C_{part.}$ (%) | $N_{imp}$ |
|---|---|---|---|---|---|
| Park et al. [12] | SARS-CoV-2_IBS_m_S2-F | 61.4 | 99.3 | 99.3 | 0 |
| Park et al. [12] | SARS-CoV-2_IBS_m_S2-R | 60.3 | 99.5 | 99.5 | 0 |
| Rahman et al. [13] | M-F | 68.9 | 99.2 | 99.2 | 1 |
| Rahman et al. [13] | M-P | 70.9 | 98.7 | 99.4 | 140 |
| Rahman et al. [13] | M-R | 66.3 | 99.4 | 99.4 | 3 |
| Rahman et al. [13] | ORF1b-F | 65.5 | 98.7 | 98.7 | 3 |
| Rahman et al. [13] | ORF1b-P | 76.1 | 97.9 | 98.7 | 179 |
| Rahman et al. [13] | ORF1b-R | 64.2 | 97.6 | 97.6 | 0 |
| Toptan et al. [14] | M-475-F | 64.8 | 99.2 | 99.2 | 0 |
| Toptan et al. [14] | M-507-P | 63.5 | 98.8 | 98.8 | 2 |
| Toptan et al. [14] | M-574-R | 65.4 | 99.4 | 99.4 | 8 |
| Toptan et al. [14] | S-1850-F | 64.6 | 99.3 | 99.3 | 1 |
| Toptan et al. [14] | S-1884-P | 62.2 | 99.2 | 99.2 | 0 |
| Toptan et al. [14] | S-1949-R | 63.0 | 99.5 | 99.5 | 0 |
| Vogels et al. [15] | HKU-N-F | 58.6 | 99.1 | 99.1 | 0 |
| Vogels et al. [15] | HKU-N-P | 63.0 | 98.8 | 98.8 | 0 |
| Vogels et al. [15] | HKU-N-R | 60.3 | 98.9 | 98.9 | 0 |
| Vogels et al. [15] | HKU-ORF1-F1 | 61.7 | 0.000 | 0.000 | 0 |
| Vogels et al. [15] | HKU-ORF1-F2 | 64.6 | 0.000 | 0.000 | 0 |
| Vogels et al. [15] | HKU-ORF1-F3 | 58.7 | 99.2 | 99.2 | 0 |
| Vogels et al. [15] | HKU-ORF1-F4 | 61.7 | 0.000 | 0.000 | 0 |

(continued on next page)

(Table S5, SARS-Cov-2 coverage, continued)

| Set | Primer/probe | $T_{\text{ref.}}$ (°C) | $C_{\text{strict}}$ (%) | $C_{\text{part.}}$ (%) | $N_{imp}$ |
|---|---|---|---|---|---|
| Vogels et al. [15] | HKU-ORF1-P1 | 64.5 | 99.3 | 99.3 | 4 |
| Vogels et al. [15] | HKU-ORF1-P2 | 63.6 | 0.0462 | 0.0462 | 0 |
| Vogels et al. [15] | HKU-ORF1-R1 | 61.7 | 99.1 | 99.1 | 0 |
| Vogels et al. [15] | HKU-ORF1-R2 | 65.5 | 0.000 | 0.000 | 0 |
| WHO | NIID_WH-1_F24381 | 61.2 | 98.6 | 98.6 | 0 |
| WHO | NIID_WH-1_F501 | 70.3 | 97.8 | 99.1 | 291 |
| WHO | NIID_WH-1_F509 | 63.3 | 99.0 | 99.0 | 2 |
| WHO | NIID_WH-1_R24873 | 61.5 | 99.3 | 99.3 | 0 |
| WHO | NIID_WH-1_R854 | 61.7 | 98.0 | 98.0 | 0 |
| WHO | NIID_WH-1_R913 | 69.2 | 99.2 | 99.4 | 39 |
| WHO | NIID_WH-1_Seq_F24383 | 60.4 | 98.6 | 98.6 | 0 |
| WHO | NIID_WH-1_Seq_F519 | 58.8 | 97.9 | 97.9 | 0 |
| WHO | NIID_WH-1_Seq_R24865 | 60.1 | 99.3 | 99.3 | 0 |
| WHO | NIID_WH-1_Seq_R840 | 60.2 | 99.0 | 99.0 | 0 |
| WHO | WH-NICN-F | 64.4 | 99.2 | 99.2 | 0 |
| WHO | WH-NICN-P | 51.3 | 99.4 | 99.4 | 0 |
| WHO | WH-NICN-R | 64.1 | 99.0 | 99.0 | 2 |
| WHO | WuhanCoV-spk1-f | 65.4 | 98.9 | 98.9 | 6 |
| WHO | WuhanCoV-spk2-r | 64.6 | 99.3 | 99.3 | 0 |
| WHO | nCoV_IP2-12669Fw | 54.3 | 99.4 | 99.4 | 0 |
| WHO | nCoV_IP2-12696bProbe | 67.0 | 99.4 | 99.5 | 6 |

(continued on next page)

(Table S5, SARS-Cov-2 coverage, continued)

| Set | Primer/probe | $T_{\text{ref.}}$ (°C) | $C_{\text{strict}}$ (%) | $C_{\text{part.}}$ (%) | $N_{imp}$ |
|---|---|---|---|---|---|
| WHO | nCoV_IP2-12759Rv | 53.7 | 99.2 | 99.2 | 0 |
| WHO | nCoV_IP4-14059Fw | 54.8 | 99.4 | 99.4 | 0 |
| WHO | nCoV_IP4-14084Probe | 61.3 | 99.4 | 99.4 | 0 |
| WHO | nCoV_IP4-14146Rv | 54.8 | 99.2 | 99.2 | 0 |
| Yip et al. [16] | nsp2f | 63.4 | 99.2 | 99.3 | 2 |
| Yip et al. [16] | nsp2r | 61.4 | 99.1 | 99.1 | 0 |

Table S6: Primers and probes, and their respective coverages for SARS-CoV-1 genomes.

| Set | Primer/probe | $T_{\text{ref.}}$ (°C) | $C_{\text{strict}}$ (%) | $C_{\text{part.}}$ (%) | $N_{imp}$ |
|---|---|---|---|---|---|
| Bhadra et al. [1] | E-PF | 62.6 | 0.000 | 0.000 | 0 |
| Bhadra et al. [1] | E-Probe | 64.2 | 100. | 100. | 0 |
| Bhadra et al. [1] | E-RP | 61.4 | 100. | 100. | 0 |
| Bhadra et al. [1] | M-PF | 62.2 | 0.000 | 0.000 | 0 |
| Bhadra et al. [1] | M-Probe | 61.3 | 0.000 | 0.000 | 0 |
| Bhadra et al. [1] | M-RP | 63.3 | 100. | 100. | 0 |
| CDC | 2019-nCoV_N1-F | 62.4 | 0.000 | 0.000 | 0 |
| CDC | 2019-nCoV_N1-P | 75.5 | 0.000 | 0.000 | 0 |
| CDC | 2019-nCoV_N1-R | 69.0 | 0.000 | 0.000 | 0 |
| CDC | 2019-nCoV_N2-F | 61.1 | 80.0 | 80.0 | 0 |
| CDC | 2019-nCoV_N2-P | 74.7 | 0.000 | 0.000 | 0 |
| CDC | 2019-nCoV_N2-R | 63.3 | 0.000 | 0.000 | 0 |
| Corman et al. [2] | E_Sarbeco_F | 65.6 | 100. | 100. | 0 |
| Corman et al. [2] | E_Sarbeco_P1 | 78.2 | 100. | 100. | 0 |
| Corman et al. [2] | E_Sarbeco_R | 65.9 | 100. | 100. | 0 |
| Corman et al. [2] | N_Sarbeco_F | 64.6 | 90.0 | 90.0 | 0 |
| Corman et al. [2] | N_Sarbeco_P | 71.9 | 100. | 100. | 0 |
| Corman et al. [2] | N_Sarbeco_R | 65.1 | 0.000 | 100. | 10 |
| Corman et al. [2] | RdRp_SARSr-FA | 69.1 | 0.000 | 90.0 | 9 |

(continued on next page)

(Table S6, SARS-CoV-1 coverage, continued)

| Set | Primer/probe | $T_{\text{ref.}}$ (°C) | $C_{\text{strict}}$ (%) | $C_{\text{part.}}$ (%) | $N_{imp}$ |
|---|---|---|---|---|---|
| Corman et al. [2] | RdRp_SARSr-FG | 70.7 | 90.0 | 90.0 | 0 |
| Corman et al. [2] | RdRp_SARSr-P1-1 | 79.2 | 0.000 | 100. | 10 |
| Corman et al. [2] | RdRp_SARSr-P1-2 | 81.7 | 0.000 | 100. | 10 |
| Corman et al. [2] | RdRp_SARSr-P1-3 | 77.3 | 10.0 | 100. | 9 |
| Corman et al. [2] | RdRp_SARSr-P1-4 | 79.9 | 90.0 | 100. | 1 |
| Corman et al. [2] | RdRp_SARSr-P1-5 | 78.2 | 0.000 | 100. | 10 |
| Corman et al. [2] | RdRp_SARSr-P1-6 | 80.7 | 0.000 | 100. | 10 |
| Corman et al. [2] | RdRp_SARSr-P1-7 | 76.2 | 0.000 | 100. | 10 |
| Corman et al. [2] | RdRp_SARSr-P1-8 | 78.8 | 0.000 | 100. | 10 |
| Corman et al. [2] | RdRp_SARSr-P2 | 76.3 | 0.000 | 10.0 | 1 |
| Corman et al. [2] | RdRp_SARSr-R1 | 61.7 | 0.000 | 0.000 | 0 |
| Corman et al. [2] | RdRp_SARSr-R2 | 62.1 | 90.0 | 90.0 | 0 |
| Corman et al. [2] | RdRp_SARSr-R3 | 63.5 | 0.000 | 0.000 | 0 |
| Corman et al. [2] | RdRp_SARSr-R4 | 63.8 | 0.000 | 0.000 | 0 |
| Davda et al. [3] | MF1 | 56.5 | 0.000 | 0.000 | 0 |
| Davda et al. [3] | MF2 | 58.6 | 0.000 | 0.000 | 0 |
| Davda et al. [3] | MR1 | 58.9 | 0.000 | 0.000 | 0 |
| Davda et al. [3] | MR2 | 60.2 | 0.000 | 0.000 | 0 |
| Davda et al. [3] | N1F1 | 66.7 | 0.000 | 0.000 | 0 |
| Davda et al. [3] | N1F2 | 70.3 | 80.0 | 90.0 | 1 |
| Davda et al. [3] | N1R1 | 62.2 | 0.000 | 0.000 | 0 |

(continued on next page)

(Table S6, SARS-CoV-1 coverage, continued)

| Set | Primer/probe | $T_{\text{ref.}}$ (°C) | $C_{\text{strict}}$ (%) | $C_{\text{part.}}$ (%) | $N_{imp}$ |
|---|---|---|---|---|---|
| Davda et al. [3] | N1R2 | 68.0 | 0.000 | 0.000 | 0 |
| Davda et al. [3] | N2F1 | 60.3 | 0.000 | 0.000 | 0 |
| Davda et al. [3] | N2F2 | 57.6 | 0.000 | 0.000 | 0 |
| Davda et al. [3] | N2R1 | 59.9 | 0.000 | 0.000 | 0 |
| Davda et al. [3] | N2R2 | 57.0 | 0.000 | 0.000 | 0 |
| Davda et al. [3] | Orf1abF1 | 69.7 | 0.000 | 0.000 | 0 |
| Davda et al. [3] | Orf1abF2 | 63.2 | 0.000 | 0.000 | 0 |
| Davda et al. [3] | Orf1abR1 | 67.4 | 0.000 | 0.000 | 0 |
| Davda et al. [3] | Orf1abR2 | 61.7 | 0.000 | 0.000 | 0 |
| Grant et al. [4] | NgeneProbe | 79.9 | 90.0 | 100. | 1 |
| Grant et al. [4] | NgeneTaq1 | 62.4 | 100. | 100. | 0 |
| Grant et al. [4] | NgeneTaq2 | 63.8 | 0.000 | 0.000 | 0 |
| Hirotsu et al. [5] | NIID-N2-F | 60.6 | 0.000 | 0.000 | 0 |
| Hirotsu et al. [5] | NIID-N2-P | 68.8 | 0.000 | 0.000 | 0 |
| Hirotsu et al. [5] | NIID-N2-R | 63.9 | 0.000 | 0.000 | 0 |
| Lanza et al. [6] | UFRN_1_F | 63.7 | 0.000 | 0.000 | 0 |
| Lanza et al. [6] | UFRN_1_P | 75.0 | 0.000 | 0.000 | 0 |
| Lanza et al. [6] | UFRN_1_R | 62.9 | 0.000 | 0.000 | 0 |
| Lanza et al. [6] | UFRN_2_F | 62.2 | 0.000 | 0.000 | 0 |
| Lanza et al. [6] | UFRN_2_P | 69.7 | 0.000 | 0.000 | 0 |
| Lanza et al. [6] | UFRN_2_R | 62.0 | 0.000 | 0.000 | 0 |

(continued on next page)

(Table S6, SARS-CoV-1 coverage, continued)

| Set | Primer/probe | $T_{\text{ref.}}$ (°C) | $C_{\text{strict}}$ (%) | $C_{\text{part.}}$ (%) | $N_{imp}$ |
|---|---|---|---|---|---|
| Lanza et al. [6] | UFRN_3_F | 62.9 | 0.000 | 0.000 | 0 |
| Lanza et al. [6] | UFRN_3_P | 69.1 | 0.000 | 0.000 | 0 |
| Lanza et al. [6] | UFRN_3_R | 60.5 | 0.000 | 0.000 | 0 |
| Lanza et al. [6] | UFRN_4_F | 62.0 | 0.000 | 0.000 | 0 |
| Lanza et al. [6] | UFRN_4_P | 69.0 | 0.000 | 0.000 | 0 |
| Lanza et al. [6] | UFRN_4_R | 64.9 | 0.000 | 0.000 | 0 |
| Lanza et al. [6] | UFRN_5_F | 62.2 | 0.000 | 0.000 | 0 |
| Lanza et al. [6] | UFRN_5_P | 73.0 | 0.000 | 90.0 | 9 |
| Lanza et al. [6] | UFRN_5_R | 63.7 | 0.000 | 0.000 | 0 |
| Lanza et al. [6] | UFRN_6_F | 63.2 | 0.000 | 0.000 | 0 |
| Lanza et al. [6] | UFRN_6_P | 71.3 | 0.000 | 90.0 | 9 |
| Lanza et al. [6] | UFRN_6_R | 62.4 | 0.000 | 0.000 | 0 |
| Lanza et al. [6] | UFRN_7_F | 62.5 | 0.000 | 0.000 | 0 |
| Lanza et al. [6] | UFRN_7_P | 71.7 | 0.000 | 90.0 | 9 |
| Lanza et al. [6] | UFRN_7_R | 61.5 | 0.000 | 0.000 | 0 |
| Lanza et al. [6] | UFRN_8_F | 61.3 | 0.000 | 0.000 | 0 |
| Lanza et al. [6] | UFRN_8_P | 70.5 | 0.000 | 0.000 | 0 |
| Lanza et al. [6] | UFRN_8_R | 62.0 | 0.000 | 0.000 | 0 |
| Lanza et al. [6] | UFRN_9_F | 60.5 | 0.000 | 0.000 | 0 |
| Lanza et al. [6] | UFRN_9_P | 72.6 | 0.000 | 0.000 | 0 |
| Lanza et al. [6] | UFRN_9_R | 62.4 | 0.000 | 0.000 | 0 |

(continued on next page)

(Table S6, SARS-CoV-1 coverage, continued)

| Set | Primer/probe | $T_{\text{ref.}}$ (°C) | $C_{\text{strict}}$ (%) | $C_{\text{part.}}$ (%) | $N_{imp}$ |
|---|---|---|---|---|---|
| Li et al. [7] | S-1 | 70.4 | 0.000 | 100. | 10 |
| Li et al. [7] | S-2 | 67.5 | 0.000 | 0.000 | 0 |
| Lu et al. [8] | ORF1a-F | 64.0 | 0.000 | 0.000 | 0 |
| Lu et al. [8] | ORF1a-P | 74.7 | 0.000 | 0.000 | 0 |
| Lu et al. [8] | ORF1a-R | 68.8 | 0.000 | 0.000 | 0 |
| Luminex | China_N_F | 67.1 | 0.000 | 90.0 | 9 |
| Luminex | China_N_P | 63.1 | 0.000 | 0.000 | 0 |
| Luminex | China_N_R | 65.6 | 0.000 | 0.000 | 0 |
| Luminex | China_ORF_F | 59.4 | 0.000 | 0.000 | 0 |
| Luminex | China_ORF_P | 81.0 | 0.000 | 100. | 10 |
| Luminex | China_ORF_R | 61.5 | 0.000 | 0.000 | 0 |
| Munnink et al. [9] | SARS-CoV-2_1_LEFT | 67.9 | 0.000 | 0.000 | 0 |
| Munnink et al. [9] | SARS-CoV-2_1_RIGHT | 69.9 | 0.000 | 0.000 | 0 |
| Munnink et al. [9] | SARS-CoV-2_2_LEFT | 69.4 | 0.000 | 0.000 | 0 |
| Munnink et al. [9] | SARS-CoV-2_2_RIGHT | 67.7 | 0.000 | 0.000 | 0 |
| Munnink et al. [9] | SARS-CoV-2_3_LEFT | 68.7 | 0.000 | 90.0 | 9 |
| Munnink et al. [9] | SARS-CoV-2_3_RIGHT | 68.4 | 0.000 | 0.000 | 0 |
| Munnink et al. [9] | SARS-CoV-2_4_LEFT | 66.8 | 0.000 | 0.000 | 0 |
| Munnink et al. [9] | SARS-CoV-2_4_RIGHT | 66.6 | 0.000 | 0.000 | 0 |
| Munnink et al. [9] | SARS-CoV-2_5_LEFT | 67.7 | 0.000 | 0.000 | 0 |
| Munnink et al. [9] | SARS-CoV-2_5_RIGHT | 68.2 | 0.000 | 0.000 | 0 |

(continued on next page)

(Table S6, SARS-CoV-1 coverage, continued)

| Set | Primer/probe | $T_{\text{ref.}}$ (°C) | $C_{\text{strict}}$ (%) | $C_{\text{part.}}$ (%) | $N_{imp}$ |
|---|---|---|---|---|---|
| Munnink et al. [9] | SARS-CoV-2_6_LEFT | 73.2 | 0.000 | 0.000 | 0 |
| Munnink et al. [9] | SARS-CoV-2_6_RIGHT | 66.3 | 0.000 | 0.000 | 0 |
| Munnink et al. [9] | SARS-CoV-2_7_LEFT | 71.3 | 0.000 | 0.000 | 0 |
| Munnink et al. [9] | SARS-CoV-2_7_RIGHT | 67.8 | 0.000 | 0.000 | 0 |
| Munnink et al. [9] | SARS-CoV-2_8_LEFT | 67.2 | 0.000 | 0.000 | 0 |
| Munnink et al. [9] | SARS-CoV-2_8_RIGHT | 65.9 | 0.000 | 0.000 | 0 |
| Munnink et al. [9] | SARS-CoV-2_9_LEFT | 72.3 | 0.000 | 0.000 | 0 |
| Munnink et al. [9] | SARS-CoV-2_9_RIGHT | 69.0 | 0.000 | 0.000 | 0 |
| Munnink et al. [9] | SARS-CoV-2_11_LEFT | 66.9 | 0.000 | 0.000 | 0 |
| Munnink et al. [9] | SARS-CoV-2_11_RIGHT | 71.3 | 0.000 | 0.000 | 0 |
| Munnink et al. [9] | SARS-CoV-2_12_RIGHT | 67.2 | 0.000 | 0.000 | 0 |
| Munnink et al. [9] | SARS-CoV-2_13_LEFT | 70.0 | 0.000 | 0.000 | 0 |
| Munnink et al. [9] | SARS-CoV-2_13_RIGHT | 68.2 | 0.000 | 0.000 | 0 |
| Munnink et al. [9] | SARS-CoV-2_14_LEFT | 72.3 | 0.000 | 0.000 | 0 |
| Munnink et al. [9] | SARS-CoV-2_14_RIGHT | 68.1 | 0.000 | 0.000 | 0 |
| Munnink et al. [9] | SARS-CoV-2_15_LEFT | 70.9 | 0.000 | 0.000 | 0 |
| Munnink et al. [9] | SARS-CoV-2_15_RIGHT | 67.9 | 0.000 | 0.000 | 0 |
| Munnink et al. [9] | SARS-CoV-2_16_LEFT | 69.9 | 0.000 | 0.000 | 0 |
| Munnink et al. [9] | SARS-CoV-2_16_RIGHT | 65.4 | 0.000 | 0.000 | 0 |
| Munnink et al. [9] | SARS-CoV-2_17_LEFT | 66.7 | 0.000 | 0.000 | 0 |
| Munnink et al. [9] | SARS-CoV-2_17_RIGHT | 67.9 | 0.000 | 0.000 | 0 |

(continued on next page)

(Table S6, SARS-CoV-1 coverage, continued)

| Set | Primer/probe | $T_{\text{ref.}}$ (°C) | $C_{\text{strict}}$ (%) | $C_{\text{part.}}$ (%) | $N_{imp}$ |
|---|---|---|---|---|---|
| Munnink et al. [9] | SARS-CoV-2_18_LEFT | 68.0 | 0.000 | 0.000 | 0 |
| Munnink et al. [9] | SARS-CoV-2_18_RIGHT | 71.0 | 0.000 | 0.000 | 0 |
| Munnink et al. [9] | SARS-CoV-2_19_LEFT | 69.6 | 0.000 | 0.000 | 0 |
| Munnink et al. [9] | SARS-CoV-2_19_RIGHT | 69.6 | 0.000 | 0.000 | 0 |
| Munnink et al. [9] | SARS-CoV-2_20_LEFT | 67.0 | 0.000 | 0.000 | 0 |
| Munnink et al. [9] | SARS-CoV-2_21_LEFT | 72.1 | 0.000 | 0.000 | 0 |
| Munnink et al. [9] | SARS-CoV-2_21_RIGHT | 66.8 | 0.000 | 0.000 | 0 |
| Munnink et al. [9] | SARS-CoV-2_22_LEFT | 66.6 | 0.000 | 0.000 | 0 |
| Munnink et al. [9] | SARS-CoV-2_22_RIGHT | 67.7 | 0.000 | 0.000 | 0 |
| Munnink et al. [9] | SARS-CoV-2_23_LEFT | 70.0 | 0.000 | 0.000 | 0 |
| Munnink et al. [9] | SARS-CoV-2_23_RIGHT | 68.1 | 0.000 | 0.000 | 0 |
| Munnink et al. [9] | SARS-CoV-2_24_LEFT | 72.0 | 0.000 | 100. | 10 |
| Munnink et al. [9] | SARS-CoV-2_24_RIGHT | 67.5 | 0.000 | 0.000 | 0 |
| Munnink et al. [9] | SARS-CoV-2_25_LEFT | 71.7 | 0.000 | 90.0 | 9 |
| Munnink et al. [9] | SARS-CoV-2_25_RIGHT | 67.5 | 0.000 | 0.000 | 0 |
| Munnink et al. [9] | SARS-CoV-2_26_LEFT | 70.5 | 0.000 | 0.000 | 0 |
| Munnink et al. [9] | SARS-CoV-2_26_RIGHT | 67.5 | 0.000 | 0.000 | 0 |
| Munnink et al. [9] | SARS-CoV-2_27_LEFT | 68.0 | 0.000 | 0.000 | 0 |
| Munnink et al. [9] | SARS-CoV-2_27_RIGHT | 68.2 | 0.000 | 90.0 | 9 |
| Munnink et al. [9] | SARS-CoV-2_28_LEFT | 67.5 | 0.000 | 10.0 | 1 |
| Munnink et al. [9] | SARS-CoV-2_28_RIGHT | 67.4 | 0.000 | 0.000 | 0 |

(continued on next page)

(Table S6, SARS-CoV-1 coverage, continued)

| Set | Primer/probe | $T_{\text{ref.}}$ (°C) | $C_{\text{strict}}$ (%) | $C_{\text{part.}}$ (%) | $N_{imp}$ |
|---|---|---|---|---|---|
| Munnink et al. [9] | SARS-CoV-2_29_LEFT | 71.6 | 0.000 | 0.000 | 0 |
| Munnink et al. [9] | SARS-CoV-2_29_RIGHT | 69.1 | 0.000 | 0.000 | 0 |
| Munnink et al. [9] | SARS-CoV-2_30_LEFT | 72.3 | 0.000 | 0.000 | 0 |
| Munnink et al. [9] | SARS-CoV-2_30_RIGHT | 69.1 | 0.000 | 0.000 | 0 |
| Munnink et al. [9] | SARS-CoV-2_31_LEFT | 68.4 | 0.000 | 0.000 | 0 |
| Munnink et al. [9] | SARS-CoV-2_31_RIGHT | 67.4 | 0.000 | 0.000 | 0 |
| Munnink et al. [9] | SARS-CoV-2_32_LEFT | 68.1 | 0.000 | 0.000 | 0 |
| Munnink et al. [9] | SARS-CoV-2_32_RIGHT | 67.8 | 0.000 | 0.000 | 0 |
| Munnink et al. [9] | SARS-CoV-2_33_LEFT | 71.5 | 0.000 | 100. | 10 |
| Munnink et al. [9] | SARS-CoV-2_33_RIGHT | 66.9 | 0.000 | 0.000 | 0 |
| Munnink et al. [9] | SARS-CoV-2_34_LEFT | 68.6 | 0.000 | 0.000 | 0 |
| Munnink et al. [9] | SARS-CoV-2_34_RIGHT | 66.1 | 0.000 | 0.000 | 0 |
| Munnink et al. [9] | SARS-CoV-2_35_LEFT | 68.7 | 0.000 | 0.000 | 0 |
| Munnink et al. [9] | SARS-CoV-2_35_RIGHT | 68.7 | 0.000 | 0.000 | 0 |
| Munnink et al. [9] | SARS-CoV-2_36_LEFT | 70.5 | 0.000 | 10.0 | 1 |
| Munnink et al. [9] | SARS-CoV-2_36_RIGHT | 68.0 | 0.000 | 90.0 | 9 |
| Munnink et al. [9] | SARS-CoV-2_37_LEFT | 68.5 | 0.000 | 90.0 | 9 |
| Munnink et al. [9] | SARS-CoV-2_37_RIGHT | 67.6 | 0.000 | 0.000 | 0 |
| Munnink et al. [9] | SARS-CoV-2_38_LEFT | 68.5 | 90.0 | 100. | 1 |
| Munnink et al. [9] | SARS-CoV-2_38_RIGHT | 67.3 | 0.000 | 0.000 | 0 |
| Munnink et al. [9] | SARS-CoV-2_39_LEFT | 70.8 | 0.000 | 90.0 | 9 |

(continued on next page)

(Table S6, SARS-CoV-1 coverage, continued)

| Set | Primer/probe | $T_{\text{ref.}}$ (°C) | $C_{\text{strict}}$ (%) | $C_{\text{part.}}$ (%) | $N_{imp}$ |
|---|---|---|---|---|---|
| Munnink et al. [9] | SARS-CoV-2_39_RIGHT | 67.6 | 0.000 | 0.000 | 0 |
| Munnink et al. [9] | SARS-CoV-2_40_LEFT | 70.7 | 0.000 | 90.0 | 9 |
| Munnink et al. [9] | SARS-CoV-2_40_RIGHT | 67.9 | 0.000 | 0.000 | 0 |
| Munnink et al. [9] | SARS-CoV-2_41_LEFT | 70.2 | 0.000 | 100. | 10 |
| Munnink et al. [9] | SARS-CoV-2_41_RIGHT | 68.4 | 0.000 | 0.000 | 0 |
| Munnink et al. [9] | SARS-CoV-2_42_LEFT | 72.3 | 0.000 | 0.000 | 0 |
| Munnink et al. [9] | SARS-CoV-2_42_RIGHT | 68.8 | 0.000 | 0.000 | 0 |
| Munnink et al. [9] | SARS-CoV-2_43_LEFT | 69.1 | 0.000 | 0.000 | 0 |
| Munnink et al. [9] | SARS-CoV-2_43_RIGHT | 66.7 | 0.000 | 0.000 | 0 |
| Munnink et al. [9] | SARS-CoV-2_44_LEFT | 72.3 | 0.000 | 90.0 | 9 |
| Munnink et al. [9] | SARS-CoV-2_44_RIGHT | 65.7 | 100. | 100. | 0 |
| Munnink et al. [9] | SARS-CoV-2_45_LEFT | 69.7 | 0.000 | 100. | 10 |
| Munnink et al. [9] | SARS-CoV-2_45_RIGHT | 65.0 | 0.000 | 0.000 | 0 |
| Munnink et al. [9] | SARS-CoV-2_46_LEFT | 69.3 | 90.0 | 100. | 1 |
| Munnink et al. [9] | SARS-CoV-2_46_RIGHT | 67.3 | 0.000 | 0.000 | 0 |
| Munnink et al. [9] | SARS-CoV-2_47_LEFT | 64.8 | 0.000 | 0.000 | 0 |
| Munnink et al. [9] | SARS-CoV-2_47_RIGHT | 67.9 | 90.0 | 90.0 | 0 |
| Munnink et al. [9] | SARS-CoV-2_48_LEFT | 66.7 | 0.000 | 0.000 | 0 |
| Munnink et al. [9] | SARS-CoV-2_48_RIGHT | 67.3 | 0.000 | 0.000 | 0 |
| Munnink et al. [9] | SARS-CoV-2_49_LEFT | 71.4 | 0.000 | 90.0 | 9 |
| Munnink et al. [9] | SARS-CoV-2_49_RIGHT | 66.8 | 0.000 | 0.000 | 0 |

(continued on next page)

(Table S6, SARS-CoV-1 coverage, continued)

| Set | Primer/probe | $T_{\text{ref.}}$ (°C) | $C_{\text{strict}}$ (%) | $C_{\text{part.}}$ (%) | $N_{imp}$ |
|---|---|---|---|---|---|
| Munnink et al. [9] | SARS-CoV-2_50_LEFT | 73.8 | 0.000 | 100. | 10 |
| Munnink et al. [9] | SARS-CoV-2_50_RIGHT | 68.1 | 0.000 | 0.000 | 0 |
| Munnink et al. [9] | SARS-CoV-2_51_LEFT | 68.9 | 0.000 | 90.0 | 9 |
| Munnink et al. [9] | SARS-CoV-2_51_RIGHT | 67.8 | 0.000 | 0.000 | 0 |
| Munnink et al. [9] | SARS-CoV-2_52_LEFT | 71.3 | 0.000 | 100. | 10 |
| Munnink et al. [9] | SARS-CoV-2_52_RIGHT | 68.5 | 0.000 | 0.000 | 0 |
| Munnink et al. [9] | SARS-CoV-2_53_RIGHT | 67.3 | 0.000 | 0.000 | 0 |
| Munnink et al. [9] | SARS-CoV-2_54_LEFT | 72.2 | 0.000 | 100. | 10 |
| Munnink et al. [9] | SARS-CoV-2_54_RIGHT | 69.3 | 90.0 | 90.0 | 0 |
| Munnink et al. [9] | SARS-CoV-2_55_LEFT | 70.0 | 0.000 | 0.000 | 0 |
| Munnink et al. [9] | SARS-CoV-2_55_RIGHT | 68.1 | 0.000 | 0.000 | 0 |
| Munnink et al. [9] | SARS-CoV-2_56_LEFT | 68.9 | 0.000 | 0.000 | 0 |
| Munnink et al. [9] | SARS-CoV-2_56_RIGHT | 68.4 | 10.0 | 100. | 9 |
| Munnink et al. [9] | SARS-CoV-2_57_LEFT | 74.2 | 0.000 | 100. | 10 |
| Munnink et al. [9] | SARS-CoV-2_57_RIGHT | 66.7 | 0.000 | 0.000 | 0 |
| Munnink et al. [9] | SARS-CoV-2_58_LEFT | 73.8 | 0.000 | 100. | 10 |
| Munnink et al. [9] | SARS-CoV-2_58_RIGHT | 67.9 | 0.000 | 0.000 | 0 |
| Munnink et al. [9] | SARS-CoV-2_59_LEFT | 68.9 | 0.000 | 90.0 | 9 |
| Munnink et al. [9] | SARS-CoV-2_59_RIGHT | 67.0 | 0.000 | 0.000 | 0 |
| Munnink et al. [9] | SARS-CoV-2_60_LEFT | 71.0 | 0.000 | 0.000 | 0 |
| Munnink et al. [9] | SARS-CoV-2_60_RIGHT | 70.4 | 0.000 | 90.0 | 9 |

(continued on next page)

(Table S6, SARS-CoV-1 coverage, continued)

| Set | Primer/probe | $T_{\text{ref.}}$ (°C) | $C_{\text{strict}}$ (%) | $C_{\text{part.}}$ (%) | $N_{imp}$ |
|---|---|---|---|---|---|
| Munnink et al. [9] | SARS-CoV-2_61_LEFT | 67.2 | 0.000 | 0.000 | 0 |
| Munnink et al. [9] | SARS-CoV-2_61_RIGHT | 67.3 | 90.0 | 90.0 | 0 |
| Munnink et al. [9] | SARS-CoV-2_62_LEFT | 74.0 | 0.000 | 0.000 | 0 |
| Munnink et al. [9] | SARS-CoV-2_62_RIGHT | 70.0 | 0.000 | 100. | 10 |
| Munnink et al. [9] | SARS-CoV-2_63_LEFT | 69.5 | 0.000 | 0.000 | 0 |
| Munnink et al. [9] | SARS-CoV-2_63_RIGHT | 69.1 | 0.000 | 10.0 | 1 |
| Munnink et al. [9] | SARS-CoV-2_64_LEFT | 74.9 | 10.0 | 10.0 | 0 |
| Munnink et al. [9] | SARS-CoV-2_64_RIGHT | 72.4 | 0.000 | 0.000 | 0 |
| Munnink et al. [9] | SARS-CoV-2_65_RIGHT | 66.8 | 0.000 | 0.000 | 0 |
| Munnink et al. [9] | SARS-CoV-2_66_RIGHT | 66.6 | 0.000 | 0.000 | 0 |
| Munnink et al. [9] | SARS-CoV-2_67_LEFT | 71.8 | 0.000 | 0.000 | 0 |
| Munnink et al. [9] | SARS-CoV-2_67_RIGHT | 67.2 | 0.000 | 0.000 | 0 |
| Munnink et al. [9] | SARS-CoV-2_68_LEFT | 70.1 | 0.000 | 0.000 | 0 |
| Munnink et al. [9] | SARS-CoV-2_68_RIGHT | 69.7 | 0.000 | 0.000 | 0 |
| Munnink et al. [9] | SARS-CoV-2_69_LEFT | 68.3 | 0.000 | 0.000 | 0 |
| Munnink et al. [9] | SARS-CoV-2_69_RIGHT | 66.0 | 0.000 | 0.000 | 0 |
| Munnink et al. [9] | SARS-CoV-2_70_LEFT | 67.2 | 0.000 | 0.000 | 0 |
| Munnink et al. [9] | SARS-CoV-2_70_RIGHT | 68.6 | 0.000 | 0.000 | 0 |
| Munnink et al. [9] | SARS-CoV-2_71_LEFT | 66.9 | 0.000 | 0.000 | 0 |
| Munnink et al. [9] | SARS-CoV-2_71_RIGHT | 69.3 | 0.000 | 0.000 | 0 |
| Munnink et al. [9] | SARS-CoV-2_72_LEFT | 67.9 | 0.000 | 0.000 | 0 |

(continued on next page)

(Table S6, SARS-CoV-1 coverage, continued)

| Set | Primer/probe | $T_{\text{ref.}}$ (°C) | $C_{\text{strict}}$ (%) | $C_{\text{part.}}$ (%) | $N_{imp}$ |
|---|---|---|---|---|---|
| Munnink et al. [9] | SARS-CoV-2_72_RIGHT | 67.1 | 0.000 | 0.000 | 0 |
| Munnink et al. [9] | SARS-CoV-2_73_LEFT | 69.6 | 0.000 | 0.000 | 0 |
| Munnink et al. [9] | SARS-CoV-2_73_RIGHT | 66.3 | 0.000 | 0.000 | 0 |
| Munnink et al. [9] | SARS-CoV-2_74_LEFT | 68.7 | 0.000 | 0.000 | 0 |
| Munnink et al. [9] | SARS-CoV-2_74_RIGHT | 64.4 | 0.000 | 0.000 | 0 |
| Munnink et al. [9] | SARS-CoV-2_75_LEFT | 68.9 | 0.000 | 0.000 | 0 |
| Munnink et al. [9] | SARS-CoV-2_75_RIGHT | 69.8 | 0.000 | 0.000 | 0 |
| Munnink et al. [9] | SARS-CoV-2_76_LEFT | 70.7 | 0.000 | 0.000 | 0 |
| Munnink et al. [9] | SARS-CoV-2_76_RIGHT | 68.4 | 90.0 | 100. | 1 |
| Munnink et al. [9] | SARS-CoV-2_77_LEFT | 68.8 | 0.000 | 0.000 | 0 |
| Munnink et al. [9] | SARS-CoV-2_77_RIGHT | 67.0 | 0.000 | 0.000 | 0 |
| Munnink et al. [9] | SARS-CoV-2_78_LEFT | 69.3 | 0.000 | 0.000 | 0 |
| Munnink et al. [9] | SARS-CoV-2_78_RIGHT | 66.8 | 90.0 | 100. | 1 |
| Munnink et al. [9] | SARS-CoV-2_79_LEFT | 68.4 | 0.000 | 0.000 | 0 |
| Munnink et al. [9] | SARS-CoV-2_79_RIGHT | 68.7 | 0.000 | 0.000 | 0 |
| Munnink et al. [9] | SARS-CoV-2_80_LEFT | 68.9 | 0.000 | 0.000 | 0 |
| Munnink et al. [9] | SARS-CoV-2_80_RIGHT | 67.1 | 0.000 | 0.000 | 0 |
| Munnink et al. [9] | SARS-CoV-2_81_LEFT | 65.9 | 60.0 | 90.0 | 3 |
| Munnink et al. [9] | SARS-CoV-2_81_RIGHT | 66.9 | 0.000 | 0.000 | 0 |
| Munnink et al. [9] | SARS-CoV-2_82_LEFT | 68.9 | 0.000 | 90.0 | 9 |
| Munnink et al. [9] | SARS-CoV-2_82_RIGHT | 68.7 | 0.000 | 0.000 | 0 |

(continued on next page)

(Table S6, SARS-CoV-1 coverage, continued)

| Set | Primer/probe | $T_{\text{ref.}}$ (°C) | $C_{\text{strict}}$ (%) | $C_{\text{part.}}$ (%) | $N_{imp}$ |
|---|---|---|---|---|---|
| Munnink et al. [9] | SARS-CoV-2_83_LEFT | 70.0 | 0.000 | 0.000 | 0 |
| Munnink et al. [9] | SARS-CoV-2_83_RIGHT | 65.7 | 0.000 | 0.000 | 0 |
| Munnink et al. [9] | SARS-CoV-2_84_LEFT | 72.3 | 0.000 | 0.000 | 0 |
| Munnink et al. [9] | SARS-CoV-2_84_RIGHT | 64.6 | 0.000 | 0.000 | 0 |
| Munnink et al. [9] | SARS-CoV-2_85_LEFT | 69.8 | 0.000 | 0.000 | 0 |
| Munnink et al. [9] | SARS-CoV-2_85_RIGHT | 67.8 | 0.000 | 90.0 | 9 |
| Munnink et al. [9] | SARS-CoV-2_86_LEFT | 67.0 | 0.000 | 0.000 | 0 |
| Munnink et al. [9] | SARS-CoV-2_86_RIGHT | 68.3 | 0.000 | 0.000 | 0 |
| Munnink et al. [9] | SARS-CoV-2_87_LEFT | 69.1 | 0.000 | 10.0 | 1 |
| Munnink et al. [9] | SARS-CoV-2_87_RIGHT | 69.8 | 0.000 | 0.000 | 0 |
| Munnink et al. [9] | SARS-CoV-2_88_LEFT | 68.4 | 0.000 | 0.000 | 0 |
| Munnink et al. [9] | SARS-CoV-2_88_RIGHT | 68.5 | 0.000 | 0.000 | 0 |
| Munnink et al. [9] | SARS-CoV-2_89_LEFT | 66.9 | 0.000 | 0.000 | 0 |
| Munnink et al. [9] | SARS-CoV-2_89_RIGHT | 68.3 | 60.0 | 60.0 | 0 |
| Nalla et al. [10] | nCoV_2019-F | 66.3 | 0.000 | 0.000 | 0 |
| Nalla et al. [10] | nCoV_2019-P1 | 51.8 | 90.0 | 90.0 | 0 |
| Nalla et al. [10] | nCoV_2019-P2 | 54.8 | 0.000 | 0.000 | 0 |
| Nalla et al. [10] | nCoV_2019-R | 68.0 | 0.000 | 100. | 10 |
| Niu et al. [11] | E-F | 59.2 | 80.0 | 80.0 | 0 |
| Niu et al. [11] | E-P | 84.2 | 20.0 | 100. | 8 |
| Niu et al. [11] | E-R | 57.6 | 100. | 100. | 0 |

(continued on next page)

(Table S6, SARS-CoV-1 coverage, continued)

| Set | Primer/probe | $T_{\text{ref.}}$ (°C) | $C_{\text{strict}}$ (%) | $C_{\text{part.}}$ (%) | $N_{imp}$ |
|---|---|---|---|---|---|
| Niu et al. [11] | RdRp-F | 64.6 | 90.0 | 90.0 | 0 |
| Niu et al. [11] | RdRp-P | 60.8 | 90.0 | 90.0 | 0 |
| Niu et al. [11] | RdRp-R | 68.4 | 0.000 | 0.000 | 0 |
| Park et al. [12] | SARS-CoV-2_IBS_E2-F | 65.2 | 0.000 | 0.000 | 0 |
| Park et al. [12] | SARS-CoV-2_IBS_E2-R | 61.3 | 100. | 100. | 0 |
| Park et al. [12] | SARS-CoV-2_IBS_N1-F | 63.3 | 0.000 | 0.000 | 0 |
| Park et al. [12] | SARS-CoV-2_IBS_N1-R | 63.9 | 0.000 | 0.000 | 0 |
| Park et al. [12] | SARS-CoV-2_IBS_RdRP2-F | 63.8 | 90.0 | 90.0 | 0 |
| Park et al. [12] | SARS-CoV-2_IBS_RdRP2-R | 64.5 | 100. | 100. | 0 |
| Park et al. [12] | SARS-CoV-2_IBS_S2-F | 63.7 | 0.000 | 0.000 | 0 |
| Park et al. [12] | SARS-CoV-2_IBS_S2-R | 61.6 | 0.000 | 0.000 | 0 |
| Park et al. [12] | SARS-CoV-2_IBS_m_N1-F | 60.8 | 0.000 | 0.000 | 0 |
| Park et al. [12] | SARS-CoV-2_IBS_m_N1-R | 65.4 | 0.000 | 0.000 | 0 |
| Park et al. [12] | SARS-CoV-2_IBS_m_N2-F | 60.3 | 100. | 100. | 0 |
| Park et al. [12] | SARS-CoV-2_IBS_m_N2-R | 62.4 | 100. | 100. | 0 |
| Park et al. [12] | SARS-CoV-2_IBS_m_RdRP1-F | 62.7 | 0.000 | 0.000 | 0 |
| Park et al. [12] | SARS-CoV-2_IBS_m_RdRP1-R | 60.8 | 0.000 | 0.000 | 0 |
| Park et al. [12] | SARS-CoV-2_IBS_m_RdRP2-F | 62.9 | 0.000 | 0.000 | 0 |
| Park et al. [12] | SARS-CoV-2_IBS_m_RdRP2-R | 62.2 | 100. | 100. | 0 |
| Park et al. [12] | SARS-CoV-2_IBS_m_S1-F | 62.5 | 0.000 | 0.000 | 0 |
| Park et al. [12] | SARS-CoV-2_IBS_m_S1-R | 59.3 | 0.000 | 0.000 | 0 |

(continued on next page)

(Table S6, SARS-CoV-1 coverage, continued)

| Set | Primer/probe | $T_{\text{ref.}}$ (°C) | $C_{\text{strict}}$ (%) | $C_{\text{part.}}$ (%) | $N_{imp}$ |
|---|---|---|---|---|---|
| Park et al. [12] | SARS-CoV-2_IBS_m_S2-F | 61.4 | 0.000 | 0.000 | 0 |
| Park et al. [12] | SARS-CoV-2_IBS_m_S2-R | 60.3 | 0.000 | 0.000 | 0 |
| Rahman et al. [13] | M-F | 68.9 | 0.000 | 0.000 | 0 |
| Rahman et al. [13] | M-P | 70.9 | 0.000 | 100. | 10 |
| Rahman et al. [13] | M-R | 66.3 | 0.000 | 0.000 | 0 |
| Rahman et al. [13] | ORF1b-F | 65.5 | 0.000 | 0.000 | 0 |
| Rahman et al. [13] | ORF1b-P | 76.1 | 0.000 | 10.0 | 1 |
| Rahman et al. [13] | ORF1b-R | 64.2 | 0.000 | 0.000 | 0 |
| Toptan et al. [14] | M-475-F | 64.8 | 0.000 | 0.000 | 0 |
| Toptan et al. [14] | M-507-P | 63.5 | 0.000 | 0.000 | 0 |
| Toptan et al. [14] | M-574-R | 65.4 | 0.000 | 0.000 | 0 |
| Toptan et al. [14] | S-1850-F | 64.6 | 0.000 | 0.000 | 0 |
| Toptan et al. [14] | S-1884-P | 62.2 | 0.000 | 0.000 | 0 |
| Toptan et al. [14] | S-1949-R | 63.0 | 0.000 | 0.000 | 0 |
| Vogels et al. [15] | HKU-N-F | 58.6 | 100. | 100. | 0 |
| Vogels et al. [15] | HKU-N-P | 63.0 | 90.0 | 90.0 | 0 |
| Vogels et al. [15] | HKU-N-R | 60.3 | 100. | 100. | 0 |
| Vogels et al. [15] | HKU-ORF1-F1 | 61.7 | 0.000 | 0.000 | 0 |
| Vogels et al. [15] | HKU-ORF1-F2 | 64.6 | 90.0 | 90.0 | 0 |
| Vogels et al. [15] | HKU-ORF1-F3 | 58.7 | 10.0 | 10.0 | 0 |
| Vogels et al. [15] | HKU-ORF1-F4 | 61.7 | 0.000 | 0.000 | 0 |

(continued on next page)

(Table S6, SARS-CoV-1 coverage, continued)

| Set | Primer/probe | $T_{\text{ref.}}$ (°C) | $C_{\text{strict}}$ (%) | $C_{\text{part.}}$ (%) | $N_{imp}$ |
|---|---|---|---|---|---|
| Vogels et al. [15] | HKU-ORF1-P1 | 64.5 | 0.000 | 0.000 | 0 |
| Vogels et al. [15] | HKU-ORF1-P2 | 63.6 | 100. | 100. | 0 |
| Vogels et al. [15] | HKU-ORF1-R1 | 61.7 | 0.000 | 0.000 | 0 |
| Vogels et al. [15] | HKU-ORF1-R2 | 65.5 | 100. | 100. | 0 |
| WHO | NIID_WH-1_F24381 | 61.2 | 0.000 | 0.000 | 0 |
| WHO | NIID_WH-1_F501 | 70.3 | 0.000 | 0.000 | 0 |
| WHO | NIID_WH-1_F509 | 63.3 | 0.000 | 0.000 | 0 |
| WHO | NIID_WH-1_R24873 | 61.5 | 0.000 | 0.000 | 0 |
| WHO | NIID_WH-1_R854 | 61.7 | 0.000 | 0.000 | 0 |
| WHO | NIID_WH-1_R913 | 69.2 | 0.000 | 0.000 | 0 |
| WHO | NIID_WH-1_Seq_F24383 | 60.4 | 0.000 | 0.000 | 0 |
| WHO | NIID_WH-1_Seq_F519 | 58.8 | 0.000 | 0.000 | 0 |
| WHO | NIID_WH-1_Seq_R24865 | 60.1 | 0.000 | 0.000 | 0 |
| WHO | NIID_WH-1_Seq_R840 | 60.2 | 0.000 | 0.000 | 0 |
| WHO | WH-NICN-F | 64.4 | 0.000 | 0.000 | 0 |
| WHO | WH-NICN-P | 51.3 | 0.000 | 0.000 | 0 |
| WHO | WH-NICN-R | 64.1 | 0.000 | 0.000 | 0 |
| WHO | WuhanCoV-spk1-f | 65.4 | 0.000 | 0.000 | 0 |
| WHO | WuhanCoV-spk2-r | 64.6 | 0.000 | 0.000 | 0 |
| WHO | nCoV_IP2-12669Fw | 54.3 | 0.000 | 0.000 | 0 |
| WHO | nCoV_IP2-12696bProbe | 67.0 | 0.000 | 0.000 | 0 |

(continued on next page)

(Table S6, SARS-CoV-1 coverage, continued)

| Set | Primer/probe | $T_{\text{ref.}}$ (°C) | $C_{\text{strict}}$ (%) | $C_{\text{part.}}$ (%) | $N_{imp}$ |
|---|---|---|---|---|---|
| WHO | nCoV_IP2-12759Rv | 53.7 | 0.000 | 0.000 | 0 |
| WHO | nCoV_IP4-14059Fw | 54.8 | 0.000 | 0.000 | 0 |
| WHO | nCoV_IP4-14084Probe | 61.3 | 0.000 | 0.000 | 0 |
| WHO | nCoV_IP4-14146Rv | 54.8 | 0.000 | 0.000 | 0 |
| Yip et al. [16] | nsp2f | 63.4 | 0.000 | 0.000 | 0 |
| Yip et al. [16] | nsp2r | 61.4 | 0.000 | 0.000 | 0 |

Table S7: Primers and probes, and their respective coverages for non-SARS genomes.

| Set | Primer/probe | $T_{\text{ref.}}$ (°C) | $C_{\text{strict}}$ (%) | $C_{\text{part.}}$ (%) | $N_{imp}$ |
|---|---|---|---|---|---|
| Bhadra et al. [1] | E-PF | 62.6 | 0.000 | 0.000 | 0 |
| Bhadra et al. [1] | E-Probe | 64.2 | 0.000 | 0.000 | 0 |
| Bhadra et al. [1] | E-RP | 61.4 | 0.000 | 0.000 | 0 |
| Bhadra et al. [1] | M-PF | 62.2 | 0.000 | 0.000 | 0 |
| Bhadra et al. [1] | M-Probe | 61.3 | 0.000 | 0.000 | 0 |
| Bhadra et al. [1] | M-RP | 63.3 | 0.000 | 0.000 | 0 |
| CDC | 2019-nCoV_N1-F | 62.4 | 0.000 | 0.000 | 0 |
| CDC | 2019-nCoV_N1-P | 75.5 | 0.000 | 0.000 | 0 |
| CDC | 2019-nCoV_N1-R | 69.0 | 0.000 | 0.000 | 0 |
| CDC | 2019-nCoV_N2-F | 61.1 | 0.000 | 0.000 | 0 |
| CDC | 2019-nCoV_N2-P | 74.7 | 0.000 | 0.000 | 0 |
| CDC | 2019-nCoV_N2-R | 63.3 | 0.000 | 0.000 | 0 |
| Corman et al. [2] | E_Sarbeco_F | 65.6 | 0.000 | 0.000 | 0 |
| Corman et al. [2] | E_Sarbeco_P1 | 78.2 | 0.000 | 0.000 | 0 |
| Corman et al. [2] | E_Sarbeco_R | 65.9 | 0.000 | 0.000 | 0 |
| Corman et al. [2] | N_Sarbeco_F | 64.6 | 0.000 | 0.000 | 0 |
| Corman et al. [2] | N_Sarbeco_P | 71.9 | 0.000 | 0.000 | 0 |
| Corman et al. [2] | N_Sarbeco_R | 65.1 | 0.000 | 0.000 | 0 |
| Corman et al. [2] | RdRp_SARSr-FA | 69.1 | 0.000 | 0.000 | 0 |

(continued on next page)

(Table S7, non-SARS coverage, continued)

| Set | Primer/probe | $T_{\text{ref.}}$ (°C) | $C_{\text{strict}}$ (%) | $C_{\text{part.}}$ (%) | $N_{imp}$ |
|---|---|---|---|---|---|
| Corman et al. [2] | RdRp_SARSr-FG | 70.7 | 0.000 | 0.000 | 0 |
| Corman et al. [2] | RdRp_SARSr-P1-1 | 79.2 | 0.000 | 0.000 | 0 |
| Corman et al. [2] | RdRp_SARSr-P1-2 | 81.7 | 0.000 | 0.000 | 0 |
| Corman et al. [2] | RdRp_SARSr-P1-3 | 77.3 | 0.000 | 0.000 | 0 |
| Corman et al. [2] | RdRp_SARSr-P1-4 | 79.9 | 0.000 | 0.000 | 0 |
| Corman et al. [2] | RdRp_SARSr-P1-5 | 78.2 | 0.000 | 19.2 | 60 |
| Corman et al. [2] | RdRp_SARSr-P1-6 | 80.7 | 0.000 | 0.000 | 0 |
| Corman et al. [2] | RdRp_SARSr-P1-7 | 76.2 | 0.000 | 0.000 | 0 |
| Corman et al. [2] | RdRp_SARSr-P1-8 | 78.8 | 0.000 | 0.000 | 0 |
| Corman et al. [2] | RdRp_SARSr-P2 | 76.3 | 0.000 | 0.000 | 0 |
| Corman et al. [2] | RdRp_SARSr-R1 | 61.7 | 0.000 | 0.000 | 0 |
| Corman et al. [2] | RdRp_SARSr-R2 | 62.1 | 0.000 | 0.000 | 0 |
| Corman et al. [2] | RdRp_SARSr-R3 | 63.5 | 0.000 | 0.000 | 0 |
| Corman et al. [2] | RdRp_SARSr-R4 | 63.8 | 0.000 | 0.000 | 0 |
| Davda et al. [3] | MF1 | 56.5 | 0.000 | 0.000 | 0 |
| Davda et al. [3] | MF2 | 58.6 | 0.000 | 0.000 | 0 |
| Davda et al. [3] | MR1 | 58.9 | 0.000 | 0.000 | 0 |
| Davda et al. [3] | MR2 | 60.2 | 0.000 | 0.000 | 0 |
| Davda et al. [3] | N1F1 | 66.7 | 0.000 | 0.000 | 0 |
| Davda et al. [3] | N1F2 | 70.3 | 0.000 | 0.000 | 0 |
| Davda et al. [3] | N1R1 | 62.2 | 0.000 | 0.000 | 0 |

(continued on next page)

(Table S7, non-SARS coverage, continued)

| Set | Primer/probe | $T_{\text{ref.}}$ (°C) | $C_{\text{strict}}$ (%) | $C_{\text{part.}}$ (%) | $N_{imp}$ |
|---|---|---|---|---|---|
| Davda et al. [3] | N1R2 | 68.0 | 0.000 | 0.000 | 0 |
| Davda et al. [3] | N2F1 | 60.3 | 0.000 | 0.000 | 0 |
| Davda et al. [3] | N2F2 | 57.6 | 0.000 | 0.000 | 0 |
| Davda et al. [3] | N2R1 | 59.9 | 0.000 | 0.000 | 0 |
| Davda et al. [3] | N2R2 | 57.0 | 0.000 | 0.000 | 0 |
| Davda et al. [3] | Orf1abF1 | 69.7 | 0.000 | 0.000 | 0 |
| Davda et al. [3] | Orf1abF2 | 63.2 | 0.000 | 0.000 | 0 |
| Davda et al. [3] | Orf1abR1 | 67.4 | 0.000 | 0.000 | 0 |
| Davda et al. [3] | Orf1abR2 | 61.7 | 0.000 | 0.000 | 0 |
| Grant et al. [4] | NgeneProbe | 79.9 | 0.000 | 0.000 | 0 |
| Grant et al. [4] | NgeneTaq1 | 62.4 | 0.000 | 0.000 | 0 |
| Grant et al. [4] | NgeneTaq2 | 63.8 | 0.000 | 0.000 | 0 |
| Hirotsu et al. [5] | NIID-N2-F | 60.6 | 0.000 | 0.000 | 0 |
| Hirotsu et al. [5] | NIID-N2-P | 68.8 | 0.000 | 0.000 | 0 |
| Hirotsu et al. [5] | NIID-N2-R | 63.9 | 0.000 | 0.000 | 0 |
| Lanza et al. [6] | UFRN_1_F | 63.7 | 0.000 | 0.000 | 0 |
| Lanza et al. [6] | UFRN_1_P | 75.0 | 0.000 | 0.000 | 0 |
| Lanza et al. [6] | UFRN_1_R | 62.9 | 0.000 | 0.000 | 0 |
| Lanza et al. [6] | UFRN_2_F | 62.2 | 0.000 | 0.000 | 0 |
| Lanza et al. [6] | UFRN_2_P | 69.7 | 0.000 | 0.000 | 0 |
| Lanza et al. [6] | UFRN_2_R | 62.0 | 0.000 | 0.000 | 0 |

(continued on next page)

(Table S7, non-SARS coverage, continued)

| Set | Primer/probe | $T_{\text{ref.}}$ (°C) | $C_{\text{strict}}$ (%) | $C_{\text{part.}}$ (%) | $N_{imp}$ |
|---|---|---|---|---|---|
| Lanza et al. [6] | UFRN_3_F | 62.9 | 0.000 | 0.000 | 0 |
| Lanza et al. [6] | UFRN_3_P | 69.1 | 0.000 | 0.000 | 0 |
| Lanza et al. [6] | UFRN_3_R | 60.5 | 0.000 | 0.000 | 0 |
| Lanza et al. [6] | UFRN_4_F | 62.0 | 0.000 | 0.000 | 0 |
| Lanza et al. [6] | UFRN_4_P | 69.0 | 0.000 | 0.000 | 0 |
| Lanza et al. [6] | UFRN_4_R | 64.9 | 0.000 | 0.000 | 0 |
| Lanza et al. [6] | UFRN_5_F | 62.2 | 0.000 | 0.000 | 0 |
| Lanza et al. [6] | UFRN_5_P | 73.0 | 0.000 | 0.000 | 0 |
| Lanza et al. [6] | UFRN_5_R | 63.7 | 0.000 | 0.000 | 0 |
| Lanza et al. [6] | UFRN_6_F | 63.2 | 0.000 | 0.000 | 0 |
| Lanza et al. [6] | UFRN_6_P | 71.3 | 0.000 | 0.000 | 0 |
| Lanza et al. [6] | UFRN_6_R | 62.4 | 0.000 | 0.000 | 0 |
| Lanza et al. [6] | UFRN_7_F | 62.5 | 0.000 | 0.000 | 0 |
| Lanza et al. [6] | UFRN_7_P | 71.7 | 0.000 | 0.000 | 0 |
| Lanza et al. [6] | UFRN_7_R | 61.5 | 0.000 | 0.000 | 0 |
| Lanza et al. [6] | UFRN_8_F | 61.3 | 0.000 | 0.000 | 0 |
| Lanza et al. [6] | UFRN_8_P | 70.5 | 0.000 | 0.000 | 0 |
| Lanza et al. [6] | UFRN_8_R | 62.0 | 0.000 | 0.000 | 0 |
| Lanza et al. [6] | UFRN_9_F | 60.5 | 0.000 | 0.000 | 0 |
| Lanza et al. [6] | UFRN_9_P | 72.6 | 0.000 | 0.000 | 0 |
| Lanza et al. [6] | UFRN_9_R | 62.4 | 0.000 | 0.000 | 0 |

(continued on next page)

(Table S7, non-SARS coverage, continued)

| Set | Primer/probe | $T_{\text{ref.}}$ (°C) | $C_{\text{strict}}$ (%) | $C_{\text{part.}}$ (%) | $N_{imp}$ |
|---|---|---|---|---|---|
| Li et al. [7] | S-1 | 70.4 | 0.000 | 0.000 | 0 |
| Li et al. [7] | S-2 | 67.5 | 0.000 | 0.000 | 0 |
| Lu et al. [8] | ORF1a-F | 64.0 | 0.000 | 0.000 | 0 |
| Lu et al. [8] | ORF1a-P | 74.7 | 0.000 | 0.000 | 0 |
| Lu et al. [8] | ORF1a-R | 68.8 | 0.000 | 0.000 | 0 |
| Luminex | China_N_F | 67.1 | 0.000 | 0.000 | 0 |
| Luminex | China_N_P | 63.1 | 0.000 | 0.000 | 0 |
| Luminex | China_N_R | 65.6 | 0.000 | 0.000 | 0 |
| Luminex | China_ORF_F | 59.4 | 0.000 | 0.000 | 0 |
| Luminex | China_ORF_P | 81.0 | 0.000 | 0.000 | 0 |
| Luminex | China_ORF_R | 61.5 | 0.000 | 0.000 | 0 |
| Munnink et al. [9] | SARS-CoV-2_1_LEFT | 67.9 | 0.000 | 0.000 | 0 |
| Munnink et al. [9] | SARS-CoV-2_1_RIGHT | 69.9 | 0.000 | 0.000 | 0 |
| Munnink et al. [9] | SARS-CoV-2_2_LEFT | 69.4 | 0.000 | 0.000 | 0 |
| Munnink et al. [9] | SARS-CoV-2_2_RIGHT | 67.7 | 0.000 | 0.000 | 0 |
| Munnink et al. [9] | SARS-CoV-2_3_LEFT | 68.7 | 0.000 | 0.000 | 0 |
| Munnink et al. [9] | SARS-CoV-2_3_RIGHT | 68.4 | 0.000 | 0.000 | 0 |
| Munnink et al. [9] | SARS-CoV-2_4_LEFT | 66.8 | 0.000 | 0.000 | 0 |
| Munnink et al. [9] | SARS-CoV-2_4_RIGHT | 66.6 | 0.000 | 0.000 | 0 |
| Munnink et al. [9] | SARS-CoV-2_5_LEFT | 67.7 | 0.000 | 0.000 | 0 |
| Munnink et al. [9] | SARS-CoV-2_5_RIGHT | 68.2 | 0.000 | 0.000 | 0 |

(continued on next page)

(Table S7, non-SARS coverage, continued)

| Set | Primer/probe | $T_{\mathrm{ref.}}$ (°C) | $C_{\mathrm{strict}}$ (%) | $C_{\mathrm{part.}}$ (%) | $N_{imp}$ |
|---|---|---|---|---|---|
| Munnink et al. [9] | SARS-CoV-2_6_LEFT | 73.2 | 0.000 | 0.000 | 0 |
| Munnink et al. [9] | SARS-CoV-2_6_RIGHT | 66.3 | 0.000 | 0.000 | 0 |
| Munnink et al. [9] | SARS-CoV-2_7_LEFT | 71.3 | 0.000 | 0.000 | 0 |
| Munnink et al. [9] | SARS-CoV-2_7_RIGHT | 67.8 | 0.000 | 0.000 | 0 |
| Munnink et al. [9] | SARS-CoV-2_8_LEFT | 67.2 | 0.000 | 0.000 | 0 |
| Munnink et al. [9] | SARS-CoV-2_8_RIGHT | 65.9 | 0.000 | 0.000 | 0 |
| Munnink et al. [9] | SARS-CoV-2_9_LEFT | 72.3 | 0.000 | 0.000 | 0 |
| Munnink et al. [9] | SARS-CoV-2_9_RIGHT | 69.0 | 0.000 | 0.000 | 0 |
| Munnink et al. [9] | SARS-CoV-2_11_LEFT | 66.9 | 0.000 | 0.000 | 0 |
| Munnink et al. [9] | SARS-CoV-2_11_RIGHT | 71.3 | 0.000 | 0.000 | 0 |
| Munnink et al. [9] | SARS-CoV-2_12_RIGHT | 67.2 | 0.000 | 0.000 | 0 |
| Munnink et al. [9] | SARS-CoV-2_13_LEFT | 70.0 | 0.000 | 0.000 | 0 |
| Munnink et al. [9] | SARS-CoV-2_13_RIGHT | 68.2 | 0.000 | 0.000 | 0 |
| Munnink et al. [9] | SARS-CoV-2_14_LEFT | 72.3 | 0.000 | 0.000 | 0 |
| Munnink et al. [9] | SARS-CoV-2_14_RIGHT | 68.1 | 0.000 | 0.000 | 0 |
| Munnink et al. [9] | SARS-CoV-2_15_LEFT | 70.9 | 0.000 | 0.000 | 0 |
| Munnink et al. [9] | SARS-CoV-2_15_RIGHT | 67.9 | 0.000 | 0.000 | 0 |
| Munnink et al. [9] | SARS-CoV-2_16_LEFT | 69.9 | 0.000 | 0.000 | 0 |
| Munnink et al. [9] | SARS-CoV-2_16_RIGHT | 65.4 | 0.000 | 0.000 | 0 |
| Munnink et al. [9] | SARS-CoV-2_17_LEFT | 66.7 | 0.000 | 0.000 | 0 |
| Munnink et al. [9] | SARS-CoV-2_17_RIGHT | 67.9 | 0.000 | 0.000 | 0 |

(continued on next page)

(Table S7, non-SARS coverage, continued)

| Set | Primer/probe | $T_{\text{ref.}}$ (°C) | $C_{\text{strict}}$ (%) | $C_{\text{part.}}$ (%) | $N_{imp}$ |
|---|---|---|---|---|---|
| Munnink et al. [9] | SARS-CoV-2_18_LEFT | 68.0 | 0.000 | 0.000 | 0 |
| Munnink et al. [9] | SARS-CoV-2_18_RIGHT | 71.0 | 0.000 | 0.000 | 0 |
| Munnink et al. [9] | SARS-CoV-2_19_LEFT | 69.6 | 0.000 | 0.000 | 0 |
| Munnink et al. [9] | SARS-CoV-2_19_RIGHT | 69.6 | 0.000 | 0.000 | 0 |
| Munnink et al. [9] | SARS-CoV-2_20_LEFT | 67.0 | 0.000 | 0.000 | 0 |
| Munnink et al. [9] | SARS-CoV-2_21_LEFT | 72.1 | 0.000 | 0.000 | 0 |
| Munnink et al. [9] | SARS-CoV-2_21_RIGHT | 66.8 | 0.000 | 0.000 | 0 |
| Munnink et al. [9] | SARS-CoV-2_22_LEFT | 66.6 | 0.000 | 0.000 | 0 |
| Munnink et al. [9] | SARS-CoV-2_22_RIGHT | 67.7 | 0.000 | 0.000 | 0 |
| Munnink et al. [9] | SARS-CoV-2_23_LEFT | 70.0 | 0.000 | 0.000 | 0 |
| Munnink et al. [9] | SARS-CoV-2_23_RIGHT | 68.1 | 0.000 | 0.000 | 0 |
| Munnink et al. [9] | SARS-CoV-2_24_LEFT | 72.0 | 0.000 | 0.000 | 0 |
| Munnink et al. [9] | SARS-CoV-2_24_RIGHT | 67.5 | 0.000 | 0.000 | 0 |
| Munnink et al. [9] | SARS-CoV-2_25_LEFT | 71.7 | 0.000 | 0.000 | 0 |
| Munnink et al. [9] | SARS-CoV-2_25_RIGHT | 67.5 | 0.000 | 0.000 | 0 |
| Munnink et al. [9] | SARS-CoV-2_26_LEFT | 70.5 | 0.000 | 0.000 | 0 |
| Munnink et al. [9] | SARS-CoV-2_26_RIGHT | 67.5 | 0.000 | 0.000 | 0 |
| Munnink et al. [9] | SARS-CoV-2_27_LEFT | 68.0 | 0.000 | 0.000 | 0 |
| Munnink et al. [9] | SARS-CoV-2_27_RIGHT | 68.2 | 0.000 | 0.000 | 0 |
| Munnink et al. [9] | SARS-CoV-2_28_LEFT | 67.5 | 0.000 | 0.000 | 0 |
| Munnink et al. [9] | SARS-CoV-2_28_RIGHT | 67.4 | 0.000 | 0.000 | 0 |

(continued on next page)

(Table S7, non-SARS coverage, continued)

| Set | Primer/probe | $T_{\text{ref.}}$ (°C) | $C_{\text{strict}}$ (%) | $C_{\text{part.}}$ (%) | $N_{imp}$ |
|---|---|---|---|---|---|
| Munnink et al. [9] | SARS-CoV-2_29_LEFT | 71.6 | 0.000 | 0.000 | 0 |
| Munnink et al. [9] | SARS-CoV-2_29_RIGHT | 69.1 | 0.000 | 0.000 | 0 |
| Munnink et al. [9] | SARS-CoV-2_30_LEFT | 72.3 | 0.000 | 0.000 | 0 |
| Munnink et al. [9] | SARS-CoV-2_30_RIGHT | 69.1 | 0.000 | 0.000 | 0 |
| Munnink et al. [9] | SARS-CoV-2_31_LEFT | 68.4 | 0.000 | 0.000 | 0 |
| Munnink et al. [9] | SARS-CoV-2_31_RIGHT | 67.4 | 0.000 | 0.000 | 0 |
| Munnink et al. [9] | SARS-CoV-2_32_LEFT | 68.1 | 0.000 | 0.000 | 0 |
| Munnink et al. [9] | SARS-CoV-2_32_RIGHT | 67.8 | 0.000 | 0.000 | 0 |
| Munnink et al. [9] | SARS-CoV-2_33_LEFT | 71.5 | 0.000 | 0.000 | 0 |
| Munnink et al. [9] | SARS-CoV-2_33_RIGHT | 66.9 | 0.000 | 0.000 | 0 |
| Munnink et al. [9] | SARS-CoV-2_34_LEFT | 68.6 | 0.000 | 0.000 | 0 |
| Munnink et al. [9] | SARS-CoV-2_34_RIGHT | 66.1 | 0.000 | 0.000 | 0 |
| Munnink et al. [9] | SARS-CoV-2_35_LEFT | 68.7 | 0.000 | 0.000 | 0 |
| Munnink et al. [9] | SARS-CoV-2_35_RIGHT | 68.7 | 0.000 | 0.000 | 0 |
| Munnink et al. [9] | SARS-CoV-2_36_LEFT | 70.5 | 0.000 | 0.000 | 0 |
| Munnink et al. [9] | SARS-CoV-2_36_RIGHT | 68.0 | 0.000 | 0.000 | 0 |
| Munnink et al. [9] | SARS-CoV-2_37_LEFT | 68.5 | 0.000 | 0.000 | 0 |
| Munnink et al. [9] | SARS-CoV-2_37_RIGHT | 67.6 | 0.000 | 0.000 | 0 |
| Munnink et al. [9] | SARS-CoV-2_38_LEFT | 68.5 | 0.000 | 0.000 | 0 |
| Munnink et al. [9] | SARS-CoV-2_38_RIGHT | 67.3 | 0.000 | 0.000 | 0 |
| Munnink et al. [9] | SARS-CoV-2_39_LEFT | 70.8 | 0.000 | 0.000 | 0 |

(continued on next page)

(Table S7, non-SARS coverage, continued)

| Set | Primer/probe | $T_{\text{ref.}}$ (°C) | $C_{\text{strict}}$ (%) | $C_{\text{part.}}$ (%) | $N_{imp}$ |
|---|---|---|---|---|---|
| Munnink et al. [9] | SARS-CoV-2_39_RIGHT | 67.6 | 0.000 | 0.000 | 0 |
| Munnink et al. [9] | SARS-CoV-2_40_LEFT | 70.7 | 0.000 | 0.000 | 0 |
| Munnink et al. [9] | SARS-CoV-2_40_RIGHT | 67.9 | 0.000 | 0.000 | 0 |
| Munnink et al. [9] | SARS-CoV-2_41_LEFT | 70.2 | 0.000 | 0.000 | 0 |
| Munnink et al. [9] | SARS-CoV-2_41_RIGHT | 68.4 | 0.000 | 0.000 | 0 |
| Munnink et al. [9] | SARS-CoV-2_42_LEFT | 72.3 | 0.000 | 0.000 | 0 |
| Munnink et al. [9] | SARS-CoV-2_42_RIGHT | 68.8 | 0.000 | 0.000 | 0 |
| Munnink et al. [9] | SARS-CoV-2_43_LEFT | 69.1 | 0.000 | 0.000 | 0 |
| Munnink et al. [9] | SARS-CoV-2_43_RIGHT | 66.7 | 0.000 | 0.000 | 0 |
| Munnink et al. [9] | SARS-CoV-2_44_LEFT | 72.3 | 0.000 | 0.000 | 0 |
| Munnink et al. [9] | SARS-CoV-2_44_RIGHT | 65.7 | 0.000 | 0.639 | 2 |
| Munnink et al. [9] | SARS-CoV-2_45_LEFT | 69.7 | 0.000 | 0.000 | 0 |
| Munnink et al. [9] | SARS-CoV-2_45_RIGHT | 65.0 | 0.000 | 0.000 | 0 |
| Munnink et al. [9] | SARS-CoV-2_46_LEFT | 69.3 | 0.000 | 0.000 | 0 |
| Munnink et al. [9] | SARS-CoV-2_46_RIGHT | 67.3 | 0.000 | 0.000 | 0 |
| Munnink et al. [9] | SARS-CoV-2_47_LEFT | 64.8 | 0.000 | 0.000 | 0 |
| Munnink et al. [9] | SARS-CoV-2_47_RIGHT | 67.9 | 0.000 | 0.000 | 0 |
| Munnink et al. [9] | SARS-CoV-2_48_LEFT | 66.7 | 0.000 | 0.000 | 0 |
| Munnink et al. [9] | SARS-CoV-2_48_RIGHT | 67.3 | 0.000 | 0.000 | 0 |
| Munnink et al. [9] | SARS-CoV-2_49_LEFT | 71.4 | 0.000 | 0.000 | 0 |
| Munnink et al. [9] | SARS-CoV-2_49_RIGHT | 66.8 | 0.000 | 0.000 | 0 |

(continued on next page)

(Table S7, non-SARS coverage, continued)

| Set | Primer/probe | $T_{\text{ref.}}$ (°C) | $C_{\text{strict}}$ (%) | $C_{\text{part.}}$ (%) | $N_{imp}$ |
|---|---|---|---|---|---|
| Munnink et al. [9] | SARS-CoV-2_50_LEFT | 73.8 | 0.000 | 0.000 | 0 |
| Munnink et al. [9] | SARS-CoV-2_50_RIGHT | 68.1 | 0.000 | 0.000 | 0 |
| Munnink et al. [9] | SARS-CoV-2_51_LEFT | 68.9 | 0.000 | 0.000 | 0 |
| Munnink et al. [9] | SARS-CoV-2_51_RIGHT | 67.8 | 0.000 | 0.000 | 0 |
| Munnink et al. [9] | SARS-CoV-2_52_LEFT | 71.3 | 0.000 | 0.000 | 0 |
| Munnink et al. [9] | SARS-CoV-2_52_RIGHT | 68.5 | 0.000 | 0.000 | 0 |
| Munnink et al. [9] | SARS-CoV-2_53_RIGHT | 67.3 | 0.000 | 0.000 | 0 |
| Munnink et al. [9] | SARS-CoV-2_54_LEFT | 72.2 | 0.000 | 0.000 | 0 |
| Munnink et al. [9] | SARS-CoV-2_54_RIGHT | 69.3 | 0.000 | 0.000 | 0 |
| Munnink et al. [9] | SARS-CoV-2_55_LEFT | 70.0 | 0.000 | 0.000 | 0 |
| Munnink et al. [9] | SARS-CoV-2_55_RIGHT | 68.1 | 0.000 | 0.000 | 0 |
| Munnink et al. [9] | SARS-CoV-2_56_LEFT | 68.9 | 0.000 | 0.000 | 0 |
| Munnink et al. [9] | SARS-CoV-2_56_RIGHT | 68.4 | 0.000 | 0.000 | 0 |
| Munnink et al. [9] | SARS-CoV-2_57_LEFT | 74.2 | 0.000 | 0.000 | 0 |
| Munnink et al. [9] | SARS-CoV-2_57_RIGHT | 66.7 | 0.000 | 0.000 | 0 |
| Munnink et al. [9] | SARS-CoV-2_58_LEFT | 73.8 | 0.000 | 0.000 | 0 |
| Munnink et al. [9] | SARS-CoV-2_58_RIGHT | 67.9 | 0.000 | 0.000 | 0 |
| Munnink et al. [9] | SARS-CoV-2_59_LEFT | 68.9 | 0.000 | 0.000 | 0 |
| Munnink et al. [9] | SARS-CoV-2_59_RIGHT | 67.0 | 0.000 | 0.000 | 0 |
| Munnink et al. [9] | SARS-CoV-2_60_LEFT | 71.0 | 0.000 | 0.000 | 0 |
| Munnink et al. [9] | SARS-CoV-2_60_RIGHT | 70.4 | 0.000 | 0.000 | 0 |

(continued on next page)

(Table S7, non-SARS coverage, continued)

| Set | Primer/probe | $T_{\text{ref.}}$ (°C) | $C_{\text{strict}}$ (%) | $C_{\text{part.}}$ (%) | $N_{imp}$ |
|---|---|---|---|---|---|
| Munnink et al. [9] | SARS-CoV-2_61_LEFT | 67.2 | 0.000 | 0.000 | 0 |
| Munnink et al. [9] | SARS-CoV-2_61_RIGHT | 67.3 | 0.000 | 0.000 | 0 |
| Munnink et al. [9] | SARS-CoV-2_62_LEFT | 74.0 | 0.000 | 0.000 | 0 |
| Munnink et al. [9] | SARS-CoV-2_62_RIGHT | 70.0 | 0.000 | 0.000 | 0 |
| Munnink et al. [9] | SARS-CoV-2_63_LEFT | 69.5 | 0.000 | 0.000 | 0 |
| Munnink et al. [9] | SARS-CoV-2_63_RIGHT | 69.1 | 0.000 | 0.000 | 0 |
| Munnink et al. [9] | SARS-CoV-2_64_LEFT | 74.9 | 0.000 | 0.000 | 0 |
| Munnink et al. [9] | SARS-CoV-2_64_RIGHT | 72.4 | 0.000 | 0.000 | 0 |
| Munnink et al. [9] | SARS-CoV-2_65_RIGHT | 66.8 | 0.000 | 0.000 | 0 |
| Munnink et al. [9] | SARS-CoV-2_66_RIGHT | 66.6 | 0.000 | 0.000 | 0 |
| Munnink et al. [9] | SARS-CoV-2_67_LEFT | 71.8 | 0.000 | 0.000 | 0 |
| Munnink et al. [9] | SARS-CoV-2_67_RIGHT | 67.2 | 0.000 | 0.000 | 0 |
| Munnink et al. [9] | SARS-CoV-2_68_LEFT | 70.1 | 0.000 | 0.000 | 0 |
| Munnink et al. [9] | SARS-CoV-2_68_RIGHT | 69.7 | 0.000 | 0.000 | 0 |
| Munnink et al. [9] | SARS-CoV-2_69_LEFT | 68.3 | 0.000 | 0.000 | 0 |
| Munnink et al. [9] | SARS-CoV-2_69_RIGHT | 66.0 | 0.000 | 0.000 | 0 |
| Munnink et al. [9] | SARS-CoV-2_70_LEFT | 67.2 | 0.000 | 0.000 | 0 |
| Munnink et al. [9] | SARS-CoV-2_70_RIGHT | 68.6 | 0.000 | 0.000 | 0 |
| Munnink et al. [9] | SARS-CoV-2_71_LEFT | 66.9 | 0.000 | 0.000 | 0 |
| Munnink et al. [9] | SARS-CoV-2_71_RIGHT | 69.3 | 0.000 | 0.000 | 0 |
| Munnink et al. [9] | SARS-CoV-2_72_LEFT | 67.9 | 0.000 | 0.000 | 0 |

(continued on next page)

(Table S7, non-SARS coverage, continued)

| Set | Primer/probe | $T_{\text{ref.}}$ (°C) | $C_{\text{strict}}$ (%) | $C_{\text{part.}}$ (%) | $N_{imp}$ |
|---|---|---|---|---|---|
| Munnink et al. [9] | SARS-CoV-2_72_RIGHT | 67.1 | 0.000 | 0.000 | 0 |
| Munnink et al. [9] | SARS-CoV-2_73_LEFT | 69.6 | 0.000 | 0.000 | 0 |
| Munnink et al. [9] | SARS-CoV-2_73_RIGHT | 66.3 | 0.000 | 0.000 | 0 |
| Munnink et al. [9] | SARS-CoV-2_74_LEFT | 68.7 | 0.000 | 0.000 | 0 |
| Munnink et al. [9] | SARS-CoV-2_74_RIGHT | 64.4 | 0.000 | 0.000 | 0 |
| Munnink et al. [9] | SARS-CoV-2_75_LEFT | 68.9 | 0.000 | 0.000 | 0 |
| Munnink et al. [9] | SARS-CoV-2_75_RIGHT | 69.8 | 0.000 | 0.000 | 0 |
| Munnink et al. [9] | SARS-CoV-2_76_LEFT | 70.7 | 0.000 | 0.000 | 0 |
| Munnink et al. [9] | SARS-CoV-2_76_RIGHT | 68.4 | 0.000 | 0.000 | 0 |
| Munnink et al. [9] | SARS-CoV-2_77_LEFT | 68.8 | 0.000 | 0.000 | 0 |
| Munnink et al. [9] | SARS-CoV-2_77_RIGHT | 67.0 | 0.000 | 0.000 | 0 |
| Munnink et al. [9] | SARS-CoV-2_78_LEFT | 69.3 | 0.000 | 0.000 | 0 |
| Munnink et al. [9] | SARS-CoV-2_78_RIGHT | 66.8 | 0.000 | 0.000 | 0 |
| Munnink et al. [9] | SARS-CoV-2_79_LEFT | 68.4 | 0.000 | 0.000 | 0 |
| Munnink et al. [9] | SARS-CoV-2_79_RIGHT | 68.7 | 0.000 | 0.000 | 0 |
| Munnink et al. [9] | SARS-CoV-2_80_LEFT | 68.9 | 0.000 | 0.000 | 0 |
| Munnink et al. [9] | SARS-CoV-2_80_RIGHT | 67.1 | 0.000 | 0.000 | 0 |
| Munnink et al. [9] | SARS-CoV-2_81_LEFT | 65.9 | 0.000 | 0.000 | 0 |
| Munnink et al. [9] | SARS-CoV-2_81_RIGHT | 66.9 | 0.000 | 0.000 | 0 |
| Munnink et al. [9] | SARS-CoV-2_82_LEFT | 68.9 | 0.000 | 0.000 | 0 |
| Munnink et al. [9] | SARS-CoV-2_82_RIGHT | 68.7 | 0.000 | 0.000 | 0 |

(continued on next page)

(Table S7, non-SARS coverage, continued)

| Set | Primer/probe | $T_{\mathrm{ref.}}$ (°C) | $C_{\mathrm{strict}}$ (%) | $C_{\mathrm{part.}}$ (%) | $N_{imp}$ |
|---|---|---|---|---|---|
| Munnink et al. [9] | SARS-CoV-2_83_LEFT | 70.0 | 0.000 | 0.000 | 0 |
| Munnink et al. [9] | SARS-CoV-2_83_RIGHT | 65.7 | 0.000 | 0.000 | 0 |
| Munnink et al. [9] | SARS-CoV-2_84_LEFT | 72.3 | 0.000 | 0.000 | 0 |
| Munnink et al. [9] | SARS-CoV-2_84_RIGHT | 64.6 | 0.000 | 0.000 | 0 |
| Munnink et al. [9] | SARS-CoV-2_85_LEFT | 69.8 | 0.000 | 0.000 | 0 |
| Munnink et al. [9] | SARS-CoV-2_85_RIGHT | 67.8 | 0.000 | 0.000 | 0 |
| Munnink et al. [9] | SARS-CoV-2_86_LEFT | 67.0 | 0.000 | 0.000 | 0 |
| Munnink et al. [9] | SARS-CoV-2_86_RIGHT | 68.3 | 0.000 | 0.000 | 0 |
| Munnink et al. [9] | SARS-CoV-2_87_LEFT | 69.1 | 0.000 | 0.000 | 0 |
| Munnink et al. [9] | SARS-CoV-2_87_RIGHT | 69.8 | 0.000 | 0.000 | 0 |
| Munnink et al. [9] | SARS-CoV-2_88_LEFT | 68.4 | 0.000 | 0.000 | 0 |
| Munnink et al. [9] | SARS-CoV-2_88_RIGHT | 68.5 | 0.000 | 0.000 | 0 |
| Munnink et al. [9] | SARS-CoV-2_89_LEFT | 66.9 | 0.000 | 0.000 | 0 |
| Munnink et al. [9] | SARS-CoV-2_89_RIGHT | 68.3 | 0.000 | 0.000 | 0 |
| Nalla et al. [10] | nCoV_2019-F | 66.3 | 0.000 | 0.000 | 0 |
| Nalla et al. [10] | nCoV_2019-P1 | 51.8 | 68.7 | 68.7 | 0 |
| Nalla et al. [10] | nCoV_2019-P2 | 54.8 | 16.9 | 16.9 | 0 |
| Nalla et al. [10] | nCoV_2019-R | 68.0 | 0.000 | 0.000 | 0 |
| Niu et al. [11] | E-F | 59.2 | 0.000 | 0.000 | 0 |
| Niu et al. [11] | E-P | 84.2 | 0.000 | 0.000 | 0 |
| Niu et al. [11] | E-R | 57.6 | 0.000 | 0.000 | 0 |

(continued on next page)

(Table S7, non-SARS coverage, continued)

| Set | Primer/probe | $T_{\text{ref.}}$ (°C) | $C_{\text{strict}}$ (%) | $C_{\text{part.}}$ (%) | $N_{imp}$ |
|---|---|---|---|---|---|
| Niu et al. [11] | RdRp-F | 64.6 | 0.000 | 0.000 | 0 |
| Niu et al. [11] | RdRp-P | 60.8 | 0.000 | 0.000 | 0 |
| Niu et al. [11] | RdRp-R | 68.4 | 0.000 | 0.000 | 0 |
| Park et al. [12] | SARS-CoV-2_IBS_E2-F | 65.2 | 0.000 | 0.000 | 0 |
| Park et al. [12] | SARS-CoV-2_IBS_E2-R | 61.3 | 0.000 | 0.000 | 0 |
| Park et al. [12] | SARS-CoV-2_IBS_N1-F | 63.3 | 0.000 | 0.000 | 0 |
| Park et al. [12] | SARS-CoV-2_IBS_N1-R | 63.9 | 0.000 | 0.000 | 0 |
| Park et al. [12] | SARS-CoV-2_IBS_RdRP2-F | 63.8 | 0.000 | 0.000 | 0 |
| Park et al. [12] | SARS-CoV-2_IBS_RdRP2-R | 64.5 | 0.000 | 0.000 | 0 |
| Park et al. [12] | SARS-CoV-2_IBS_S2-F | 63.7 | 0.000 | 0.000 | 0 |
| Park et al. [12] | SARS-CoV-2_IBS_S2-R | 61.6 | 0.000 | 0.000 | 0 |
| Park et al. [12] | SARS-CoV-2_IBS_m_N1-F | 60.8 | 0.000 | 0.000 | 0 |
| Park et al. [12] | SARS-CoV-2_IBS_m_N1-R | 65.4 | 0.000 | 0.000 | 0 |
| Park et al. [12] | SARS-CoV-2_IBS_m_N2-F | 60.3 | 0.000 | 0.000 | 0 |
| Park et al. [12] | SARS-CoV-2_IBS_m_N2-R | 62.4 | 0.000 | 0.000 | 0 |
| Park et al. [12] | SARS-CoV-2_IBS_m_RdRP1-F | 62.7 | 0.000 | 0.000 | 0 |
| Park et al. [12] | SARS-CoV-2_IBS_m_RdRP1-R | 60.8 | 0.000 | 0.000 | 0 |
| Park et al. [12] | SARS-CoV-2_IBS_m_RdRP2-F | 62.9 | 0.000 | 0.000 | 0 |
| Park et al. [12] | SARS-CoV-2_IBS_m_RdRP2-R | 62.2 | 0.000 | 0.000 | 0 |
| Park et al. [12] | SARS-CoV-2_IBS_m_S1-F | 62.5 | 0.000 | 0.000 | 0 |
| Park et al. [12] | SARS-CoV-2_IBS_m_S1-R | 59.3 | 0.000 | 0.000 | 0 |

(continued on next page)

(Table S7, non-SARS coverage, continued)

| Set | Primer/probe | $T_{\text{ref.}}$ (°C) | $C_{\text{strict}}$ (%) | $C_{\text{part.}}$ (%) | $N_{imp}$ |
|---|---|---|---|---|---|
| Park et al. [12] | SARS-CoV-2_IBS_m_S2-F | 61.4 | 0.000 | 0.000 | 0 |
| Park et al. [12] | SARS-CoV-2_IBS_m_S2-R | 60.3 | 0.000 | 0.000 | 0 |
| Rahman et al. [13] | M-F | 68.9 | 0.000 | 0.000 | 0 |
| Rahman et al. [13] | M-P | 70.9 | 0.000 | 0.000 | 0 |
| Rahman et al. [13] | M-R | 66.3 | 0.000 | 0.000 | 0 |
| Rahman et al. [13] | ORF1b-F | 65.5 | 0.000 | 0.000 | 0 |
| Rahman et al. [13] | ORF1b-P | 76.1 | 0.000 | 0.000 | 0 |
| Rahman et al. [13] | ORF1b-R | 64.2 | 0.000 | 0.000 | 0 |
| Toptan et al. [14] | M-475-F | 64.8 | 0.000 | 0.000 | 0 |
| Toptan et al. [14] | M-507-P | 63.5 | 0.000 | 0.000 | 0 |
| Toptan et al. [14] | M-574-R | 65.4 | 0.000 | 0.000 | 0 |
| Toptan et al. [14] | S-1850-F | 64.6 | 0.000 | 0.000 | 0 |
| Toptan et al. [14] | S-1884-P | 62.2 | 0.000 | 0.000 | 0 |
| Toptan et al. [14] | S-1949-R | 63.0 | 0.000 | 0.000 | 0 |
| Vogels et al. [15] | HKU-N-F | 58.6 | 0.000 | 0.000 | 0 |
| Vogels et al. [15] | HKU-N-P | 63.0 | 0.000 | 0.000 | 0 |
| Vogels et al. [15] | HKU-N-R | 60.3 | 0.000 | 0.000 | 0 |
| Vogels et al. [15] | HKU-ORF1-F1 | 61.7 | 0.000 | 0.000 | 0 |
| Vogels et al. [15] | HKU-ORF1-F2 | 64.6 | 0.000 | 0.000 | 0 |
| Vogels et al. [15] | HKU-ORF1-F3 | 58.7 | 0.000 | 0.000 | 0 |
| Vogels et al. [15] | HKU-ORF1-F4 | 61.7 | 0.000 | 0.000 | 0 |

(continued on next page)

(Table S7, non-SARS coverage, continued)

| Set | Primer/probe | $T_{\text{ref.}}$ (°C) | $C_{\text{strict}}$ (%) | $C_{\text{part.}}$ (%) | $N_{imp}$ |
|---|---|---|---|---|---|
| Vogels et al. [15] | HKU-ORF1-P1 | 64.5 | 0.000 | 0.000 | 0 |
| Vogels et al. [15] | HKU-ORF1-P2 | 63.6 | 0.000 | 0.000 | 0 |
| Vogels et al. [15] | HKU-ORF1-R1 | 61.7 | 0.000 | 0.000 | 0 |
| Vogels et al. [15] | HKU-ORF1-R2 | 65.5 | 0.000 | 0.000 | 0 |
| WHO | NIID_WH-1_F24381 | 61.2 | 0.000 | 0.000 | 0 |
| WHO | NIID_WH-1_F501 | 70.3 | 0.000 | 0.000 | 0 |
| WHO | NIID_WH-1_F509 | 63.3 | 0.000 | 0.000 | 0 |
| WHO | NIID_WH-1_R24873 | 61.5 | 0.000 | 0.000 | 0 |
| WHO | NIID_WH-1_R854 | 61.7 | 0.000 | 0.000 | 0 |
| WHO | NIID_WH-1_R913 | 69.2 | 0.000 | 0.000 | 0 |
| WHO | NIID_WH-1_Seq_F24383 | 60.4 | 0.000 | 0.000 | 0 |
| WHO | NIID_WH-1_Seq_F519 | 58.8 | 0.000 | 0.000 | 0 |
| WHO | NIID_WH-1_Seq_R24865 | 60.1 | 0.000 | 0.000 | 0 |
| WHO | NIID_WH-1_Seq_R840 | 60.2 | 0.000 | 0.000 | 0 |
| WHO | WH-NICN-F | 64.4 | 0.000 | 0.000 | 0 |
| WHO | WH-NICN-P | 51.3 | 0.000 | 0.000 | 0 |
| WHO | WH-NICN-R | 64.1 | 0.000 | 0.000 | 0 |
| WHO | WuhanCoV-spk1-f | 65.4 | 0.000 | 0.000 | 0 |
| WHO | WuhanCoV-spk2-r | 64.6 | 0.000 | 0.000 | 0 |
| WHO | nCoV_IP2-12669Fw | 54.3 | 0.000 | 0.000 | 0 |
| WHO | nCoV_IP2-12696bProbe | 67.0 | 0.000 | 0.000 | 0 |

(continued on next page)

(Table S7, non-SARS coverage, continued)

| Set | Primer/probe | $T_{\text{ref.}}$ (°C) | $C_{\text{strict}}$ (%) | $C_{\text{part.}}$ (%) | $N_{imp}$ |
|---|---|---|---|---|---|
| WHO | nCoV_IP2-12759Rv | 53.7 | 0.000 | 0.000 | 0 |
| WHO | nCoV_IP4-14059Fw | 54.8 | 0.000 | 0.000 | 0 |
| WHO | nCoV_IP4-14084Probe | 61.3 | 0.000 | 0.000 | 0 |
| WHO | nCoV_IP4-14146Rv | 54.8 | 0.000 | 0.000 | 0 |
| Yip et al. [16] | nsp2f | 63.4 | 0.000 | 0.000 | 0 |
| Yip et al. [16] | nsp2r | 61.4 | 0.000 | 0.000 | 0 |

[3] J. N. Davda, K. Frank, S. Prakash, G. Purohit, D. P. Vijayashankar, D. Vedagiri, K. B. Tallapaka, K. H. Harshan, A. B. Siva, R. K. Mishra, et al., An Inexpensive RT-PCR Endpoint Diagnostic Assay for SARS-CoV-2 Using Nested PCR: Direct Assessment of Detection Efficiency of RT-qPCR Tests and Suitability for Surveillance, bioRxiv .

[4] P. R. Grant, M. A. Turner, G. Y. Shin, E. Nastouli, L. J. Levett, Extraction-free COVID-19 (SARS-CoV-2) diagnosis by RT-PCR to increase capacity for national testing programmes during a pandemic, BioRxiv .

[5] Y. Hirotsu, H. Mochizuki, M. Omata, Double-Quencher Probes Improved the Detection Sensitivity of Severe Acute Respiratory Syndrome Coronavirus 2 (SARS-CoV-2) by One-Step RT-PCR, medRxiv .

[6] D. C. Lanza, J. P. Lima, S. M. Jerônimo, Design and in silico validation of polymerase chain reaction primers to detect severe acute respiratory syndrome coronavirus 2 (SARS-CoV-2) .

[7] C. Li, D. Debruyne, J. Spencer, V. Kapoor, L. Y. Liu, B. Zhang, L. Lee, R. Feigelman, G. Burdon, J. Liu, et al., High sensitivity detection of SARS-CoV-2 using multiplex PCR and a multiplex-PCR-based metagenomic method, bioRxiv .

[8] R. Lu, X. Zhao, J. Li, P. Niu, B. Yang, H. Wu, W. Wang, H. Song, B. Huang, N. Zhu, et al., Genomic characterisation and epidemiology of 2019 novel coronavirus: implications for virus origins and receptor binding, The Lancet 395 (10224) (2020) 565–574.

[9] B. B. O. Munnink, D. F. Nieuwenhuijse, M. Stein, A. O'Toole, M. Haverkarte, M. Mollers, S. K. Kamga, C. Schapendonk, P. Lexmond, M. Pronk, et al., Rapid SARS-CoV-2 whole genome sequencing for informed public health decision making in the Netherlands, bioRxiv .

[10] A. K. Nalla, A. M. Casto, M.-L. W. Huang, G. A. Perchetti, R. Sampoleo, L. Shrestha, Y. Wei, H. Zhu, K. R. Jerome, A. L. Greninger, Comparative performance of SARS-CoV-2 detection assays using seven different primer-probe sets and one assay kit, Journal of clinical microbiology 58 (6).
